# Timesweeper: Accurately Identifying Selective Sweeps Using Population Genomic Time Series

**DOI:** 10.1101/2022.07.06.499052

**Authors:** Logan S. Whitehouse, Daniel R. Schrider

## Abstract

Despite decades of research, identifying selective sweeps, the genomic footprints of positive selection, remains a core problem in population genetics. Of the myriad methods that have been developed to tackle this task, few are designed to leverage the potential of genomic time-series data. This is because in most population genetic studies of natural populations only a single period of time can be sampled. Recent advancements in sequencing technology, including improvements in extracting and sequencing ancient DNA, have made repeated samplings of a population possible, allowing for more direct analysis of recent evolutionary dynamics. Serial sampling of organisms with shorter generation times has also become more feasible due to improvements in the cost and throughput of sequencing. With these advances in mind, here we present Timesweeper, a fast and accurate convolutional neural network-based tool for identifying selective sweeps in data consisting of multiple genomic samplings of a population over time. Timesweeper population genomic time-series data by first simulating training data under a demographic model appropriate for the data of interest, training a one-dimensional Convolutional Neural Network on said simulations, and inferring which polymorphisms in this serialized dataset were the direct target of a completed or ongoing selective sweep. We show that Timesweeper is accurate under multiple simulated demographic and sampling scenarios, identifies selected variants with high resolution, and estimates selection coefficients more accurately than existing methods. In sum, we show that more accurate inferences about natural selection are possible when genomic time-series data are available; such data will continue to proliferate in coming years due to both the sequencing of ancient samples and repeated samplings of extant populations with faster generation times, as well as experimentally evolved populations where time-series data are often generated. Methodological advances such as Timesweeper thus have the potential to help resolve the controversy over the role of positive selection in the genome. We provide Timesweeper as a Python package for use by the community.

## INTRODUCTION

Over a century after the modern evolutionary synthesis, there is still a great deal of controversy over the role of natural selection in molecular evolution. In particular, the impact of positive selection, in which beneficial alleles are favored by selection and thus increase in frequency in the population, is hotly debated (Hahn 2008; Kern and Hahn 2018; Jensen *et al*. 2019). This controversy extends to the role of selective sweeps (Stephan 2010), in which positively selected mutations rapidly sweep through a population thereby drastically reducing and altering patterns of genetic diversity in the vicinity of the selected locus (Smith and Haigh 1974; Kaplan *et al*. 1989). with some studies claiming that signatures of selective sweeps alter genome-wide patterns of polymorphism (Enard *et al*. 2014; Garud *et al*. 2015; Garud and Petrov 2016; Schrider and Kern 2017; Booker *et al*. 2021), and others purporting that these signatures may be largely or even entirely false positives (Harris *et al*. 2018).

A number of methods have been devised to make inferences about positive selection in the recent past, including recently completed or even ongoing selective sweeps, on the basis of snapshots of present-day variation obtained from a set of genomes sampled from the population(s) of interest. These methods typically examine one of several aspects of genetic diversity that are each indicative of an allele that has spread rapidly enough such that there has been little time for recombination and mutation to introduce diversity into the class of haplotypes carrying the sweeping allele. These signatures of selection include: characteristic skews in the frequencies of neutral alleles linked to the selected site (Fay and Wu 2000; Kim and Stephan 2002; Li 2011a), reduced haplotypic diversity around the selected site (Hudson *et al*. 1994; Garud *et al*. 2015; Harris and DeGiorgio 2020), the presence of abnormally long-range haplotypes bearing the selected allele (Sabeti *et al*. 2002; Voight *et al*. 2006; Ferrer-Admetlla *et al*. 2014), increased linkage disequilibrium (Kelly 1997), especially on either flank of the selected site (Kim and Nielsen 2004), and elevated divergence between closely related populations in the case of local adaptation (Chen *et al*. 2010).

One pernicious obstacle to efforts to accurately detect sweeps is demographic change. For example, strong population bottlenecks, which involve a recovery from a period of drastically reduced population size, can result in rapid changes in allele frequencies that can produce false-positive signals of selective sweeps, especially if the demographic history is unknown (Jensen *et al*. 2005; Nielsen *et al*. 2005; Mughal and DeGiorgio 2019). Some recent methods combine each of the signatures of selection described above via machine learning in order to identify the multidimensional spatial patterns of diversity along the chromosome that sweeps produce (Lin *et al*. 2011; Pybus *et al*. 2015; Sugden *et al*. 2018; Mughal and DeGiorgio 2019; Xue *et al*. 2021; Caldas *et al*. 2022; Lauterbur *et al*. 2022), have proved to be far more robust to demographic events (Schrider and Kern 2016; Mughal and DeGiorgio 2019). However, even these methods may experience an appreciable loss of power under non-equilibrium demographic histories, especially if they are not adequately modeled during the training process (Schrider and Kern 2016; Mughal and DeGiorgio 2019). Thus, in spite of decades of theoretical and empirical research, detecting selective sweeps remains a major challenge.

The difficulty in discriminating between positive selection and purely neutral processes could potentially be alleviated by using population genomic time-series data, in which genomes are sampled across a range of timepoints rather than a single contemporaneous point. Such data allow us to more directly track the trajectories of any potentially sweeping alleles/haplotypes, and to determine if they appear to be spreading faster than other alleles segregating in the population. Population genomic time-series data are becoming more and more prevalent, in part because it is often feasible, and increasingly affordable, to collect and sequence longitudinal samples of populations from species with rapid generation times (Pennings *et al*. 2014; Bergland *et al*. 2014; Feder *et al*. 2021; Machado *et al*. 2021). For example, studies conducted in collections repeated each season in *Drosophila* over a period of several years have presented evidence that seasonal oscillations in the frequencies of some alleles may be adaptive <u>(Bergland *et al*. 2014; but see Buffalo and Coop 2020)</u>, and that temporal variances in allele frequencies are explained in part by positive selection (Bertram 2021). Even in organisms with longer generation times, the production of time-series data may be possible in some cases. For example, a large number of ancient and archaic human genomes have been sequenced, and this has allowed researchers to directly ask whether alleles proposed to be under positive selection have indeed rapidly increased in frequency, and to more precisely infer when these frequency changes occurred (Hummel *et al*. 2005; Bollback *et al*. 2008; Wilde *et al*. 2014; Sverrisdóttir *et al*. 2014; Enard *et al*. 2014; Malaspinas 2016; Jeong Choongwon *et al*. 2016; Olalde *et al*. 2019).

As population genomic time-series continue to proliferate, novel methods that can fully take advantage of the potential of such data are required. One area where such methods have been in development for a number of years is experimental evolve-and-resequence (E&R) studies (reviewed in Schlötterer *et al*. 2015), in which lab populations are subjected to controlled selective pressures and often sampled during the course of the experiment (e.g. Illingworth *et al*. 2012; Feder *et al*. 2014; Terhorst *et al*. 2015; Otte and Schlötterer 2021). These methods include statistical tests to distinguish between selected an unselected loci on the basis of allele frequency changes through the course of the experiment, methods that use hidden Markov models to obtain maximum likelihood estimates of parameters such as the selection coefficient (Mathieson and McVean 2013; Steinrücken *et al*. 2014; Iranmehr *et al*. 2017), and Bayesian methods for parameter estimation (Schraiber *et al*. 2016; Ferrer-Admetlla *et al*. 2016). Some of these methods take advantage of the fact that E&R studies often produce experimental replicates, allowing for more confident identification of selected loci in cases where positive selection has acted on these loci in more than one replicate (Vlachos *et al*. 2019). However, in natural populations, controlled replicates are unavailable, and we therefore require methods that can accurately detect targets of selection in a single population genomic time-series.

Here, we describe a deep learning method for detecting positive selection from population genomic time series. This method, called Timesweeper, constructs an image representation of temporal changes in allele or haplotype frequencies around a focal site and then, with the aid of a convolutional neural network (CNN), uses these data to 1) classify the site as either experiencing a recent selective sweep or not, and 2) estimate the selection coefficient of a candidate sweeping mutation. We show that this method can localize sweeps and infer selection coefficients with better accuracy than competing methods. This method is trained on simulated data under a user-specified demographic model and sampling scheme, but we show that it can be robust to at least some scenarios of demographic model misspecification during training. Moreover, because the default mode of Timesweeper examines only changes in sample allele frequencies over time, it is applicable to data for which phased haplotypes or even high-confidence genotypes are not available, including pooled population genomic sequence data or lower coverage genomes such as ancient human DNA. We demonstrate this practical utility on an E&R dataset of 10 experimental replicate population of *D. simulans* (Barghi *et al*. 2019), finding that we are able to identify selective sweep signatures and replicate candidate sweep loci at a high rate. Timesweeper is also computationally efficient, completing the training step within several minutes even without the use of GPUs. We argue that methodologies such as Timesweeper, which seek to better leverage population genomic time-series to detect natural selection, have the potential to not only uncover targets of positive selection with unprecedented accuracy, but to help resolve the lingering controversy over the role of positive selection in shaping patterns of diversity within species.

## METHODS

### Overview of the Timesweeper workflow

Timesweeper is a Python package consisting of multiple interconnected modules that can be used in sequence to simulate training and test data consisting of genomic regions with and without selective sweeps given a user-specified demographic model and sampling scheme (i.e. the time and size of each sample in the series), process the resulting simulation output and convert it into useable data formats, use these data to train neural networks to detect selective sweeps and infer the selection coefficient (*s*), and finally to perform inference on real data using the trained network. Each of these steps is described in detail below, with full command line examples and options described in the software’s readme found at https://github.com/SchriderLab/Timesweeper/blob/master/README.md. Timesweeper is developed as separate modules as opposed to a single workflow with modularity and customizability in mind—the code is intended to be used out of the box but can also be adapted to a user’s needs by modifying or replacing any of the constituent modules if desired.

Timesweeper can detect sweeps using one of two pieces of information: allele frequency trajectories surrounding a focal polymorphism, and haplotype frequency trajectories within a focal window. All modules in Timesweeper can be run using a single YAML configuration file across the workflow. Most modules are optimized for multiprocessing and allow for specification of the number of threads using the --threads flag at runtime. Timesweeper’s simulation modules allow for the partitioning of replicate ranges, allowing for straightforward parallelization across a high-performance computing system during what is the most computational-heavy stage of the workflow.

### Simulating using custom SLiM scripts

Timesweeper has two modules for simulating training data using SLiM (Haller and Messer 2019 p. 3): a module for simulating from custom-made SLiM scripts and another for using pre- made stdpopsim (Adrion *et al*. 2020a) demographic models. The former is relatively straightforward: a user supplies a working directory, the path to their SLiM script, the path to the SLiM executable, and the number of replicate simulations they’d like to produce. Other optional parameters supplied at runtime are passed to SLiM through the simulate_custom module of Timesweeper. This module is intended to be modified to suit the needs of the experiment, and is written such that it is easy to incorporate utility functions, stochastic variable draws, and more into each replicate.

Users must write their SLiM script such that it accepts two constant parameters at runtime: the sweep type, and the output file name. The sweep types are denoted by the “scenarios” list in the YAML configuration file, and a user may write their SLiM script to use these arguments however they wish. In our default workflow, they specify the class of training/test data to be generated, with “neut” denoting simulations lacking a selective sweep, “sdn” denoting simulations with a selective sweep on a single-origin *de novo* mutation (hereafter referred to as the SDN model), and “ssv” denoting simulations with a selective sweep from standing variation (the SSV model). An example SLiM script is included in the software package, and many more included with the GitHub repository associated with the experiments in this manuscript. Note that in all of our simulations of the SDN and SSV models, we condition on the selected allele not being lost— if the allele is lost during a simulation run, the simulation then jumps back to the point at which the beneficial mutation is introduced in the center of the simulated chromosome (SDN) or the neutral allele nearest to the center is chosen to be given a positive selection coefficient (SSV).

Note that SSV sweeps with lower starting allele frequencies are more likely to be lost, and thus if we were to condition on a randomly selected polymorphism to go to fixation, we would obtain a downwardly biased initial selected frequency relative to what would be expected under an SSV model that correctly deals with different fixation probabilities (see (Hermisson and Pennings 2017)). To mitigate this problem in our simulations, when an SSV sweep is lost, we restart the simulation from 500 generations prior to the onset of selection so that there is the opportunity for different mutations to be selected in subsequent attempts. The rationale for this design choice is that those sweep attempts that happen to select a more common polymorphism will be more likely to result in fixation, although we did not investigate the extent to which this avoids the frequency bias described above. We also note that our SSV mutations do not condition on a soft sweep—it is possible that SSV sweeps may involve only a single ancestral copy of the adaptive allele that reaches fixation. Users writing their own custom scripts are free to adjust these conditions or indeed any other aspect of Timesweeper’s three classes however they wish.

The amount of time required to run simulations varies greatly by the complexity of the demographic model and the parameterization. For all of the constant population size simulations in this manuscript the average runtime of a single replicate was 10 seconds for the Neutral and SSV and scenarios and 25 seconds for the SDN scenario, which is in concordance with the average number of restarts for each class being 0 for Neutral (as no restarts are needed), 0.5 for SSV, and 15 for SDN simulations, respectively. No conditioning on allele frequency upon sweep start nor upon sampling finishing was implemented for any experiments described. The average runtimes given here were obtained on a cluster where the typical compute node has two Model E5-2680 v3 2.50 GHz Intel Xeon processors (24 cores in total, although each simulation used only a single core), 30M cache, and 256-GB RAM.

### Simulating using stdpopsim

The second option for simulating training data is by injecting time-series sampling code into a SLiM script generated by the stdpopsim package (Adrion *et al*. 2020a). By using stdpopsim with the slim and --slim-script flags, stdpopsim will write out a SLiM script for running whichever demographic model the user selects from the stdpopsim catalog. This script can then be used as input to Timesweeper’s simulate_stdpopsim module, along with information about sampling times (specified in years as per stdpopsim’s convention), sampling sizes, the identity of the population that is sampled and experiences any positive selection (hereafter referred to as the “target population”), and the range of selection coefficients to use in a log-uniform distribution. This module then adds the desired events to the stdpopsim SLiM script before running the simulation. The output of this module is identical in format to that generated when using the simulate_custom module described above, and is processed identically downstream. We note that although stdpopsim supports selective sweeps with some degree of customizable parameters, this feature does not currently support all of the scenarios we desired for our workflow (e.g. selection on a randomly chosen polymorphism that was previously fitness-neutral, while specifying only the time at which the derived allele later became beneficial). As more features are added to the stdpopsim software, we may update our workflow to better take advantage of this resource.

### VCF file processing post-simulation

For both simulation modules described above, at each sampling timepoint, SLiM draws a random sample of the specified number of individuals from the target population without replacement, and concatenates a VCF entry to an output file, hereafter referred to as a “multiVCF”, for that replicate. After a simulation replicate is complete, the multiVCF is split into separate files consisting of one timepoint each, labeled in numerical order according to when they were sampled in the series. Note that because our workflow uses forward simulations, the sweeping mutation may often be lost, requiring the simulation to jump back to an earlier point. This may result in repeated samplings drawn for one or more sampled timepoints, but our workflow ensures that only the last sample for each timepoint (corresponding to the simulation run where the sweeping mutation was not lost) is retained. VCF files are then sorted and indexed using BCFtools (Li 2011b; Danecek *et al*. 2021) sort and index. Individual VCFs are then merged using the command bcftools merge --force-samples -0 to create a VCF containing all polymorphisms among the set of all genomes sampled across all timepoints. Information from these VCFs can then be loaded into a format suitable for input to our neural network as described in the next two sections.

### Time-series allele frequency matrices

Timesweeper’s allele frequency-tracking (AFT) method seeks to classify each individual polymorphism on the basis of allele frequencies at each of *l* biallelic polymorphisms in a surrounding window that consists of the central polymorphism and (*l* − 1)/2 polymorphisms on either side, measured at *m* distinct timepoints. We therefore chose to represent allele frequencies around a focal polymorphism by an *l* × *m* matrix, which serves as our input for both training and prediction. This was constructed by first obtaining sample allele frequency trajectories from merged VCF files. For a single merged VCF, whether obtained from either real or simulated data, genotypes are loaded using scikit-allel (Miles *et al*. 2021), and the two alleles at each polymorphism are labeled as “net-increasing” or “net-decreasing” according to their change in frequency between the first and final timepoint. For polymorphisms with no net frequency change, the “net-increasing” or “net-decreasing” are randomly assigned to the two alleles. The desired *l* × *m* matrix showing the frequency of the net-increasing allele for each polymorphism at each timepoint is then created (shown for several individual replicates as well as mean values in Figure 1A and Figure S1). For this paper, we set *l* to 51 (i.e. 25 flanking polymorphisms are examined around each focal polymorphism to be classified) unless otherwise noted. When dealing with simulated data, the number of polymorphisms obtained was always greater than *l*, and thus the matrix was cropped to obtain *l* polymorphisms as described below in “Classifier Training and Validation.” When applying Timesweeper to real (or test) data, a sliding window of *l* polymorphisms can be used to obtain a classification for each individual polymorphism (see “Detecting Sweeps in Genomics Windows” below). Given that some users may wish to examine a larger window, we give users the option of changing the value of *l* when running Timesweeper.

**Figure 1.**
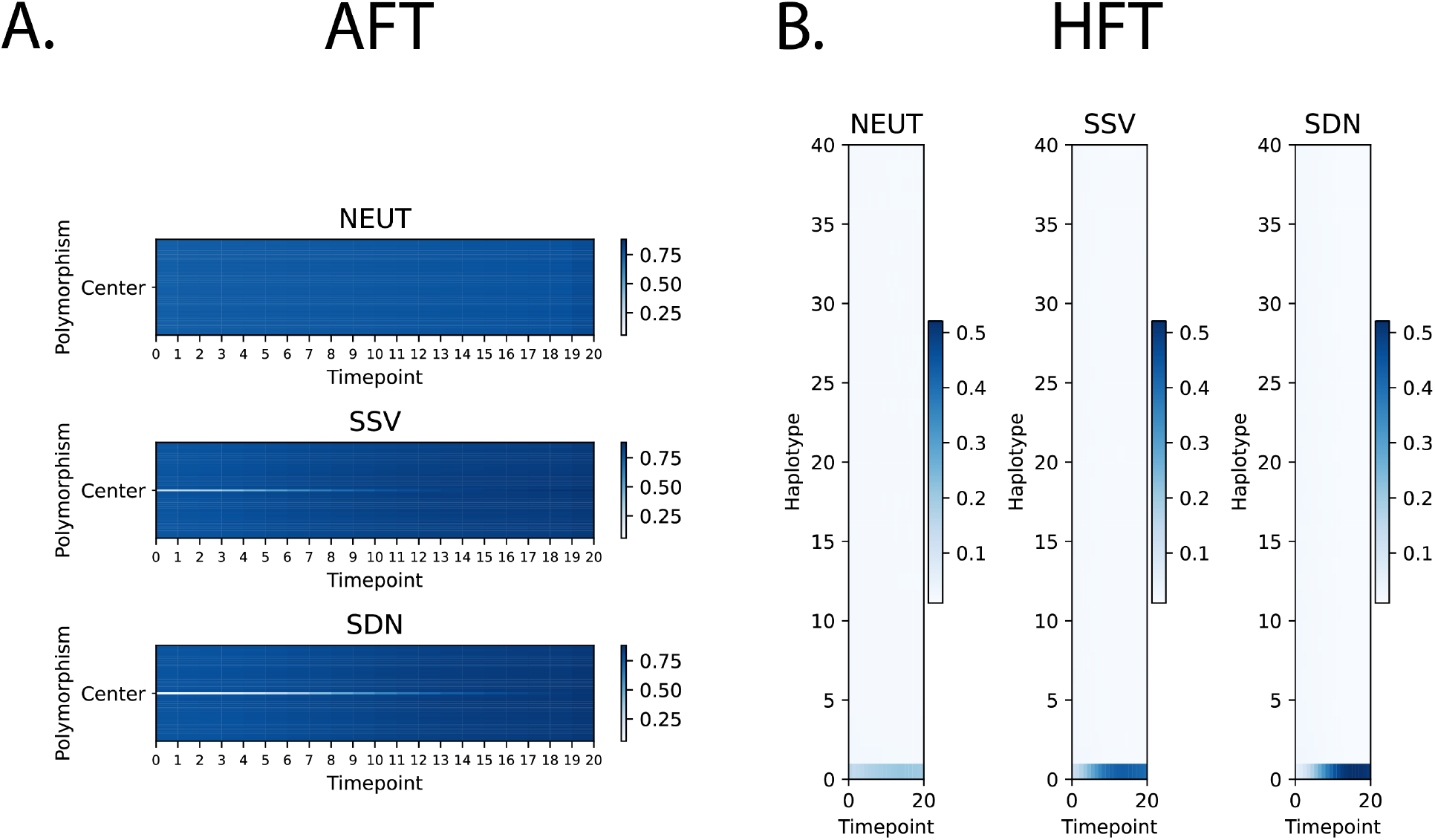
Timesweeper input formats. (A) Allele Frequency Tracker (AFT) data format: mean values of allele frequencies of the allele with the largest increase in frequency from the first to final timepoint in a 51-polymorphism window over 20 timepoints. (B) Haplotype Frequency Tracker (HFT) data format: mean values of haplotype frequencies for the haplotype with the largest frequency increase from the first to last timepoints (bottom row) and the 39 most-similar, ranked by decreasing similarity to the bottom haplotype. For both formats, the mean input images were calculated from 10,000 individual replicates for each class. For the SDN and SSV classes, selection coefficients were drawn from a uniform distribution with bounds [0.00025, 0.25] and with a starting sampling generation drawn from a uniform distribution with bounds [-50, 50] generations from the onset of selection.

### Time-series haplotype frequency matrices

Timesweeper’s haplotype frequency-tracking method constructs matrices containing information about haplotype frequency trajectories. For a given window of *l* polymorphisms, the frequencies of all haplotypes are recorded at each timepoint using information from the merged VCF (again processed using scikit-allel), and recorded in a *k* × *m* matrix, where *m* is again the number of timepoints and *k* is the number of distinct haplotypes observed across all samples (shown for individual replicates as well as mean values in Figure 1B and Figure S1B). This matrix is then sorted such that the haplotype with the highest net increase in frequency between the first and last timepoints is located at the “bottom” of the frequency matrix, i.e. the 0^th^ index. The remaining haplotypes are then sorted by their similarity (Hamming distance) to the haplotype at the 0^th^ index (i.e. the haplotype at index 1 is the most similar to the haplotype with the highest net increase in frequency, the haplotype at index 2 is the second-most similar, and so on). In the context of using CNNs to perform inference on population genetic alignments, it has been noted that sorting haplotypes based on similarity increases accuracy for multiple tasks (Flagel *et al*. 2019; Ray *et al*. 2023), because such sorting makes informative structures (e.g. similarities between populations when searching for evidence of gene flow) more readily visible. Our rationale was that sorting haplotypes by their frequency clearly distinguishes selected from neutrally evolving regions (e.g. Figure 3 from (Garud *et al*. 2015), and this appears to carry over into 2-dimensional visualizations of haplotype frequency time series (Figure S1B). However, we did not test the impact of this sorting on accuracy, and we note that for the one-dimensional CNN used for most of our analyses, the order of haplotypes chosen would not impact performance so long as the manner in which haplotypes are ranked is consistent between training and test sets. For two-dimensional CNNs, the order could potentially impact performance even if it is held consistent across training and testing.

### Neural network architectures

We implemented all neural network models in Keras (Chollet and others 2015) with Tensorflow backends, including: a shallow 1D convolutional neural network (1DCNN) described below; a similar network but with shallow 2D rather than 1D convolutional layers (2DCNN);a Recurrent Neural Network (RNN) model for time-series data; and, for single-timepoint data, series data and a shallow fully-connected network (FCN) consisting of several fully connected layers each followed by dropout. Architectures for both AFT and HFT data types (allele frequencies and haplotype frequencies) are identical in structure, although the number of parameters differ due to the difference in input image size aside from the adjustment in input layer dimensions.

The 1DCNN time-series network begins with two 1D convolutional layers with 64 filters and a kernel size of 3 each (i.e. the first convolutional layer runs rectangular filters across an area of all polymorphisms/haplotypes by 3 timepoints at a time, sliding across timepoints with a stride length of 1. These are followed by a 1D maximum pooling layer with a pool size of 3, followed by a dropout layer with a dropout rate of 0.15. A flattening layer then precedes two blocks of 264- unit dense layers followed by dropout layers with a rate of 0.2. This is followed by a 128-unit dense layer, a dropout layer with dropout rate of 0.3, and a final dense layer with 3 output units. All layers described use ReLu activation function other than the final output layer, which uses a softmax activation function for classification (identifying and classifying sweeps), and linear activation for regression (inferring *s*). Our rationale for having the 1D convolutional filters stride across timepoints was that they would be able to examine the selected allele and any linked hitchhiking alleles in any given timepoint, and track them together across timepoints. We did not experiment with transposing the input such that filters stride across polymorphisms rather than time.

The one-timepoint network is a fully connected network (FCN) consisting of two blocks of 512-unit dense layers followed by dropout layers with rates of 0.2, followed by a 128-unit dense layer with dropout of 0.1, and a final dense layer with 3 output units. This network also uses ReLu activations for all layers aside from the final output layer, which uses a softmax activation function for classification and linear activation for regression.

The full architectures for the 1DCNN, single-timepoint FCN, and other benchmarked architectures including a 2DCNN, a more complex 1DCNN with more trainable parameters, and a recursive neural network (RNN) are shown in Figure S2.

### Network training and validation

To create training labeled data for models we first process simulated VCFs as discussed above. During this process the mutation type specified in the SLiM simulation is obtained from each VCF entry, allowing us to retrieve the selected mutation (the only non-neutral mutation present in the simulation). When constructing training sets for the SDN and SSV scenarios for the AFT method, by default the selected mutation is retrieved and set to be the center of the 51-polymorphism window (i.e. only the 25 closest polymorphisms on either side of the selected mutation are retained). For the neutral scenario the central-most polymorphism is set to be the center of the 51- polymorphism window. For the HFT method, the data are formatted as described above without regard to the selected mutation (i.e. the haplotype found at the highest frequency is placed at the bottom of the matrix, regardless of whether or not it contains a selected mutation). Windows of the designated size are saved in a compressed pickle file and loaded into memory at training time along with the simulation replicate’s selection coefficient and sweep type (if applicable), and a replicate identifier.

Data is split into training/validation/testing partitions consisting of 70%/15%/15% of the entire dataset, respectively, and each partition is stratified such that classes are evenly split among them. Models are trained for a maximum of 100 epochs with the potential for early stopping based on validation accuracy: training stops if at any point 10 consecutive training epochs fail to improve upon the highest validation accuracy (classification) or validation mean-squared error (regression) achieved up to that point. The validation partition is used for hyperparameter tuning of the ADAM optimizer (Kingma and Ba 2017), while the test partition is used exclusively evaluating performance post-training. If early stopping occurs model weights are restored to the epoch with the highest validation accuracy, otherwise the full number of epochs is trained and then the weights with the highest validation accuracy are saved using Keras’ `ModelCheckpoint` functionality.

### Detecting sweeps and inferring selection coefficients in genomic windows

Timesweeper’s final module, cleverly named find_sweeps, reports AFT classifier probabilities and estimated selection coefficients and optionally HFT classifier probabilities and selection coefficients for each sliding window of polymorphisms across all polymorphisms in a given VCF, whether obtained from simulated data (for testing) or real data (for inference). Trained neural networks are first loaded into memory and the input VCF is loaded in chunks. For each polymorphism in a chunk (except for those within (*l*-1)/2 bp of the edge of a chromosome), data is processed into the required formats described above for each method prior to classification/inference. The find_sweeps module also allows for testing on simulated data using the --benchmark flag, which causes the module to check each prediction against the ground truth from the simulation, allowing for easy validation of all methods on labeled test data.

### Simulations for testing Timesweeper on time-series data from a constant-sized population

As in the standard Timesweeper workflows described above, all simulations used to evaluate Timesweeper’s accuracy were performed via SLiM (Haller and Messer 2019). For the constant-sized population scenarios a population of 500 individuals was created with a mutation rate of 1×10^-7^ and recombination rate of 1×10^-7^. Simulations underwent burn-in for 20*N* generations. In the SDN scenario a *de novo* mutation is introduced at the physical midpoint of the chromosome with a given selection coefficient. In the SSV scenario the closest standing neutral mutation to the center of the chromosome is selected and the selection coefficient of that mutation is set to a value from a uniform distribution with bounds [0.00025, 0.25) with the exception of the selection coefficient experiments where *s* was set at a constant value for each experiment. Sampling began at a time, measured in generations after the onset of selection, sampled from a uniform distribution with bounds [-50, 50) except when noted in the selection timing experiments. A number of sampling schemes and sweep parameterizations were tested under this constant-size scenario, as described in the Results. For each of these experiments 10,000 replicates of each scenario were simulated: sweeps under the SDN model, SSV sweeps, and neutrality. For experiments testing the impact of the number of replicates, data were subsampled from a total pool of 30,000 replicates for each class to create a dataset of the desired size. Data were then partitioned into training, testing, and validation sets as described above.

### Testing the effects of demographic model misspecification on Timesweeper’s accuracy

We simulated three demographic models of increasing complexity: a constant-sized population (described above), herein referred to as the “simple model”, a three-epoch bottleneck model fit to a dataset of Utah CEPH individuals by (Marth *et al*. 2004), herein the “bottleneck model”, as well as the Out-of-Africa model described in (Gutenkunst *et al*. 2009) as implemented in stdpopsim, herein “OoA model”. For the OoA model we sampled only individuals from the population modeled after the Han Chinese (CHB). For all models we sampled 20 evenly spaced timepoints with 10 diploid individuals at each timepoint. Sampling started at a generation drawn from a uniform distribution between -50 and 50 generations post-selection and proceeded for a total of 200 generations. For the SSV and SDN scenarios of each model, selection coefficients were randomly drawn from a uniform distribution with bounds [0.00025, 0.25). We simulated 10,000 replicates of the neutral, SDN, and SSV scenarios under all models and trained Timesweeper as described above. We then used each trained CNN to detect sweeps on 5,000 additional independent test replicates from each demographic model. Performance metrics were then calculated on the class probabilities and selection coefficient estimate predictions across all replicates.

### Previous methods assessed in this study

We compared the performance of Timesweeper to a number of previously published methods for detecting selection or inferring selection coefficients from genomic time-series, and we briefly describe these below. Each method’s performance was evaluated on 5,000 test replicates of each class, simulated with identical parameterization as described above, with selection coefficients drawn from a uniform distribution with bounds [0.00025, 0.25) and the initial sampling generation again sampled from a uniform distribution with bounds [-50, 50).

### Binary classification using the Frequency Increment Test

The Frequency Increment Test (FIT) detects selection in time-series data by determining whether the allele frequency increments from timepoint to timepoint significantly differ from zero. We implemented FIT in Python following Feder et. al. (2014). In brief, SLiM output was parsed identically to the method described above for the time-series allele frequency spectrum, and the rescaled allele frequency increments for each polymorphism in the simulated region are calculated as 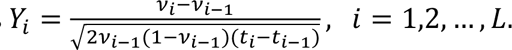 where *v*_*i*_ and *t*_*i*_refer to the allele frequency and time at timepoint *i* of *L*, respectively. The frequency increments *Y*_*i*_ are used in a one-sample student’s *t*-test for each polymorphism. We used a *p*-value threshold of 0.05 to classify a polymorphism as either experiencing a sweep or evolving neutrally. No effort is made to distinguish between sweep types using this test.

### Fisher’s exact test

Fisher’s exact test can be used to detect a significant difference in allele frequencies between timepoints of a longitudinal dataset, and we used it to test for frequency differences between the first and last timepoints (following (Barghi *et al*. 2019)). We used the SciPy (Virtanen *et al*. 2020) implementation of the Fisher’s exact test, with its input being a contingency table consisting of the and major and minor allele counts for the first and final samples.

#### Slattice

slattice (Mathieson and McVean 2013) is a Hidden Markov-Model (HMM) approach originally developed for inferring selection coefficients from spatially structured populations using time-series data. For input it requires sample sizes and allele counts from time- series data for every generation during the sampling window. We ran slattice in a similar manner as described in the software’s vignettes (see https://github.com/SchriderLab/timesweeper-experiments/blob/main/scripts/comp_methods/vcf_to_slattice.py for our commands). VCFs were parsed using scikit-allel and focal loci (the centermost polymorphism in the case of neutral scenarios, and the true target of selection for the SDN or SSV scenarios) were formatted and written to a temporary input file that was then used to run slattice with the default options.

#### ApproxWF

ApproxWF (Ferrer-Admetlla *et al*. 2016) is another HHM-based approach that uses an approximated Wright-Fisher process, resulting in faster runtimes than other HMM approaches to estimate selection coefficients from time-series data. Similar to the manner in which we ran slattice, VCFs were parsed, allele counts and sampling sizes were converted into temporary files in concordance with the format required to use ApproxWF, which we then ran using default parameters.

#### WFABC

WFABC (Foll *et al*. 2015) is an approximate Bayesian computation (ABC)-based method used to infer effective population size and selection coefficients using time-series data. WFABC was run with default parameters using input formatted from VCF files similarly as described above.

#### Sweepfinder2

Sweepfinder2 (DeGiorgio *et al*. 2016) is an updated version of the popular Sweepfinder (Nielsen *et al*. 2005) tool, which examines the allele frequencies at polymorphisms at various genetic distances from a focal site to determine if it was the target of a recently completed selective sweep. SweepFinder accomplishes this by using a composite likelihood ratio test comparing the probabilities of these allele frequencies under a hitchhiking model to the probabilities under neutrality (i.e. the background site frequency spectrum). Sweepfinder2 was run on VCFs containing only the final timepoint of each replicate with a grid size of 1kb (with only the central point in this grid examined to calculate entries of the confusion matrix), after which the 95% composite likelihood ratio (CLR) threshold was calculated as the 95^th^ percentile of the neutral CLRs across all windows. Regions with reported a CLR value above this threshold were labeled as a predicted sweep, otherwise the region was labeled as neutral.

#### diploSHIC

diploSHIC (Kern and Schrider 2018) is a version of S/HIC (Schrider and Kern 2016) that has been updated to be applicable to unphased diploid as well as phased haploid data. diploSHIC uses a neural-network approach for sweep detection that utilizes a variety of summary statistics calculated in a number of sub-windows surrounding (and including) a focal window to be classified as a sweep (hard or soft), closely linked to a sweep (hard or soft), or as neutrally evolving. In diploSHIC, these summary statistics are arranged into a two-dimensional feature array used as input to a CNN. The dimensions of this array are *f* × *l*, where *f* the number of features (12) and *l* is the number of subwindows (a default value of 11, which we used here). We converted VCFs of the final timepoint of each replicate into diploSHIC’s input images for each replicate in the training and testing data. The diploSHIC CNN was trained for 10 epochs and then evaluated by running predictions on the central window of held out testing data.

#### TS-SHIC

We developed an extended version of diploSHIC that examines time-series data, which we refer to as TS-SHIC. TS-SHIC first calculates the *f* × *l* feature input array as described above for each timepoint, and then concatenating them into an *f* × *t* format, where *t* is the number of timepoints, that was then used to train and test the standard 1DCNN architecture described above. All training and testing was performed identically to other Timesweeper benchmarks, including 10,000 replicates for each class for training, 5,000 replicates of each class for testing.

### Applying Timesweeper to a *Drosophila simulans* evolve-and-resequence dataset

To demonstrate Timesweeper’s flexibility and applicability to real data, we applied the AFT method to a dataset published by Barghi *et al*. (2019) comprised of time-series *Drosophila simulans* pool-sequencing data from an evolve-and-resequence study. In brief, Barghi et al. exposed 10 replicates of an ancestral population to extreme temperatures in 12-hour cycles for 60 generations. Each experimental replicate population was sampled every 10 generations, yielding a total of 7 sampled timepoints including the initial ancestral population. Barghi *et al*. (2019) then performed pooled sequencing, read mapping, and genotyping on each replicate population for each of these timepoints. We downloaded the read counts for each allele at each SNP at each timepoint from https://doi.org/10.5061/dryad.rr137kn (the relevant .sync file is in the F0-F60SNP_CMH_FET_blockID.sync.zip archive, along with its associated README). The data for each timepoint were converted into allele frequency estimates by simply taking, for each allele, the number of reads supporting that allele divided by the total number of reads mapped to that site. Net alleles frequency changes were then calculated by taking the allele with the largest net frequency increase over the course of the experiment, and the allele with the largest frequency of all the remaining alleles (i.e. only two alleles were considered at each site), and renormalizing so that these two allele frequency estimates summed to one for each timepoint. This process was repeated for each experimental replicate.

The training data for this analysis were simulated using a modified simulate_custom module of Timesweeper to allow for population-size rescaling by a factor of 100 during the burn-in, with recombination and mutation rates multiplied by this factor to compensate, followed by the remainder of the simulation (i.e. the 60 generations of selection) being carried out without rescaling. The purpose of rescaling during the burn-in only was that the population size was small during the experiment (*N*=1000), but most likely dramatically larger in the ancestral population. The unscaled recombination rate was drawn from a uniform distribution between 0 and 2×10^-8^ and the unscaled mutation rate was set to 5×10^-9^, with only neutral mutations occurring. The simulation began with a burn-in period of 20 times the scaled population size (*N*=2,500, corresponding to an unscaled size of 250,000), for 50,000 generations. We note that this ancestral population size is a rough estimate that we obtained by calculating nucleotide diversity within each replicate’s founder sample (≈0.005), and is likely smaller than the *D. simulans* population used to found the experimental lines as it does not account for the impact of repetitive/low quality regions or natural selection on diversity, resulting in simulations with a substantially lower density of polymorphism than found in the real data (an average of 0.0065 polymorphisms per base pair in our simulations, vs. 0.45 for chromosome 2L in the empirical data). As we show in the Results, Timesweeper performed well on this dataset in spite of this difference between the training and empirical data. Following the burn-in, the population then contracted to 1,000 individuals, where it remained for the rest of the simulation, and a sample was taken to represent the starting “ancestral” state of the replicate, following the design used by Barghi *et al*. (2019).

In simulations with selection, the mutation closest to the physical center of the chromosome was then changed from neutral to beneficial with a selection coefficient drawn from a log-uniform distribution with a lower bound of *s*=0.02 and an upper bound of *s*=0.2—note that only SSV sweeps were generated under this model, as the SDN model is unlikely to be relevant in such a small population under selection for such a short period of time. The simulations then proceeded for another 60 generations post-selection with sampling occurring every 10 generations, resulting in a total of 7 sampled timepoints as in in Barghi *et al*. (2019). For each of these 7 timepoints in each replicate population, we sampled 100 diploid individuals. This was performed for a total of 7,500 replicate simulations for both the neutral and sweep SSV conditions that were then processed and used to train a Timesweeper network using the AFT method as described above, which was then used to detect sweeps in the *D. simulans* data using the find_sweeps module.

## RESULTS

### Representations of population genomic time-series data

We devised two representations of genomic time-series data with the goal of capturing information that would be informative about the presence of selective sweeps occurring during the sampling period. The first of these is an *l* × *m* matrix, where *l* is the number of polymorphisms and *m* is the number of sampling timepoints. The motivation for this format is that the frequency trajectories of all alleles, including that of a focal allele at the center of the window, can be tracked through time, along with spatial information along the chromosome. Not only should the focal allele increase in frequency if it is positively selected, but some nearby alleles may also hitchhike to higher frequencies, with more distant polymorphisms experiencing less of a hitchhiking effect. We refer to the mode of Timesweeper that uses this information to detect sweeps as the Allele Frequency Tracker (AFT). The AFT format specifically examines, at each site in its window, the frequency of the allele that has increased the most from the first to the final timepoint. Our motivation for this is that this will better capture the positive shift in frequency for both the sweeping and hitchhiking alleles, a signal that our network should have little difficulty in detecting. If we instead tracked the frequency of the derived alleles (or minor allele in unpolarized data), this would yield a mixture of both positive and negative changes in allele frequency in the presence of hitchhiking, which might be harder for the network to identify as the consequence of a sweep.

The second data representation is a *k* × *m* matrix, where *k* is the number of distinct haplotypes observed across all timepoints and *m* is again the number of timepoints. The haplotypes in this matrix are sorted so that the haplotype with the highest net change in frequency across the sampling period is shown at the bottom, and the remaining haplotypes are sorted by similarity to this focal haplotype which is *a priori* the one most likely to be experiencing any positive selection. This second representation thus tracks the frequency changes of all haplotypes observed in the entire sample, which should be especially pronounced for any haplotypes harboring a sweeping allele. We refer to the mode of Timesweeper that uses data in this format as the Haplotype Frequency Tracker (HFT). Both of these formats allow for easy input into convolutional neural networks, and the two modes of Timesweeper both use 1D convolutions over the time axis to track allele or haplotype frequency changes through time, respectively (Methods).

Figure 1 shows the expected values of each entry in these matrices, as computed from 10,000 forward simulations of selective sweeps under a constant population size of 500 and a selection coefficient, *s*, drawn from a uniform distribution with bounds [0.00025, 0.25) (Methods). This visualization shows that the rapid spread of selected alleles (the central row in the sweep examples shown in Figure 1A) and haplotypes (the bottom row in Figure 1B) is readily observable in this representation of the mean, and the selected and unselected cases are readily discernable. However, we note that individual selective sweeps may depart substantially from this expectation, and we illustrate this by showing randomly chosen individual examples from each class in Figure S1A and S1B. In the next section we investigate the extent to which individual examples are correctly classified as undergoing selective sweeps or evolving neutrally on the basis of these representations.

### Neural networks can detect and localize sweeps from time-series data with high accuracy

Having constructed the data representations described above, we sought to train neural networks to distinguish between three distinct classes: 1) Neutrally evolving regions, 2) Sweeps caused by selection on a single-origin *de novo* mutation (the SDN model, which produces classic “hard” selective sweeps; (Smith and Haigh 1974)), and 3) selection on standing variation (the SSV model, which may produce “soft sweeps” involving multiple distinct haplotypes carrying the selected allele; see (Orr and Betancourt 2001; Hermisson and Pennings 2005)). Other approaches using these data are possible—for example, one could adapt existing composite-likelihood ratio approaches (Nielsen *et al*. 2005; Vy and Kim 2015) to use the information in the AFT representation to detect sweeps from time-series data. However, a simulation-based CNN would allow us to model the joint distribution of all of the allele or haplotype frequency changes observed in the vicinity of a sweep. As we show below, this approach yields excellent inferential power.

Here and in the following sections, we show that allele frequency trajectories are highly informative when processed by 1DCNNs, demonstrating high accuracy across all time-series schemes tested. We began by benchmarking performance on a scenario of 20 timepoints with 10 individuals at each timepoint on a test data set held out during training, and found that the AFT method achieves an area under the ROC curve (AUROC) of 0.95 when selection coefficient *s* is drawn from a uniform distribution with bounds [0.00025, 0.25) and the sweep began at a time ranging uniformly from 50 generations prior to the first sampling timepoint to 50 generations after this sampling timepoint (Figure 2A); we examine more combinations of these parameters in the section below. The HFT method is somewhat less accurate, achieving an AUROC of 0.87. Qualitatively similar results are shown in the precision-recall curves in Figure 2B. We find that Timesweeper can estimate *s* with high accuracy across a range of selection strengths with both the AFT and HFT data formats (Figure 1 C-D).

**Figure 2.**
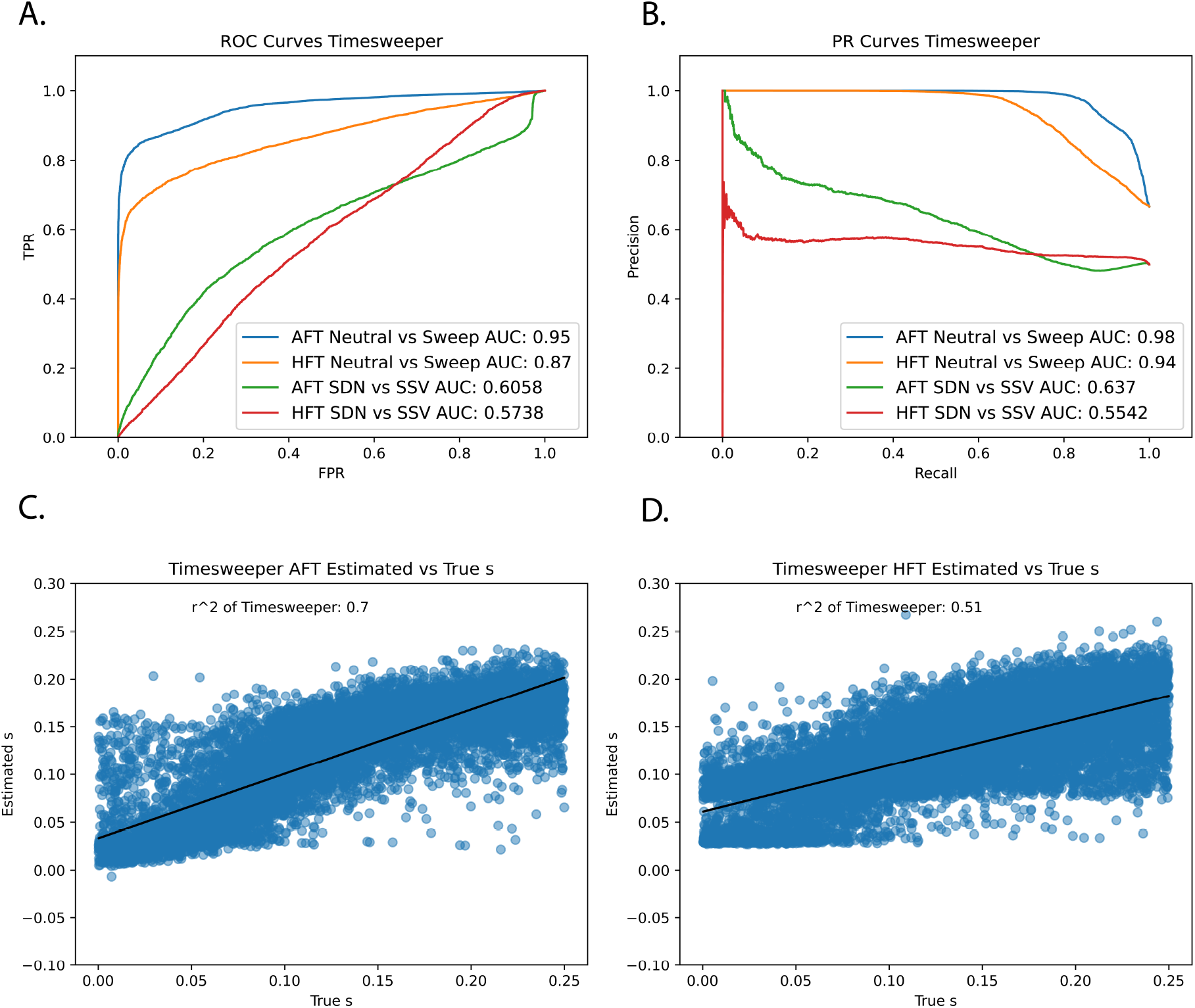
Timesweeper accuracy benchmarking. (A) ROC curves and (B) PR curves for AFT and HFT formats for both Neutral vs Sweep and SDN vs SSV binary classification tasks. (C-D) Estimated vs True selection coefficient for C) AFT and D) HFT. Neural networks were trained on a dataset of 10,000 replicates for each of the neutral, SDN, and SSV classes, with selection coefficients for the latter two classes drawn from a uniform distribution with bounds [0.00025, 0.25), and sampling start drawn from a uniform distribution with bounds [-50, 50) generations from the onset of selection. Models were tested on 5,000 independently simulated replicates for each class that parameterized identically to the training data.

The results described above were obtained by examining simulated test data where either the central polymorphism was selected, or not, and making a single classification for this focal site (hereafter referred to as single-locus testing). However, in practice one would scan across an entire chromosome and make classifications at each polymorphism before moving on to the next. Thus, we next benchmarked the same Timesweeper classifiers described above on a set of 15,000 simulated chromosomes 500 kilobases long—5,000 replicates each of the neutral, SSV, and SDN scenarios. We found that Timesweeper was not only able to accurately detect true sweep locations but has a low false positive rate across the flanking neutral regions along the chromosome (Figures S3-4). The HFT method, on the other hand, exhibits comparatively low resolution in localizing sweeps (Figures S3-4), likely because it examines a representation of haplotype frequencies that has no spatial information about any potentially sweeping alleles. Timesweeper also performs classification accurately for a variety of selection coefficients using the sampling scheme described above (Figures S5-7). Testing on a range of *s* we found that AUROC is minimized at 0.79 with *s*=0.01 and maximized at 0.99 with *s*=0.5.

Training each network on 21,000 replicates (7,000 of each class) without GPUs required roughly one minute on 4 CPU cores and less than 1Gb of RAM. Classifying a simulated test set of 1,500 replicates for each class with 10 diploid individuals sampled at each of 20 timepoints required roughly 60 seconds for each trained neural network, resulting in a total of 6 minutes runtime when both the AFT and HFT methods are utilized.

### Choice of Neural Network Architecture

There are multiple neural network architectures suited to time-series or sequence data (1DCNN, transformers, RNNs) as well as models suited to spatial data (2DCNNs and variants). We initially chose a 1DCNN architecture due to its lightweight architecture and demonstrated performance in population genetics tasks (Flagel *et al*. 2019). 1DCNNs have the advantage of being more computationally efficient than 2DCNNs and theoretically can capture the entire spatial axis (i.e. each of the *l* polymorphisms in the *l* x *m* AFT feature matrix) at each time-step of a sequence, as opposed to a 2DCNN that convolves over both axes in a blocked fashion and thus could miss associations between the selected site and more distant loci that would be expected if a sweep occurred at the target locus, thereby potentially missing some evidence of hitchhiking information that can only be obtained by observing the entire window a timepoint or set of timepoints. We tested this assumption by implementing and training a variety of common neural network architectures to benchmark held-out test accuracies. We found that the 1DCNN performed marginally better than all competing architectures while simultaneously being the fastest by far (2- fold faster than RNN, over 10-fold faster than 2DCNN; Figures S8-9).

### Understanding feature importance with saliency maps

In order to make our models more interpretable and to better understand what features are explicitly used by the network, we adopted the approach of Gower *et al*. (2021), who trained a CNN to detect adaptive introgression. We produced saliency maps for our 2DCNN architecture using the tf-keras-vis library (Kubota 2022) for both the AFT and HFT data formats. We found that the AFT and HFT networks both have high levels of saliency in the regions that one might expect to be most important for identifying sweeps: the central-most polymorphism in the AFT format and the bottom-most haplotype in the HFT format, with rapid decay of saliency at increasing distances from the focal polymorphism/haplotype. In addition, we observe that saliency is highest at the earliest and latest timepoints in the series. Together these observations suggest that the network is searching for direct evidence of focal alleles/haplotypes that have increased in frequency across the sampling interval. However, we emphasize that even though attention is much higher at target site in the AFT network, more distant loci do influence the predictions of our AFT 1DCNN, a point that we address in the following section.

### Localization of sweeps improves when flanking sites are incorporated

In order to test the optimal window size (the number of polymorphisms, *l*) for training and testing a 1DCNN we trained and tested sizes *l*=1, 3, 11, 51, 101, and 201. For each window size the focal polymorphism is located in the center of the window, i.e. for *l*=51 the 26^th^ polymorphism in the window is the focal site. When assessing classification and regression accuracy on these various window sizes, we applied the Timesweeper to not only the central polymorphism in each of 5,000 test replicates for each class, but to all other polymorphisms within these simulated chromosomes as well. This allows us to address 1) how accurately Timesweeper detects sweeps/infers s for a given window size, and 2) how precisely Timesweeper localizes the sweep (i.e. to what extent it avoids misclassifying polymorphisms closely linked to the selected allele as being themselves selected).

When examining only the focal polymorphism within each simulated example, we found that for AFT there is no noticeable improvement in classification accuracy when increasing from a single allele to larger windows (AUROC ranges varies from 0.94 to 0.97 across all window sizes, with no noticeable trend as *l* increases). This may be expected, as if one’s goal is only to determine whether a single pre-identified polymorphism is under selection, adding data from flanking sites will yield little information that is not already captured by the allele frequency trajectory— although we note that this may not hold for smaller numbers of timepoints, a possibility that we did not investigate here. Interestingly, HFT performs worse the larger the window size becomes (AUROC of 0.96 for *l* =1 vs 0.80 for *l*=201; Figures S11-13). The regression task follows a similar trend, with no major differences noted across the different window sizes for AFT and decreasing performance with larger window sizes for HFT (Figure S14).

In the context of genome-wide scans for selection, one will test many sites potentially affected by selection, and in this context our goal is not only accurately detect sweeps, but to localize them to a reasonably small candidate region. We found that, although small window sizes produced accurate inferences at the target sites themselves, they were much more likely to misclassify closely linked polymorphisms as being targets of selection as well than were networks trained on larger window sizes (Figures S3-4). Indeed, the number of linked polymorphisms within 500 bp of the true target of selection that are misclassified as sweeps decreases from 50 to 39 as *l* increases from 1 to 201. We have adopted *l*=51 as our default as it appears to reach a good balance between accuracy and computational speed (as larger window sizes require more trainable network parameters), but if users wish to use larger values when running Timesweeper this is an option. We note that the optimal value for any given dataset may depend on the density of polymorphism, the recombination rate, and the distribution of selection coefficients, so we encourage users to experiment with different window sizes before adopting one for their analysis.

### The impact of sampling scheme and selection strength on accuracy to detect selection

To test effectiveness of Timesweeper under different sampling scenarios we experimented with varying five parameters: the sample size per timepoint, the number of timepoints sampled, the timing of sampling relative to the onset of selection, the selection coefficient, and the size of the training set. For each experiment described below, the parameters of the training set matched those of the test set. With the exception of these parameter changes, all simulation, training, and testing was performed in the same manner as above. Each experimental model reported in this section was trained on 8,500 replicates for each class (SDN, SSV, and neutral classes), and tested on 1,500 held-out test replicates for each class for each parameter tested.

#### Sample size

We examined the effect of sample size on classification accuracy by training Timesweeper for each of four scenarios: 1, 2, 5, 10, and 20 diploid individuals sampled at each of 20 timepoints. Again, the beginning of the sampling interval ranged from 50 generations before to 50 generations after the onset of selection for both the SDN and SSV scenarios, and other than sample sizes all other components of the simulation were identical (example and average inputs shown in S2 Fig). We found that increasing number of samples has little effect on the performance of the AFT model: with one diploid individual per timepoint the AUROC for the AFT method was 0.95, and with 20 individuals per timepoint this had only increased to 0.96 (Figures S15-17). While HFT had a noticeable increase in accuracy with increased sample sizes (AUROC of 0.81 for one diploid individual per timepoint increased to 0.87 AUROC for 20 individuals) it was consistently less accurate than AFT across all sample sizes (Figures S15-17).

While increased sample sizes had no drastic effect for classification accuracy, there was a sizeable difference in accuracy of selection coefficient estimates with increased sample sizes (*R^2^* of 0.65 for one individual to 0.81 for 20 individuals for AFT, 0.47 to 0.69 for HFT for SDN scenario estimates; Figure S18). These results indicate that Timesweeper is flexible enough to handle small sample sizes and still accurately detect sweeps, but will obtain more accurate estimates *s* when larger sample sizes throughout the time series are available.

#### Number of timepoints

Next, we tested the effect of adjusting the number of timepoints on Timesweeper’s accuracy. In this scenario the total number of sampled individuals was 200 for every sampling scheme, so accuracy would be solely affected by the number of timepoints and their timing relative to the sweep as opposed to the total amount of genetic data being examined. Scenarios tested included 1, 2, 5, 10, 20, and 40 timepoints. In all scenarios, aside from the single- timepoint scenario, the sampling times were evenly distributed across a 200-generation span starting 50 generations after the onset of selection. For the single-timepoint scenario only the final generation of this 200-generation span was sampled (i.e. a sample of 200 individuals was taken 250 generations after the onset of selection). Among time-series schemes, there is an initial accuracy increase from 1 (AUROC of 0.76) to 2 timepoints (AUROC of 0.94), however there is only a marginal increase in performance as the number of timepoints increases to 40 (AUROC of 0.96; Figures S19-21). There is a similar increase in performance for the 3-class classification accuracy, with the majority of misclassifications being between SDN and SSV scenarios, and few false positives or negatives (sweeps labeled as neutral or vice versa; see confusion matrices in Figure S19). Overall 3-class classification performance improves with more than 1 timepoint (47% accuracy) to 69% with 2 timepoints, and only marginal improvement with more timepoints (71% accuracy for 40 timepoints). These results indicate Timesweeper is accurate under a variety of sampling schemes, and even small numbers of sampling timepoints may yield substantially better performance than single-timepoint data. For selection coefficient estimation, there is a large jump in accuracy from 1 (*R^2^* of 0.37 for SDN *s* estimates) to 2 (0.44) to 5 timepoints (0.78), after which there are marginal increases up to 40 timepoints (*R^2^* of 0.83). Thus, the lion’s share of Timesweeper’s performance gains are achieved by adding only a reasonably small number of timepoints (Figure S22).

To determine whether the higher accuracy observed for sampling schemes with more timepoints was a result of their being more evolutionary information present in the dataset or the larger number of parameters in the networks trained for data with more timepoints, we tested two networks examining the same number of timepoints (20) in their input but with a large difference in the number of weights: one 1DCNN architecture containing 11,321,107 parameters, and another with 421,835 parameters (the standard 1DCNN model used elsewhere in the manuscript). We did not observe a substantial difference in accuracy between these two networks (Figures S8-9), suggesting that it is the number of timepoints driving the differences in performance seen in Figures S19-22.

#### Post-selection timing

We also simulated a set of scenarios that test the effect of varying when sampling occurs relative to the start the sweep. In these experiments all scenarios were simulated with 20 sampled timepoints each with 10 sampled individuals. The sampling window for each remained a span of 200 generations with the 20 timepoints evenly spaced within. The starting time for the sampling window was set to either -100, -50, 0, 25, 50, 100, or 200 generations after the onset of selection. Timesweeper performed best (AUROC of 0.96) on data sampled beginning -100 to 100 generations post-selection (AUROC 0.94-0.96), with the lowest AUROC observed being 0.72 AUROC for sampling beginning 200 generations post-selection. Reduced accuracy under sampling schemes occurring very late relative to the onset of selection is expected, as some sweeps may be at or near fixation prior to the completion of sampling—in these cases the sojourn of the sweeping mutation would be largely/entirely absent from the time series, and for those sweeps completing prior to sampling, the sweep signature may even have begun to erode before sampling. In a similar vein, we found that Timesweeper estimated selection coefficients most accurately when the initial sampling time ranged from -100 generations (*R^2^* of 0.79 for SDN *s*) to 50 generations (0.85) post-selection, with a steep drop in accuracy at 200 generations post-selection (Figure S26).

We also found that Timesweeper was able to differentiate between SDN and SSV sweeps with high accuracy (SDN vs SSV AUROC ranging from 0.78 to 0.82) for most starting times (-100 to 50 generations post-selection). We hypothesize the model performs so well in these cases due to the earlier starting times capturing the full onset of selection and the initial frequency of the sweeping allele. These tests show that Timesweeper can be trained to accurately detect sweeps that occurred at a variety of times relative to the sampling period, and is best able to differentiate between SDN and SSV sweeps when the sampling scheme captures as much of the sweep trajectory as possible (Figures S23-26).

#### Selection coefficient

We simulated training and test sets with selection coefficients of 0.005, 0.01, 0.05, 0.1, and 0.5. For each of these simulations, we sampled at 20 timepoints, with the first sampling timepoint again occurring between 50 generations before and 50 generations after the onset of selection, and sampled 10 diploid individuals at each timepoint. We find that, for the AFT, accuracy is fairly low when *s*=0.005 (AUROC=0.81 for distinguishing between sweeps and neutrality) but increases rapidly with *s* (AUROC=0.97, 0.98, 0.99 when *s*=0.05*, s*=0.1, and *s*=0.5, respectively; Figures S5-S7).

#### Number of simulation replicates

We simulated 30,000 replicates for each scenario (neutral, SDN, SSV) and trained and tested on our network on datasets of varying size (30,000, 20,000, 10,000, 5,000, 2,000, and 1,000 replicates of each scenario). For each size a 75/15/15% train/test/validation split was performed as described in the Methods. We found that the number of replicates have little effect on accuracy of either AFT or HFT sweep classification (0.94 to 0.96 AUROC from 1,000 to 30,000 replicates for AFT, 0.86 to 0.87 for HFT) (Figures S27-29), but had a large impact on *s* estimation. After an initial jump from 1,000 replicates (*R^2^* of 0.46 for SDN *s* estimates) to 2,000 replicates (0.75) to 5,000 (0.84), accuracy continued to increase, albeit at an attenuated rate, even at 30,000 replicates (*R^2^* of 0.87; Figure S30).

### Timesweeper is relatively robust to the effects of demographic model misspecification

As with all parametric approaches to population genetic inference, the demographic model used could impact the accuracy of downstream inference (Johri *et al*. 2022a; b). To characterize Timesweeper’s AFT accuracy in of the presence of highly misspecified demographic history we performed cross-model benchmarking between three different demographic models using the (see Methods): a constant-size demographic model (referred to as the constant model), a 3-epoch bottleneck model (the bottleneck model) as described in (Marth *et al*. 2004) and the Out-of-Africa model (the OoA model) described in (Gutenkunst *et al*. 2009).

We found that overall Timesweeper retains high classification accuracy even in the case of extreme model misspecification of training the classifier under the constant model and applying it to the OoA models, with an AUROC of 0.86 when trained on the constant model and tested on OoA, and an AUROC of 0.92 when trained on OoA and tested on the simple model. Indeed, as shown in Table 1 and Figure S32, there is almost no decrease in AUROC values for detecting sweeps when training on the constant model and testing the bottleneck and OoA models compared to networks that are trained on and applied to data from the same non-equilibrium demographic model. Thus, the reduction in accuracy when performing inference on the non-equilibrium models is a consequence of sweep signals being harder to detect in these models rather than misspecification. A modest reduction in the ability to distinguish SSV from SDN sweeps is observed, however, when training on the constant model and applying to the bottleneck or OoA models. Demographic misspecification has a much larger impact on selection coefficient estimation, and we observe much larger losses in accuracy when predicting on the bottleneck and OoA models after training on the constant model (Table 1, Figure S33).

**Table 1.** Timesweeper. ’s classification and regression accuracy when trained on properly specified and misspecified demographic models.

| **<u>Trained on constant</u>** | **Tested on constant** | **Tested on bottleneck** | **Tested on OoA** |
| --- | --- | --- | --- |
| Sweep AUROC | 0.94 | 0.94 | 0.86 |
| SDN vs SSV AUROC | 0.71 | 0.63 | 0.54 |
| Sweep AUPR | 0.97 | 0.98 | 0.93 |
| SDN vs SSV AUPR | 0.69 | 0.58 | 0.51 |
| SSV $R^2$ | 0.58 | 0.78 | -0.41 |
| SDN $R^2$ | 0.81 | 0.66 | -0.39 |
| **<u>Trained on bottleneck</u>** | **Tested on constant** | **Tested on bottleneck** | **Tested on OoA** |
| Sweep AUROC | 0.93 | 0.94 | 0.85 |
| SDN vs SSV AUROC | 0.71 | 0.63 | 0.53 |
| Sweep AUPR | 0.97 | 0.98 | 0.92 |
| SDN vs SSV AUPR | 0.69 | 0.58 | 0.51 |
| SSV $R^2$ | 0.43 | 0.79 | -0.71 |
| SDN $R^2$ | 0.78 | 0.66 | -0.66 |

| **Trained on OoA** | **Tested on constant** | **Tested on bottleneck** | **Tested on OoA** |
| --- | --- | --- | --- |
| Sweep AUROC | 0.92 | 0.94 | 0.86 |
| SDN vs SSV AUROC | 0.68 | 0.56 | 0.53 |
| Sweep AUPR | 0.96 | 0.98 | 0.93 |
| SDN vs SSV AUPR | 0.64 | 0.51 | 0.51 |
| SSV R<sup>2</sup> | 0.48 | 0.61 | 0.37 |
| SDN R<sup>2</sup> | 0.62 | 0.63 | 0.44 |

### Timesweeper outperforms previous methods

We compared Timesweeper to a variety of other methods created for the task of either detecting sweeps or estimating selection coefficients in both time-series and single-point data (Methods). For sweep detection/classification we benchmarked against two single-timepoint methods, diploSHIC (Kern and Schrider 2018) and Sweepfinder2 (DeGiorgio *et al*. 2016), and two time-series methods, the Frequency Increment Test (FIT) (Feder *et al*. 2014) and Fisher’s Exact Test (previously implemented in PoPoolation2 for this purpose; (Kofler *et al*. 2011)). We also extended diploSHIC to work with time-series data, an approach we call TS-SHIC (Methods), and included this method in the comparison. For the diploSHIC variants, we assessed accuracy in discriminating between SDN and SSV sweeps, and for these and all other classification methods we assessed accuracy in discriminating between sweeps and neutrally evolving regions. We found that Timesweeper’s AFT outperformed the other methods both in discriminating between sweeps and neutrally evolving regions (AUROC of 0.96 for Timesweeper’s AFT, vs 0.92 for the next-best method), and in distinguishing between SDN and SSV sweeps (AUROC of 0.76 for Timesweeper’s AFT, 0.69 for Timesweeper’s HFT, and 0.63 or less for the diploSHIC variants; Table 2, Figures 3,S31).

**Figure 3.**
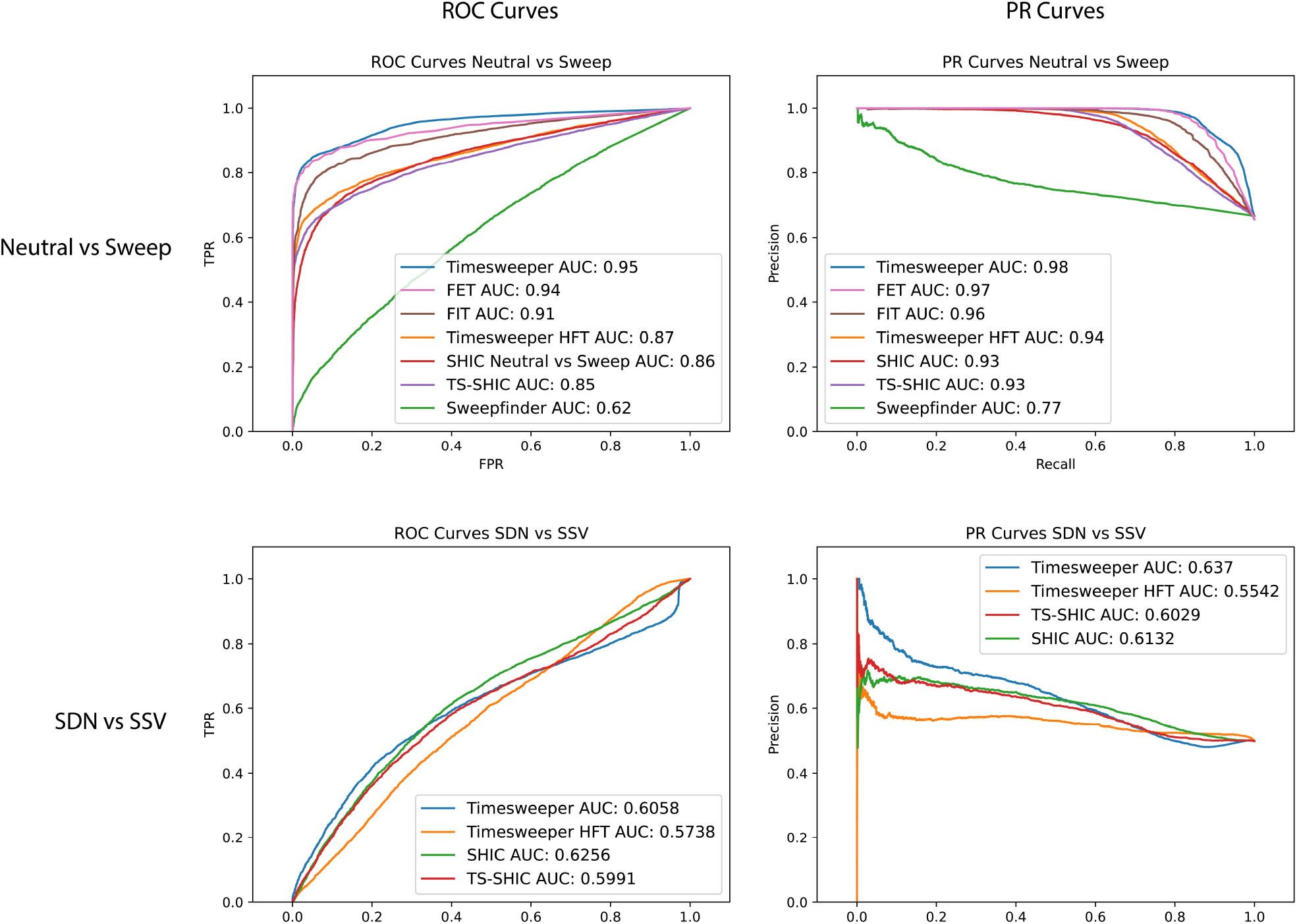
ROC and PR curves comparing the performance of **Timesweeper** to that of competing methods on the sweep detection task. The top row shows performance on the binary task of discriminating between simulated test data with and without selective sweeps, with the sweep class containing equal numbers of SDN and SSV examples. The bottom row shows performance on distinguishing between SDN and SSV examples in the test set.

**Table 2.** Accuracy of **Timesweeper** and competing methods on the sweep detection task.

| Classification Method | Sweep vs Neutral AUROC | SDN vs SSV AUROC | Sweep vs Neutral AUPR | SDN vs SSV AUPR |
| --- | --- | --- | --- | --- |
| Timesweeper AFT 1DCNN | 0.95 | 0.61 | 0.98 | 0.64 |
| Fisher's Exact Test (FET) | 0.94 |  | 0.97 |  |
| Frequency Increment Test (FIT) | 0.91 |  | 0.96 |  |
| Timesweeper HFT 1DCNN | 0.87 | 0.57 | 0.94 | 0.55 |
| SHIC | 0.86 | 0.63 | 0.93 | 0.6 |
| TS-SHIC | 0.85 | 0.59 | 0.93 | 0.6 |
| Sweepfinder2 | 0.62 |  | 0.77 |  |

**Table 3.** Accuracy of **Timesweeper** and competing methods on the selection coefficient estimation task.

| Selection coefficient inference Method | R2 |
| --- | --- |
| Timesweeper AFT 1DCNN | 0.7 |
| ApproxWF | 0.57 |
| Timesweeper HFT 1DCNN | 0.51 |
| TS-SHIC | 0.4 |
| WFABC | 0.34 |
| slattice | -1.03 |

For *s-*estimation we compared our performance to three existing time-series methods: slattice (Mathieson and McVean 2013), ApproxWF (Ferrer-Admetlla *et al*. 2016), and WFABC (Foll *et al*. 2015). We also adapted the time-series implementation of diploSHIC to estimate *s*, and included this in our comparison. Again, we found that Timesweeper’’s AFT method outperformed all other methods (*R*^2^ between estimated and true *s* of 0.77; Table 3), with ApproxWF being the next best performer (*R*^2^=0.6). Note that slattice obtained a negative *R*^2^ value, because the magnitude of error was quite large for a substantial fraction of cases, but visual inspection shows that many predictions by this methods were quite accurate (Figure 4).

**Figure 4.**
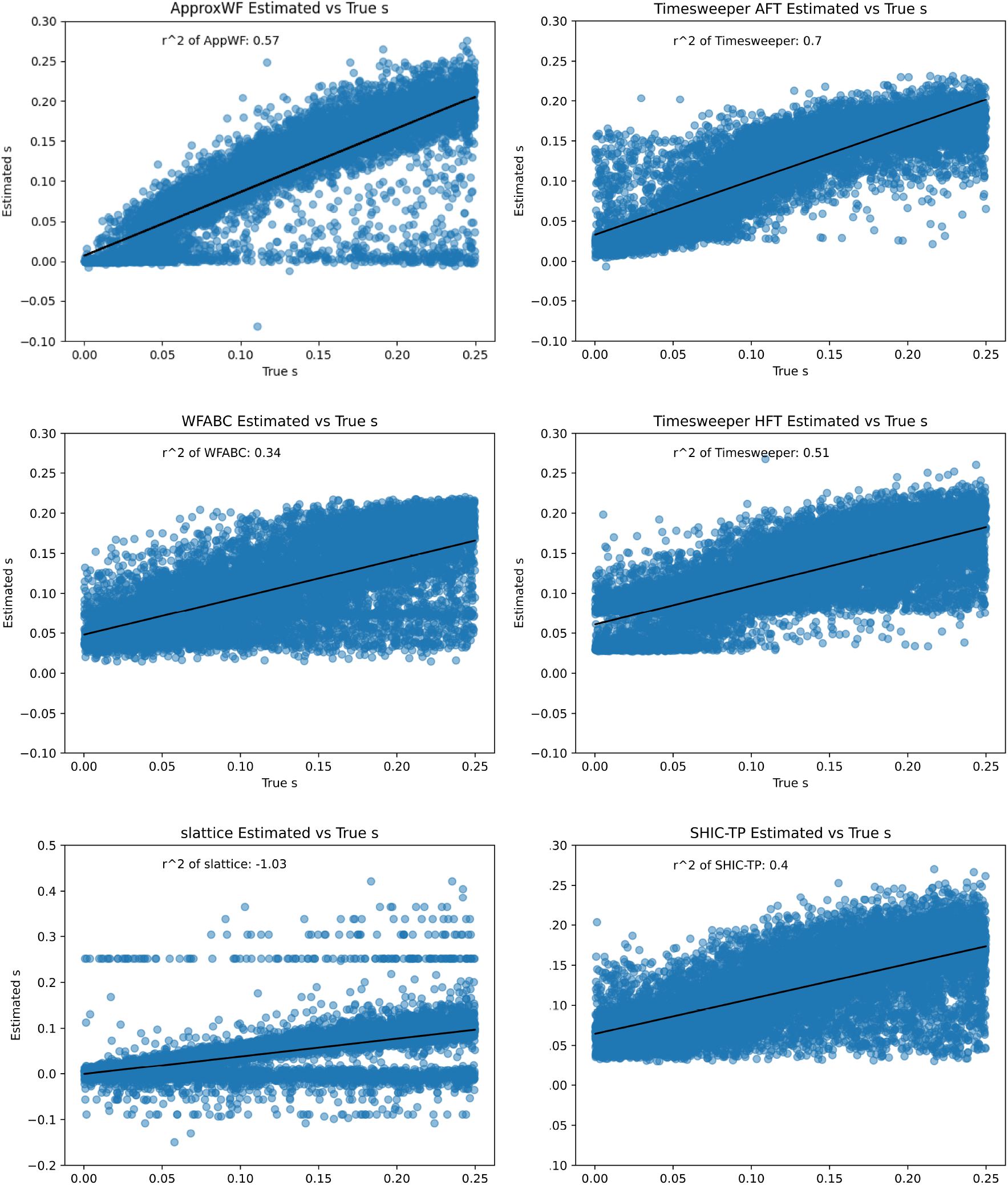
Comparison of **Timesweeper** to competing methods in the s inference task. Each method was applied to a test set containing an equal number of SDN and SSV sweep examples.

**Figure 5.**
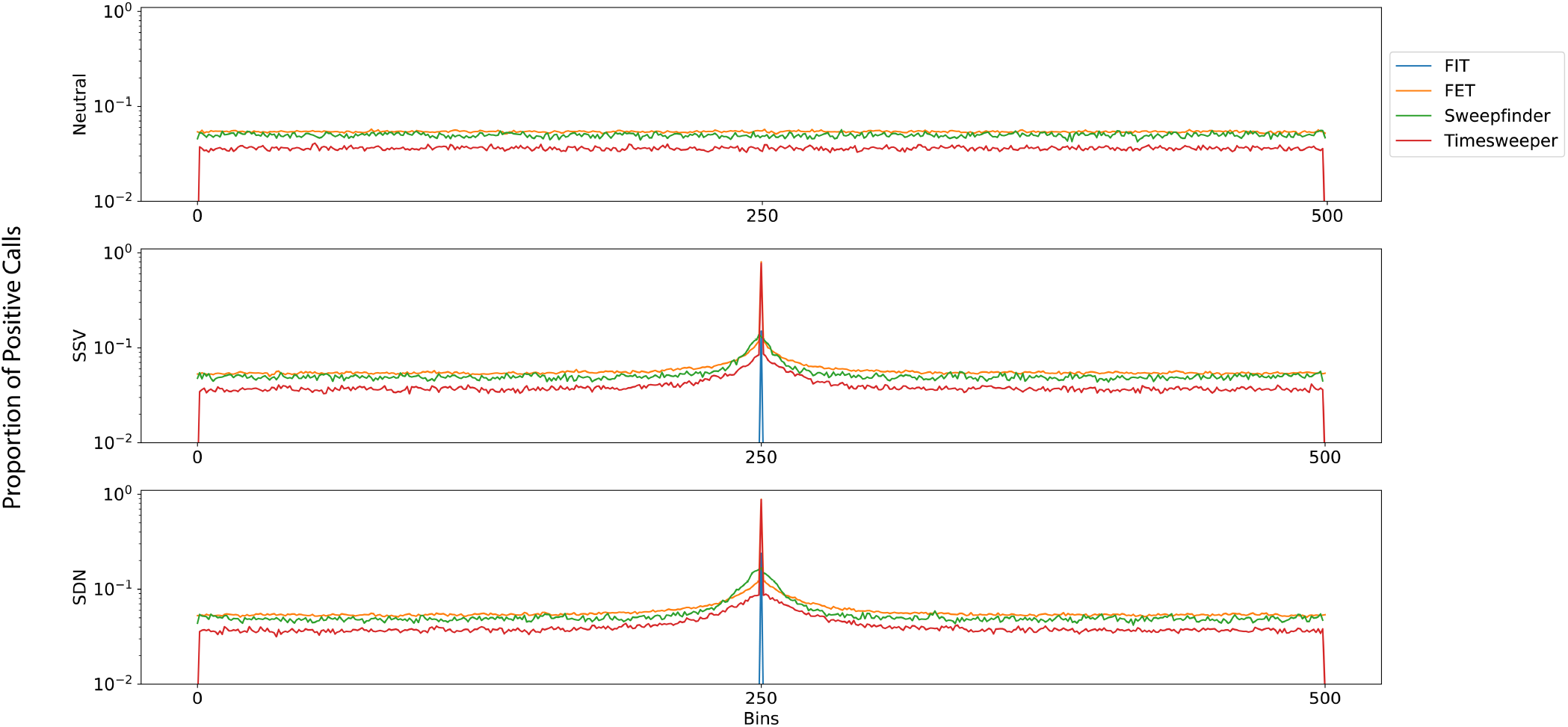
Fraction of sweep class within a 500kb region, averaged across 5,000 replicates each of Neutral (top), SSV (middle), and SDN (bottom) scenarios. Calls are binned into 500 equally-sized bins of polymorphisms along the chromosome. For the SSV and SDN scenarios, the central bin consists only of the polymorphism under selection.

**Table 3.** The numbers of top sweep candidates replicated by Timesweeper and Fisher’s Exact Test (FET) on data from experimental D. simulans populations. SNPs among the top 100 scoring sweep candidates for a given replicate were considered to be replicated if they were also found in the top-scoring 1% predictions by that same method in another replicate. Note that some candidate SNPs were replicated in more than one of the 9 additional experimental replicates (see Figure S36), and thus a total number of replicated hits >100 is possible.

| Replicate | Timesweeper's Replicated Hits | FET's Replicated Hits |
| --- | --- | --- |
| 1 | 44 | 11 |
| 2 | 97 | 86 |
| 3 | 44 | 144 |
| 4 | 43 | 43 |
| 5 | 37 | 17 |
| 6 | 43 | 18 |
| 7 | 30 | 46 |
| 8 | 20 | 8 |
| 9 | 52 | 18 |
| 10 | 47 | 21 |

### Detecting Sweeps in an experimentally evolved *Drosophila simulans* population

In order to assess Timesweeper’s utility on experimental evolution data we applied it to data from a publicly available *Drosophila simulans* study published by Barghi et al. (2019). This study contains 10 experimental replicates, each exposed to a warm environment for 60 generations, giving us the ability to assess our method’s performance on real data by examining the extent to which Timesweeper’s sweep candidates were replicated across these 10 datasets. We compare our results to those obtained by Barghi et al., who used Fisher’s Exact Test (FET) to assess the significance of the difference between the starting versus final allele frequency for a target polymorphism within a given experimental replicate.

In brief, we simulated a training set to match the experimental design described in Barghi et al. and used these simulations to train a Timesweeper AFT classifier as described in Methods. We note that this experiment lasted for a relatively short period of time (60 generations), such that positively selected *de novo* mutations would rarely be expected to arise and reach high frequency by the end of the experiment; we therefore trained Timesweeper to distinguish between neutrality and the SSV model (example and average inputs in Figure S34), and refer to the latter as the “sweep” model for the remainder of this section. We then used the resulting classifier, which we found to be highly accurate (AUROC=0.96; Figure S35) to scan allele frequency estimates from the pooled-sequencing data from Barghi et al for signatures of sweeps on each of the five major chromosome arms (Methods). This scan was performed independently in each of the 10 experimental replicates. The resulting predictions were filtered to retain the top 1% of the class score distribution, these prediction scores were then compared to the top 1% of the reported - log10(*p*-value) FET scores. We found that Timesweeper predictions have a Spearman correlation of 0.20 with FET scores. Although this correlation across the whole range of scores was modest, we found that SNPs receiving the largest sweep probabilities from Timesweeper have high FET scores and vice-versa (Tables S1 and S2).

We next investigated the degree to which the top sweep candidate SNPs (the 100 highest-ranking SNPs) from a given replicate were also classified as sweeps in other replicates (according to a less stringent cutoff of whether the SNP was in the top 1% of sweep scores in the replicate population). This was done for both Timesweeper and FET’s top positive selection candidates, using the sweep class-membership probability as the score for Timesweeper, and the -log10(*p*- value) as the sweep score for FET. We found that Timesweeper replicated a larger number of hits on average than FET with the exception of replicate 3 (Table 3). Upon closer examination, we found that this was due to the presence of a large sweep region on chromosome 3 where FET predicted high scores for a large number of polymorphisms in the surrounding region in multiple replicates. Importantly, Timesweeper was able to detect and replicate the sweep signature in this region, it just did not classify as many SNPs in this region as sweeps as FET did. This illustrates that these numbers of replicated classifications can be somewhat inflated by linked SNPs receiving the same classification. One would expect this is probably a greater issue for the FET (which evaluates each SNP independently) than Timesweeper, which evaluates a focal SNP while including information about flanking SNPs as well.

To more directly test the notion that Timesweeper’s replication rate is less affected by linked selection, we subdivided the genome into adjacent 100 kb windows and asked, for each window with at least one top SNP candidate, how many additional experimental replicates also contained a strong selection signature somewhere in this window. Because each window is counted only once, regardless of how many significant SNPs in contains, this approach is less influenced by the same sweep signature being counted multiple times. As shown in Figure S36, windows inferred by Timesweeper to contain sweeping mutation are much more likely to be replicated and are typically recovered in a larger number of experimental replicates. In summary, the consistent ability of Timesweeper to identify regions that show signals of selective sweeps across replicates underscores its ability to detect positive selection in real data sets.

## DISCUSSION

Detecting recent positive selection is an important problem in population genetics. Such efforts can help reveal the extent and mode of recent adaptation, as well as clues about the genetic and phenotypic targets of selection (Nielsen *et al*. 2005; Voight *et al*. 2006; Garud *et al*. 2015). Moreover, signatures of selective sweeps have been shown to co-localize with disease-associated mutations (Blekhman *et al*. 2008; Chun and Fay 2011; Schrider and Kern 2017), although one recent study has argued that Mendelian disease genes are less likely to exhibit signatures of selection than non-disease genes after controlling for potentially confounding factors such as the number of protein-protein interactions (Di *et al*. 2021). One possible explanation is that deleterious alleles on the positively selected haplotype may hitchhike to higher frequencies than they might otherwise reach (Chun and Fay 2011), although other potential explanations exist (Otto 2004; Corbett *et al*. 2018).

For these reasons, there has been a great deal of effort to develop methods to detect the signatures of positive selection from a single sample of recently collected genomes (e.g. (Kelly 1997; Fay and Wu 2000; Kim and Stephan 2002; Sabeti *et al*. 2002; Kim and Nielsen 2004; Voight *et al*. 2006; Li 2011b; Ferrer-Admetlla *et al*. 2014; Harris and DeGiorgio 2020). Recent approaches have incorporated combinations of these tests through the use of machine learning (e.g. (Lin *et al*. 2011; Ronen *et al*. 2013; Pybus *et al*. 2015; Schrider and Kern 2016; Sugden *et al*. 2018; Gower *et al*. 2021), and it may even be feasible to completely bypass the step of computing summary statistics by training deep neural networks to operate directly on population genomic sequence alignments as input, as has been done effectively for other problems in population genetics (Chan *et al*. 2018; Flagel *et al*. 2019; Adrion *et al*. 2020b; Sanchez *et al*. 2021).

While the use of modern data to make inferences about the past is the cornerstone of evolutionary genomics, one could potentially achieve greater statistical power by including genomic samples across a variety of timepoints, allowing for a more direct interrogation of evolutionary history. Genomic time-series data has been used to great effect to find positively selected loci in experimentally controlled studies (Schlötterer *et al*. 2015). There is growing interest in performing similar analyses in natural populations, as the acquisition of time-series genomic data has become feasible both in species with short generation times such as *Drosophila* (Bergland *et al*. 2014; Machado *et al*. 2021; Kapun *et al*. 2021; Lange *et al*. 2022), and in those for which ancient DNA is available, such as humans (Allentoft *et al*. 2022).

With the potential benefits of time-series data and machine learning methods for population genetics in mind, we developed Timesweeper, a deep learning method for detecting selective sweeps and inferring selection coefficients from population genomic time series. We experimented with two approaches: one using a matrix representation of haplotype frequencies estimated in each sampled timepoint, and one tracking allele frequencies through time in a window of polymorphisms centered around a focal polymorphism. We found that while the haplotype frequency tracker (HFT) had decent power to detect selective sweeps, the allele frequency tracker (AFT), when compared to competing methods, had superior accuracy and was able to localize selected variants with much higher resolution, and accurately infer selection coefficients. We note that while the AFT method was able to distinguish between the SDN and SSV selection models with greater accuracy than competing methods, accuracy on this task was considerably lower than that for distinguishing between selection and neutrality. This is partially a consequence of how we defined these models: the SSV model involves selection on a previously standing polymorphism, but if by chance only a single ancestral copy of the adaptive allele survives the sweep, then the result will be indistinguishable from the SDN model.

We also note that the next-best method for identifying sweeps on our initial benchmarking dataset was the simplest: Fisher’s exact test (FET) for allele frequency differences between the first and final timepoint. We note that FET’s performance dropped considerably when applied to our test datasets with non-equilibrium demographic histories (Figure S32), demonstrating that this approach can only be used in settings where one knows the degree of genetic drift over the course of the sampling interval will be small—scenarios involving periods of small effective populations sizes, which many species experience in nature, and/or long sampling intervals will be inappropriate for FET. In addition, we found that Timesweeper is better able to narrow down the target of selection, because it examines flanking polymorphisms, perhaps giving it some power to distinguish between the selected locus and linked regions containing hitchhiking alleles, whereas FET examines each polymorphism separately and an allele with a large enough frequency change will result in a rejection of the null. Nonetheless, in some settings a simple method such as FET may be preferable, such as experimental studies where the population size is held constant and the sampling interval is sufficiently short (although more complex methods would be required if one wishes to infer the mode of selection, or estimate *s* or other parameters of the sweep).

When evaluating the performance of model-based methods for population genetic inference, it is important to consider the impact of misspecified demographic histories (Schrider and Kern 2016; Mo and Siepel 2023) We therefore experimented with training the AFT method on simulated data with a constant population size history and then testing the classifier on data simulated under two nonequilibrium models of population contraction followed later by either instantaneous or exponential expansion (Marth *et al*. 2004; Gutenkunst *et al*. 2009), and vice-versa. The AFT method was able to accurately detect sweeps even in the extreme scenario of unmodeled population size change, but selection coefficient estimation accuracy suffered in this scenario. More work testing on additional demographic histories would be required to fully understand how our method and other time-series methods would perform in various scenarios of demographic model misspecification. Another important, albeit intuitive result of our analyses is that AFT generally performs better on smaller samples obtained across a larger number of timepoints than on datasets consisting of the same total number of individuals sampled from a smaller number of timepoints. This implies that researchers interested in detecting adaptation from longitudinal collections may wish to prioritize sampling individuals across a wider range of times, even if the number of individuals collected at each timepoint is fairly small—five individuals per timepoint may be sufficient if many timepoints are sampled.

The relative failure of our haplotype-based method may contain lessons for future improvements. The goal of this approach was to contain all of the information present in a Muller plot (e.g. Figure 2 from Herron and Doebeli 2013), which shows the frequency trajectories of each haplotype present in a (typically clonal) population sampled at regular intervals. Although this information is very useful for revealing whether a haplotype may have been favored by selection, it does not contain any information about the degree of dissimilarity between distinct haplotypes, nor does it contain any information about the locations of any variants that may be present on multiple haplotypes. The latter shortcoming implies that this approach should have little power to distinguish between the targets of selection and closely linked loci, and this is supported by our findings shown in Figures S3-4. Future work to develop haplotype-frequency tracking representations that contain information both about the location of polymorphisms differing among haplotypes, and the degree of dissimilarity between haplotypes sharing a given allele, could yield improvements in detecting and localizing selected polymorphisms. Such advancements could also aid in distinguishing between SDN and SSV models of sweeps—a problem that both our HFT and AFT methods struggled with, and for which information about the frequencies of different haplotypes bearing the adaptive allele is paramount (Garud *et al*. 2015).

Our AFT method, on the other hand, was highly successful at localizing sweeps. This is perhaps because this approach makes an inference for a single focal polymorphism while taking into account spatial information about the presence or absence of nearby hitchhiking variants, before sliding on to the next focal polymorphism. Although this method does not explicitly use haplotype information, it may implicitly be able to track LD among polymorphisms by seeing correlated shifts in allele frequencies (see Figure1). Thus, it is possible that little information is lost by relying on allele frequency trajectories rather than haplotype frequencies.

An added benefit of the AFT approach is that it can be used with unphased data, or even pooled sequencing data. This allowed us to test the method on data from an evolve-and-resequence study that repeatedly used pool-seq to sample experimental populations of *D. simulans* over a period of 60 generations of selection to thermal tolerance (Barghi *et al*. 2019). We found that Timesweeper was able to detect regions with alleles that experienced large shifts in frequency over the course of this selection experiment. Moreover, Timesweeper often detected the same putatively sweeping alleles in multiple experimental replicates, suggesting that its results are accurate. We stress that these encouraging results were obtained in spite of no adjustment of our method to these data: the founder lines of the *D. simulans* experiment exhibit a very high density of polymorphism, and thus our approach to examine 51-SNP windows caused us to examine only a relatively small genetic map distance when making each classification—incorporating a larger flanking window in this setting would likely yield even better performance. We believe that the successful application of Timesweeper to these data in spite of this limitation bodes well for the broad applicability and robustness of our approach.

In light of Timesweeper’s strong performance on simulated and real genomic data, we have released the software package implementing it to the public. We strove to make this package easy to use and computationally efficient, as we believe that fast, user-friendly, and powerful computational tools are essential if we wish to realize the potential of population genomic time-series data. We hope that further research in this area of population genetic methods development, combined with the application of these methods to time-series data from natural populations, will help us better characterize the genomic targets and evolutionary importance of positive selection. For example, the continued development and analysis of these data sets can help us resolve the controversy over the impact of positive selection in nature (Hahn 2008; Kern and Hahn 2018; Jensen *et al*. 2019). Such work may also play an important role in the genomic surveillance of both organisms relevant to human health (e.g. vectors and pathogens) as well as populations threatened by climate change or other anthropogenic disruptions.

### Data and code availability

Timesweeper’s source code and documentation can be found at https://github.com/SchriderLab/Timesweeper, it is available to be built from source as well as installed as a pip package. All scripts and workflows can be found at https://github.com/SchriderLab/timesweeper-experiments to allow for a complete reconstruction of all data/figures used in this manuscript. Data from the *D. simulans* evolve-and-resequence dataset were obtained from Dryad (https://doi.org/10.5061/dryad.rr137kn) as described in the Methods.

## Acknowledgments

We thank three reviewers and Amjad Dabi for comments on the manuscript.

## Funding

This work was funded by NIH awards R35GM138286 and R01AI153523.

## SUPPLEMENTARY INFORMATION LEGENDS

Figure S1: E**x**ample **and average Timesweeper inputs for AFT and HFT formats.** The top two rows are individual replicates randomly selected for AFT (left column) and HFT (right column). The bottom two rows are averaged inputs across 10,000 replicates for each class: the third row is zoomed in to show the vicinity of the central polymorphism in closer detail for the AFT method, as well as the HFT method’s input zoomed in to focus on the haplotype with the highest frequency increase and those most similar two it; the bottom row shows the entirety of these inputs. For the scenarios with selection (i.e. SSV and SDN) initial sampling time was drawn from a uniform distribution with bounds [-50, 50) and selection coefficient was drawn from a uniform distribution with bounds [0.00025, 0.25). 10 diploid individuals were sampled every 10 generations for 200 generations (see Methods).

Figure S2: N**e**ural **network architecture diagrams.** Full diagrams with layer types and dimensions for the (A) 1DCNN, (B) Fully Connected Network (FCN), (C) 2DCNN, (D) 1DCNN with an increased number of parameters, and (E) RNN.

Figure S3: R**e**solution **of the AFT and HFT methods assessed on simulated 500 kb chromosomes using varying window sizes.** The fraction of polymorphisms classified as a sweep at varying locations along the 500 kb chromosome binned into 501 adjacent windows. When a sweep is present (SSV and SDN columns), it occurs in the center of the chromosome. The three left columns show the classification results of the AFT method, the right three columns show the classification results of the HFT method, the window size of the feature vector increases from the top to bottom row.

Figure S4: R**e**solution **of the AFT and HFT methods assessed on central 501 polymorphisms using varying window sizes.** The fraction of polymorphisms classified as a sweep at each polymorphism along the centralmost 501 polymorphisms. When a sweep is present (SSV and SDN columns), it occurs in the center of the window. The three left columns show the classification results of the AFT method, the right three columns show the classification results of the HFT method, and the window size of the feature vector increases from the top to bottom row.

Figure S5: T**i**mesweeper’s **classification performance on datasets of varying selection coefficient.** Timesweeper was trained and tested on 10,000 replicates of each scenario (neutrality, SSV, SDN) with a sampling generation drawn from a uniform distribution with bounds [-50, 50) for selection coefficients of 0.005, 0.01, 0.05, 0.1, and 0.5. Precision-recall (PR) curve, ROC curve, Normalized and non-normalized confusion matrices, and training losses for AFT (left half) and HFT (right half) for *s* values of 0.005, 0.01, 0.05, 0.1, and 0.5.

Figure S6: R**O**C **Curves for varying selection coefficient classification**. ROC curves for varying selection coefficients as described in Figure S5 for classifying sweeps versus neutrality (top row) and the SDN scenario versus the SSV scenario (bottom row) for both AFT (left column) and HFT (right column).

Figure S7: P**R Curves for varying selection coefficient classification.** PR curves for varying selection coefficients as described in Figure S5 for classifying sweeps versus neutrality (top row) and the SDN scenario versus the SSV scenario (bottom row) for both AFT (left column) and HFT (right column).

Figure S8: T**i**mesweeper’s **classification performance using different neural network architectures.** Timesweeper was trained and tested on the dataset described in Figure S1 using different neural network architectures. PR curves, ROC curves, normalized and non-normalized confusion matrices, and training losses for the classification task (neutrality versus sweep, SDN versus SSV classification) for 1DCNN (top row), 1DCNN with increased parameterization (second row), 2DCNN (third row), and RNN (bottom row) for both AFT (left half) and HFT (right half).

Figure S9: T**i**mesweeper’s **selection coefficient-inference performance using different neural network architectures.** Timesweeper was benchmarked on a 20-timepoint dataset with selection coefficients drawn from a uniform distribution with bounds [0.00025, 0.25) and a starting sampoint generation drawn from a uniform distribution with bounds [-50, 50) post-selection onset. True *s* versus estimated *s values* are plotted for both SDN and SSV scenarios with training losses for each network for 1DCNN (top row), 1DCNN with increased parameterization (second row), 2DCNN (third row), and RNN (bottom row) for both AFT (left half) and HFT (right half).

Figure S10: S**a**liency **maps of a trained 2DCNN implementation of Timesweeper.** Saliency maps for the 2DCNN architecture trained on the same benchmark data as described in Figure S8, saliency was calculated using examples from the neutral (left column), SSV (central column), and SDN scenarios (right column) for (A) AFT and (B) HFT data formats.

Figure S11: T**i**mesweeper’s **classification performance on a benchmark dataset using different window sizes.** Timesweeper was trained and tested on the dataset described in Figure S1 using window sizes of 1, 3, 11, 51, 101, and 201 SNPs. Precision Recall (PR) curve, ROC curve, normalized and non-normalized confusion matrices, and training losses for AFT (left half) and HFT (right half) are displayed for each window size.

Figure S12: ROC curves for classification performance on datasets with various window sizes. Same as Figure S6 but for the data described in Figure S11.

Figure S13: PR curves for classification performance on datasets with various window sizes. Same as Figure S7 but for the data described in Figure S11.

Figure S14: Timesweeper’s regression performance on datasets with various window sizes. Same as Figure S9 but for the data described in Figure S11.

Figure S15: T**i**mesweeper’s **classification performance on datasets with various sample sizes.** For all simulations 20 timepoints were taken using the same selection coefficient and sampling time parameterization as described in Figure S1. Sample sizes of 1, 2, 5, 10, and 20 individuals were taken at each timepoint. Precision Recall (PR) curve, ROC curve, Normalized and non-normalized confusion matrices, and training losses for AFT (left half) and HFT (right half) are displayed for each sample size.

Figure S16: ROC curves for classification performance on datasets with various sample sizes. Same as Figure S6 but for the data described in Figure S15.

Figure S17: PR curves for classification performance on datsets with various sample sizes. Same as Figure S7 but for the data described in Figure S15.

Figure S18: Timesweeper’s regression performance on datasets with various sample sizes. Same as Figure S9 but for the data described in Figure S15.

Figure S19: T**i**mesweeper’s **classification performance on datasets with various numbers of sampled timepoints.** For all simulations a total of 200 diploid individuals were sampled evenly across the specified number of timepoints (i.e., 200 for the 1-timepoint scenario, 100 per timepoint for the 2-timepoint scenario, etc). Simulation parameterizations for selection coefficient and sampling start time are the same as described in Figure S1. Precision-recall (PR) curve, ROC curve, normalized and non-normalized confusion matrices, and training losses for AFT (left half) and HFT (right half) are displayed for 1, 2, 5, 10, 20, and 40 timepoints.

Figure S20: ROC curves for classification performance on datasets with various numbers of sampled timepoints. Same as Figure S6 but for the data described in Figure S19.

Figure S21: PR curves for classification performance on datasets with various numbers of sampled timepoints. Same as Figure S7 but for the data described in Figure S19.

Figure S22: Timesweeper’s regression performance on datasets with various numbers of sampled timepoints. Same as Figure S9 but for the data described in Figure S19.

Figure S23: T**i**mesweeper’s **classification performance on datasets with various sampling start times**. Simulation parameterizations for selection coefficient are the same as described in Figure S1. Sampling start time was −100, −50, 0, 25, 100, or 200 generations post-onset of selection. Precision Recall (PR) curve, ROC curve, normalized and non-normalized confusion matrices, and training losses for AFT (left half) and HFT (right half) are displayed.

Figure S24: ROC curves for classification performance on datasets with various sampling start times. Same as Figure S6 but for the data described in Figure S23.

Figure S25: PR curves for classification performance on datasets with various sampling start times. Same as Figure S7 but for the data described in Figure S23.

Figure S26: Timesweeper’s regression performance on datasets with various sampling start times. Same as Figure S9 but for the data described in Figure S23.

Figure S27: T**i**mesweeper’s **classification performance under various training set sizes.** Simulation parameterizations for selection coefficient and sampling start times are the same as described in Figure S1. Data was subsampled to sizes 30,000, 20,000, 10,000, 5,000, 2,000, and 1,000 for each class (neutral, SSV, SDN). Each dataset was partitioned into training, validation, and test data as described in the Methods. Precision Recall (PR) curve, ROC curve, normalized and non-normalized confusion matrices, and training losses for AFT (left half) and HFT (right half) are displayed.

Figure S28: ROC curves for classification performance under various training set sizes. Same as Figure S6 but for the data described in Figure S27.

Figure S29: PR curves for classification performance under various training set sizes. Same as Figure S7 but for the data described in Figure S27.

Figure S30: Timesweeper’s regression performance on datasets with varyious training set sizes. Same as Figure S9 but for the data described in Figure S27.

Figure S31: Proportion of positive sweep calls for Timesweeper and competing methods, zoomed in towards the center of the window. The same as Figure S4 but for the data described in Figure 5.

Figure S32: M**i**sspecification **classification ROC and PR curves.** Pairwise ROC (left) and PR (right) curves for neutral versus sweep (top row) and SDN versus SSV (middle row) tasks. The legend specifies the demographic models used for training and testing (in that order) for each curve. FET’s ROC (bottom left) and PR (bottom right) curves are shown for test data simulated under the Bottleneck and OoA models.

Figure S33: M**i**sspecification **regression accuracies.** Pairwise plots of true versus estimated *s* for each model trained (rows) and tested on (columns) for both SSV (left) and SDN (right) scenario selection coefficients.

Figure S34: Example and average Timesweeper inputs from simulated training data for the *D. simulans* evolve-and-resequence dataset. (A) – (C) show individual example inputs and (D) shows average inputs of the Neutral and SSV classes for the AFT method where data were simulated for the *D. simulans* evolve-and-resequence dataset as described in the Methods.

Figure S35: T**i**mesweeper’s **classification performance on simulated test data for the *D. simulans* evolve-and-resequence dataset.** (A) – (D) Confusion matrix, ROC curve, training trajectory, and precision-recall (PR) curve showing the performance of the AFT method on simulated *D. simulans* data as described in the Methods.

Figure S36: R**e**plication **of top hits for Timesweeper and FET in the *D. simulans* evolve-and- resequence dataset.** A datapoint was counted as replicated sweep if it A) was in the top 100 most confidently-detected sweeps (ranked by Timesweeper sweep probability or by FET 1-pvalue) and B) it occurred in a 100kb window containing at least one outlier SNP (i.e. top 1% of all tested SNPs for a replicate) detected in the other replicate.

Table S1: Summaries of Timesweeper’s classification results on D. simulans evolve-and- resequence dataset binned by Fisher’s Exact Test p-value.

Table S2: **Summaries of Fisher’s Exact Test results on *D. simulans* evolve-and-resequence dataset binned by Timesweeper’s sweep probability score.**

## Notes

### Competing Interest Statement

The authors have declared no competing interest.

### Summary of Updates

We have substantially revised the manuscript and added a number of new analyses. Most notably, we have expanded Timesweeper to estimate selection coefficients, and we now compare its performance to a larger set of previously published methods.

