## Supplementary figures and images for "Timesweeper: Accurately Identifying Selective Sweeps Using Population Genomic Time Series"

### A9Rn4dpui_yk6b6r_jhc.tmp

SSV Estimated vs True s for Misspecification

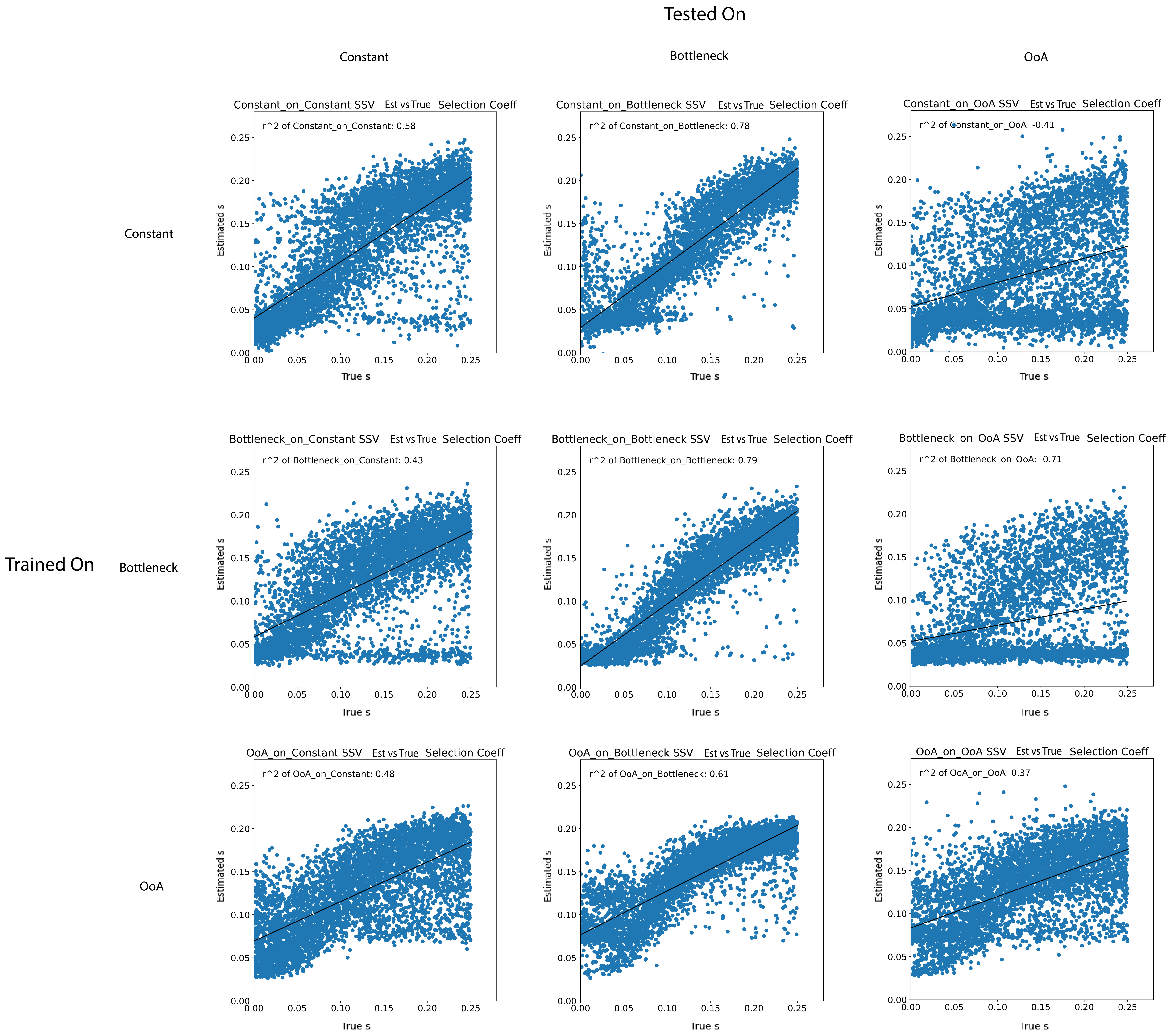


SDN Estimated vs True s for Misspecification

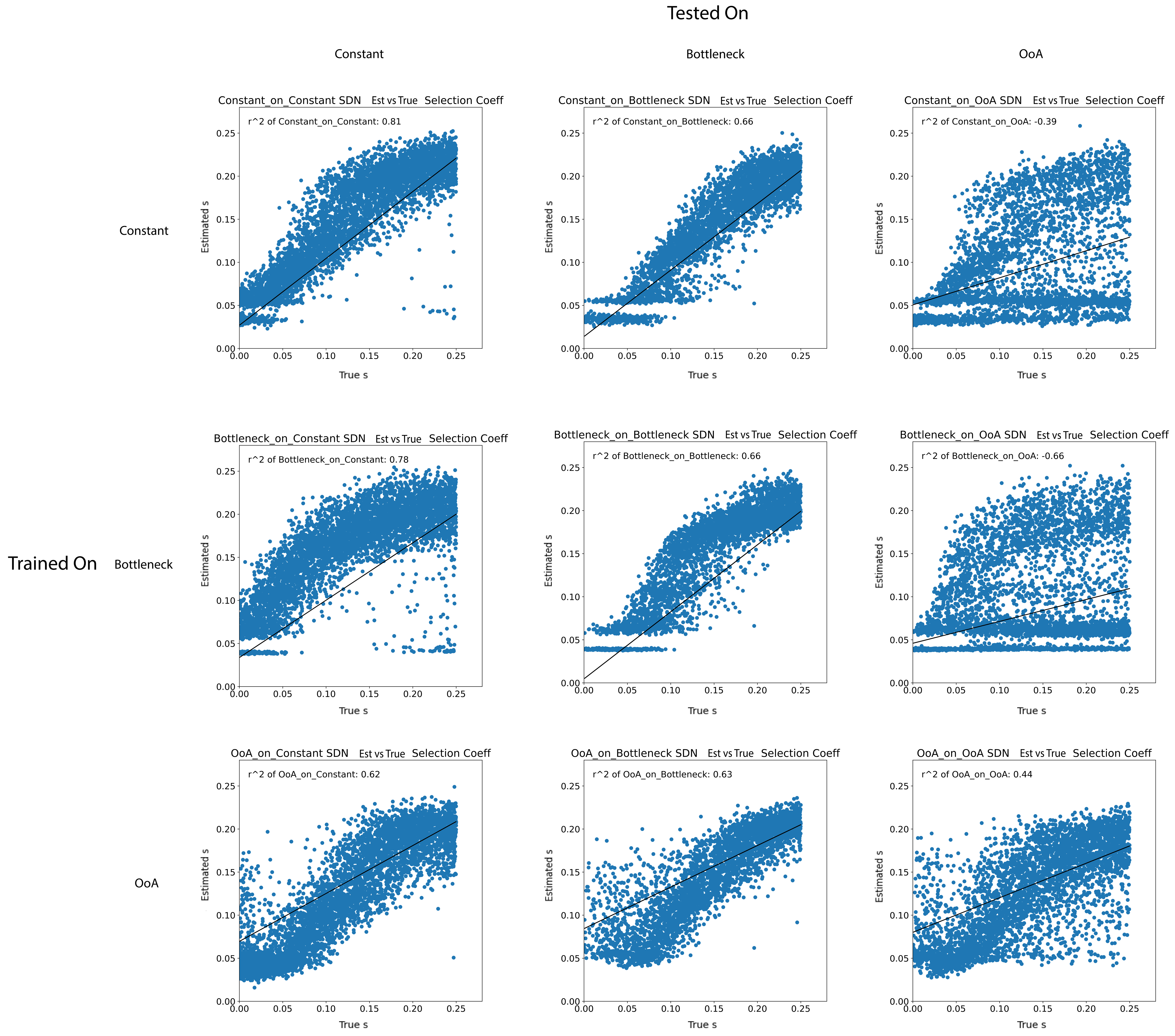

### S1_All_Inputs.pdf

#### Input Representations for Timesweeper

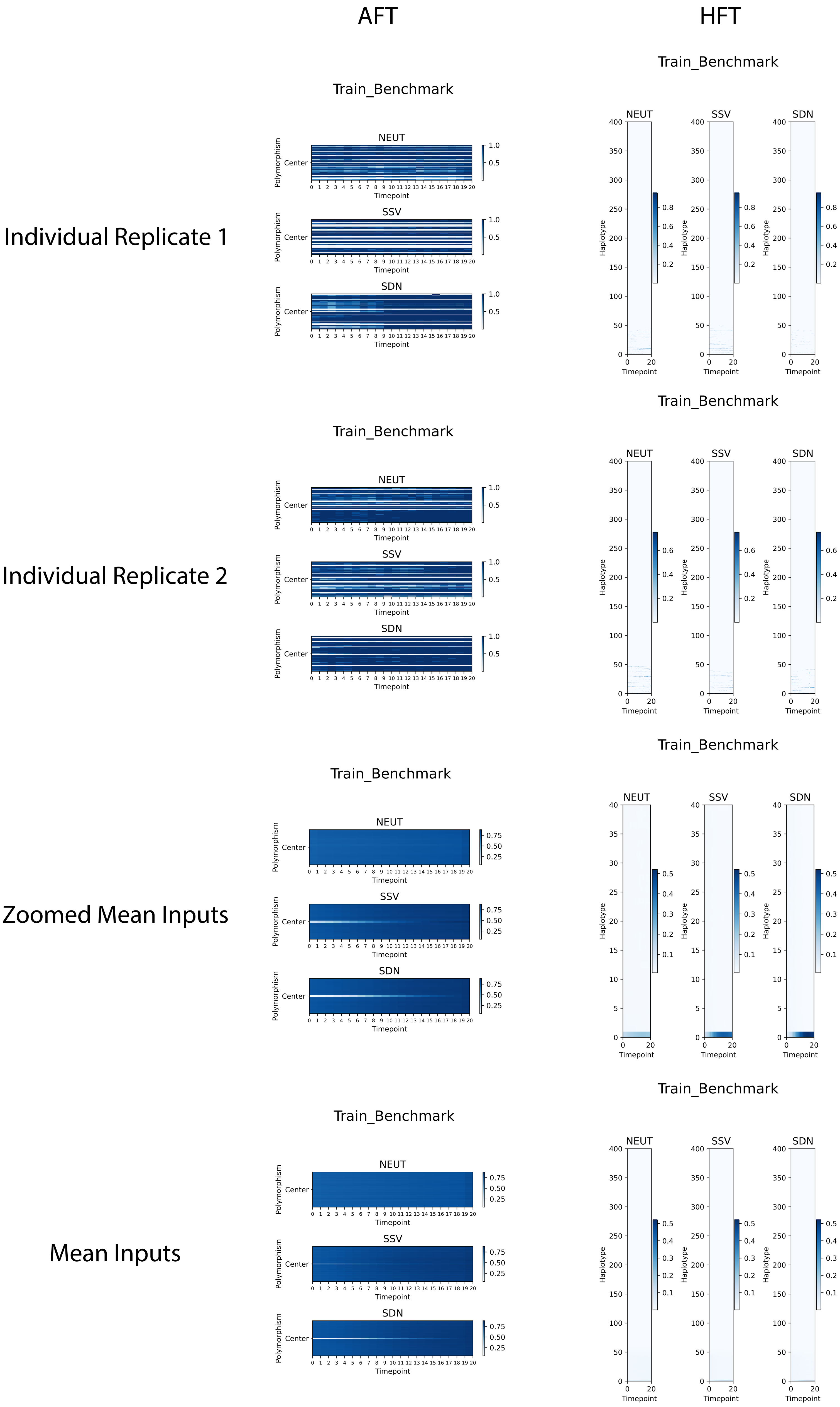

### S2_Architectures.png

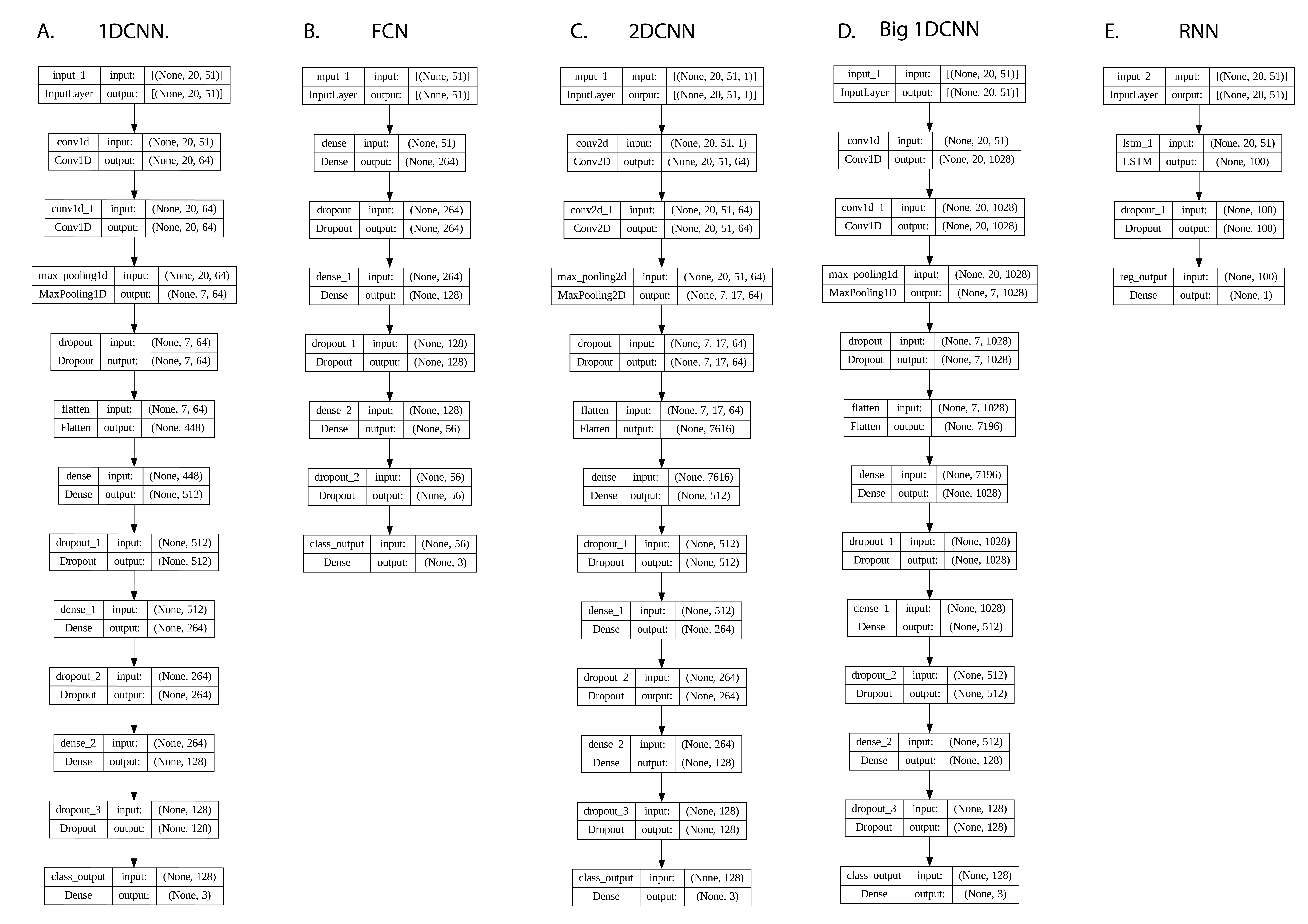

### S3_Unzoomed_Spikes.pdf

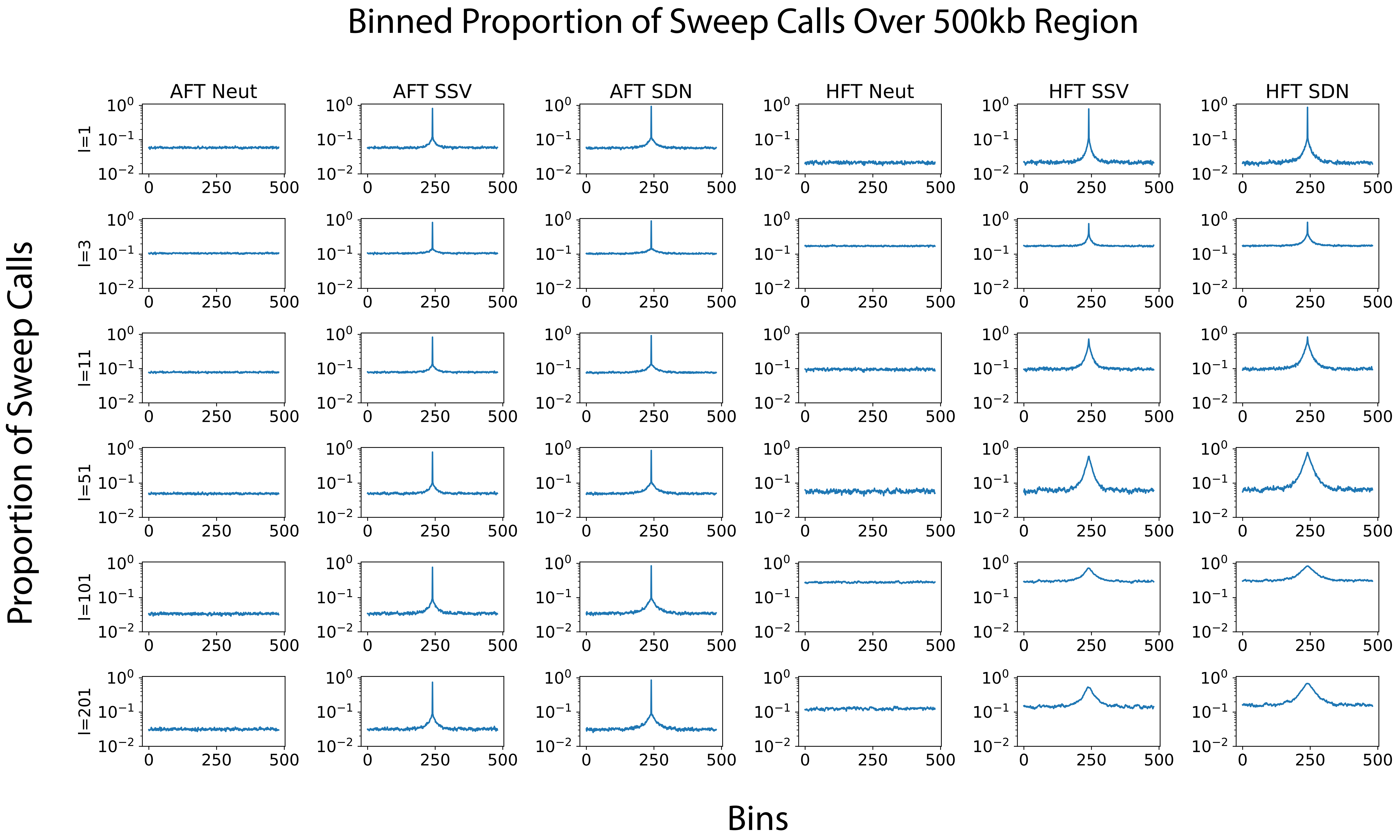

### S4_Zoomed_Spikes.pdf

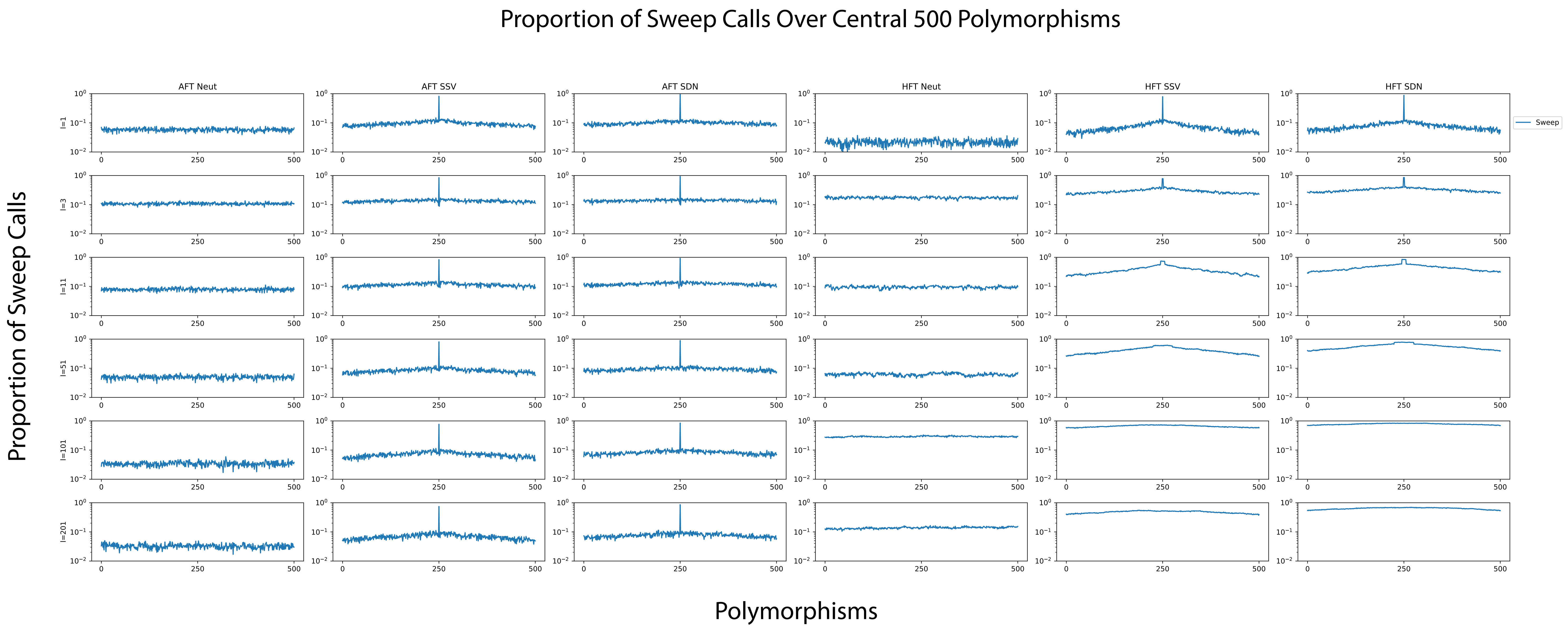

### S5_Sel_Coeff_Class.pdf

#### Classification - Selection Coefficient

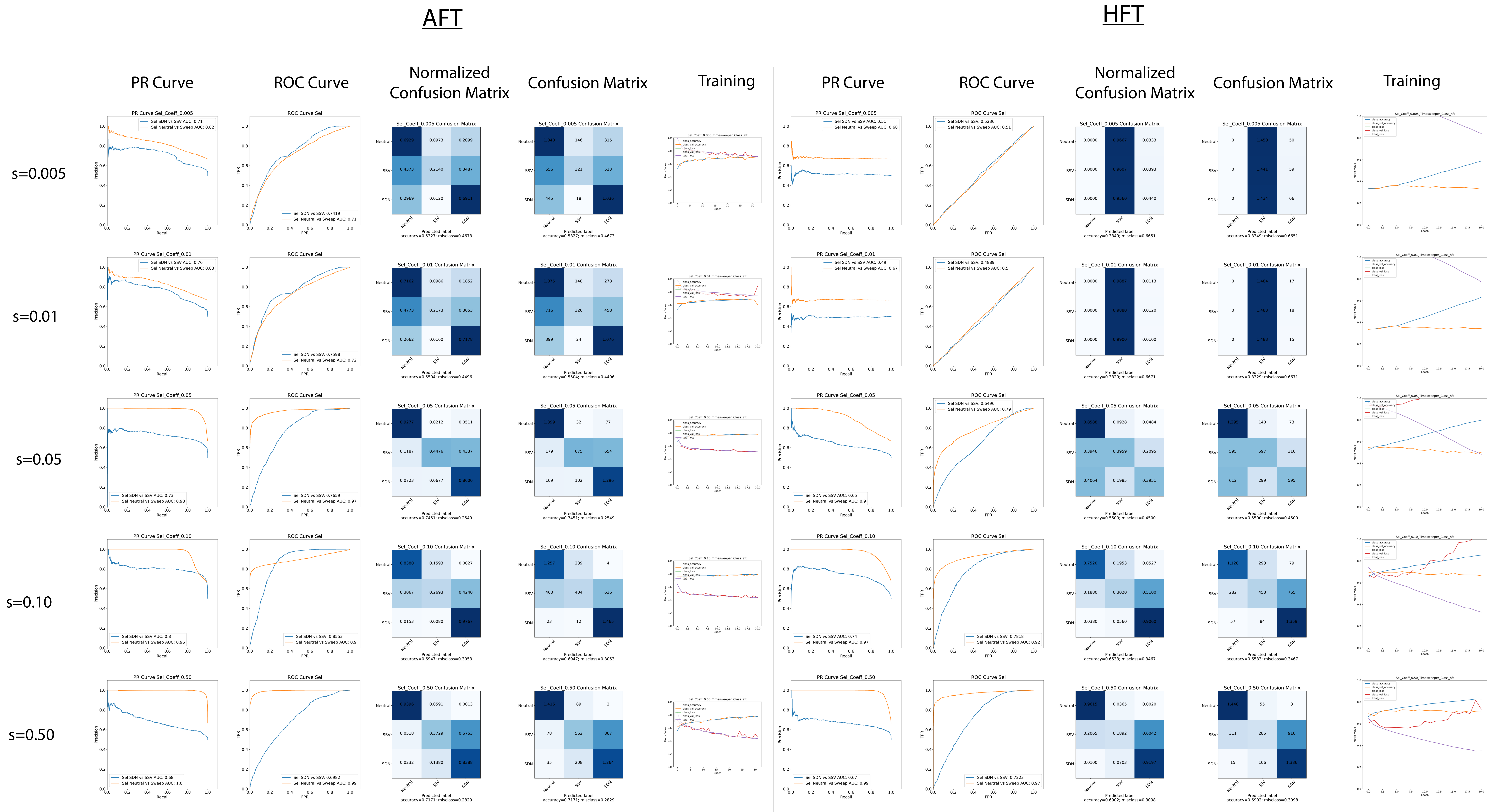

### S6_Sel_Coeff_ROC.pdf

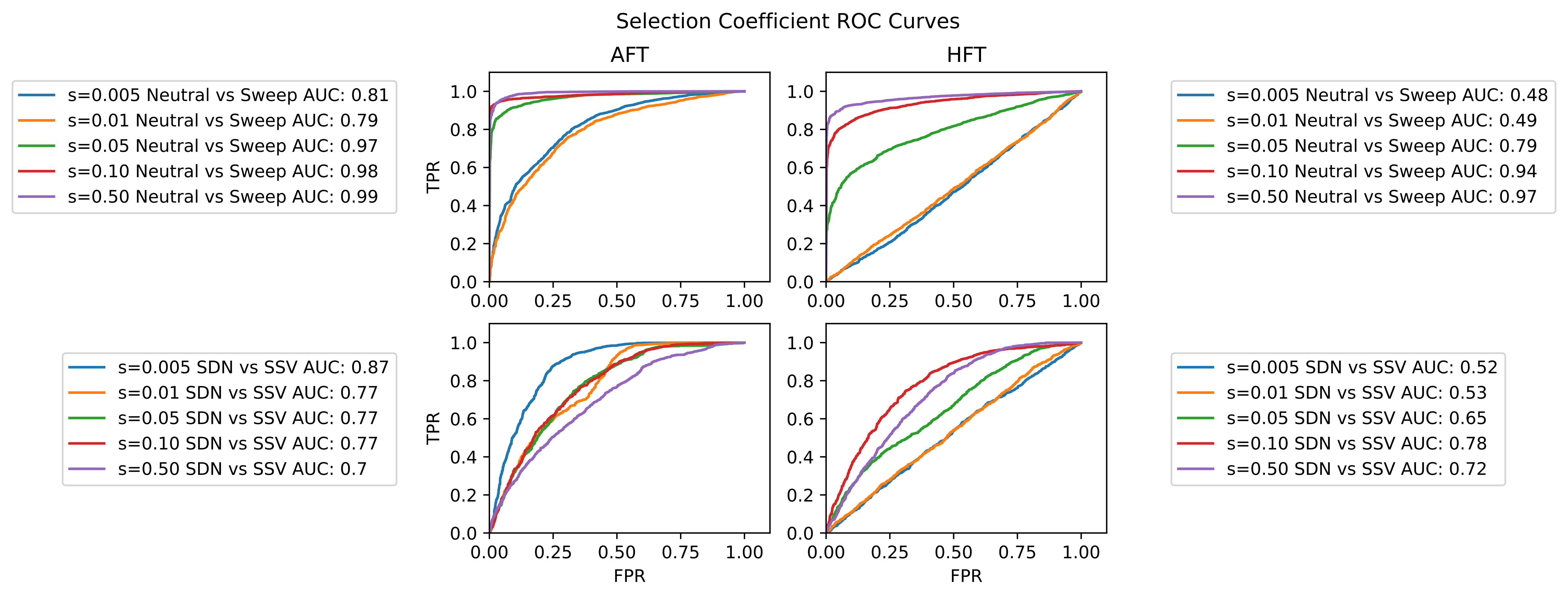

### S7_Sel_Coeff_PR.pdf

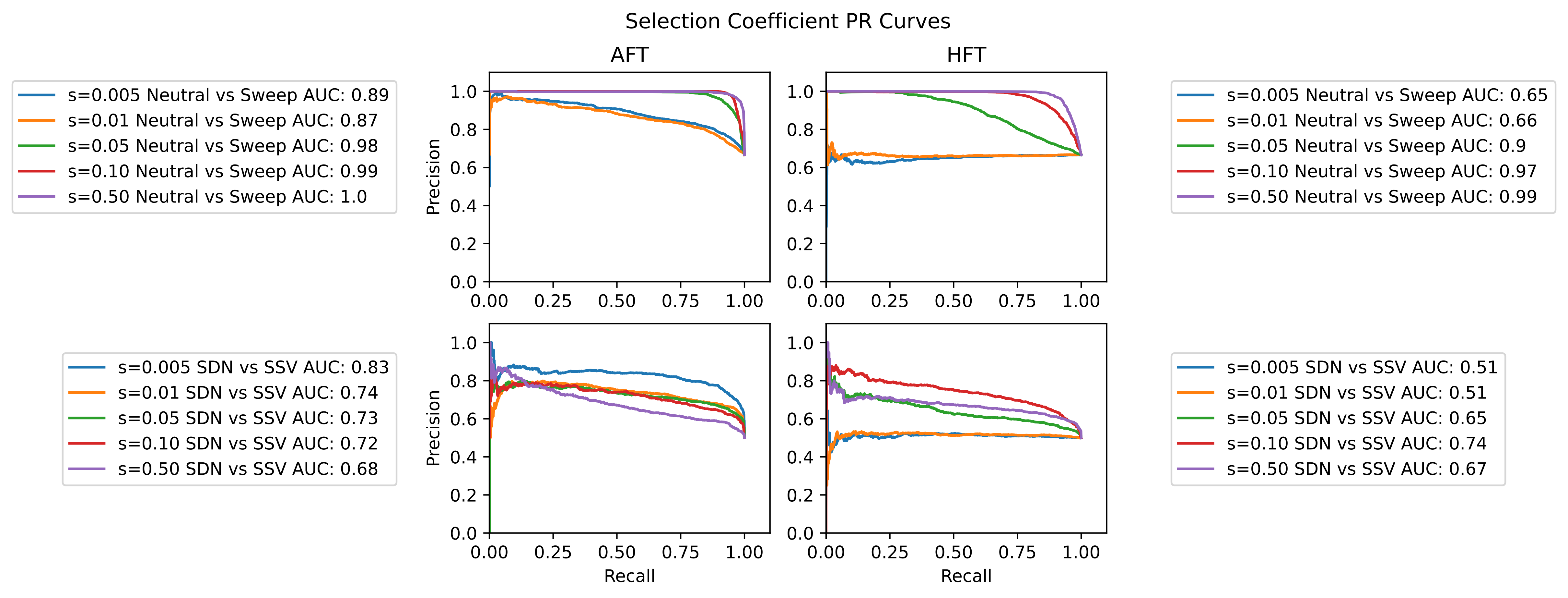

### S8_Architectures_Classification.pdf

#### Classification - Architectures

##### AFT

##### HFT

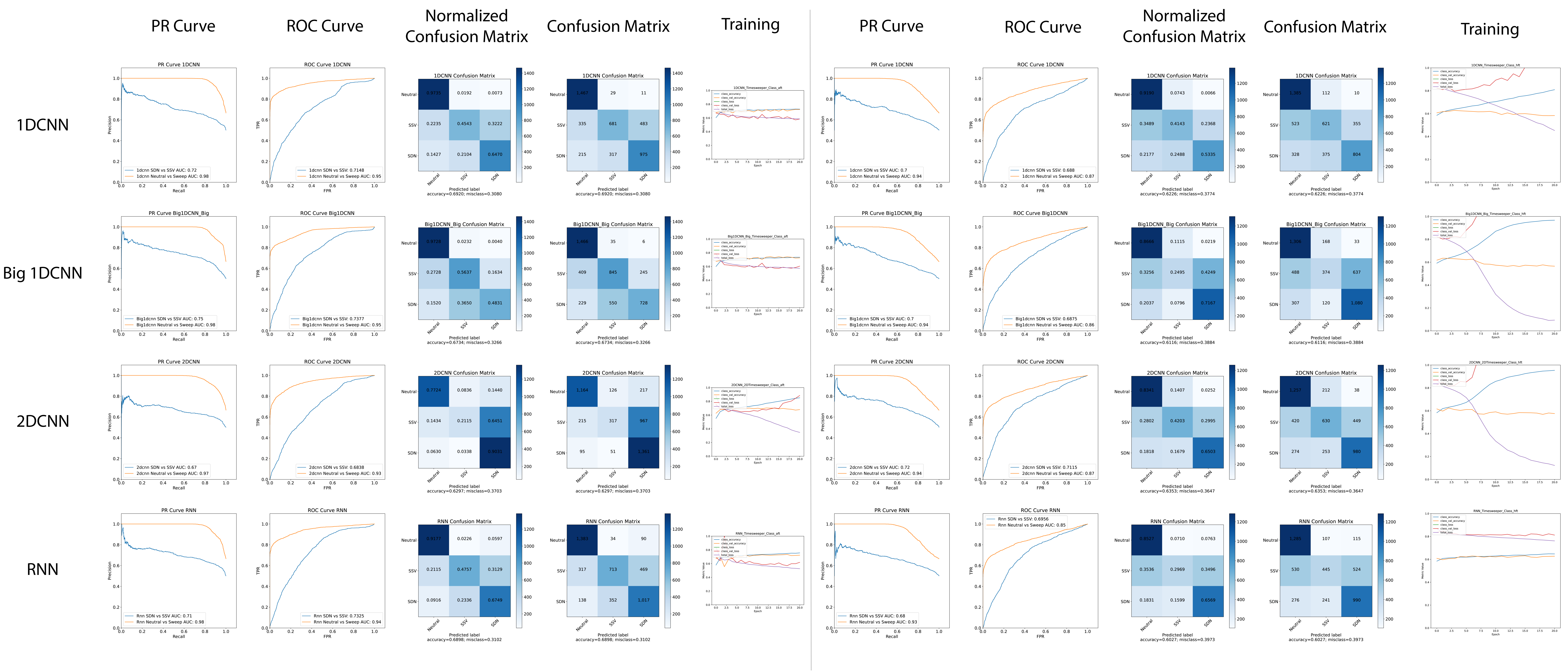

### S9_Architectures_Regression.pdf

#### Regression - Architectures

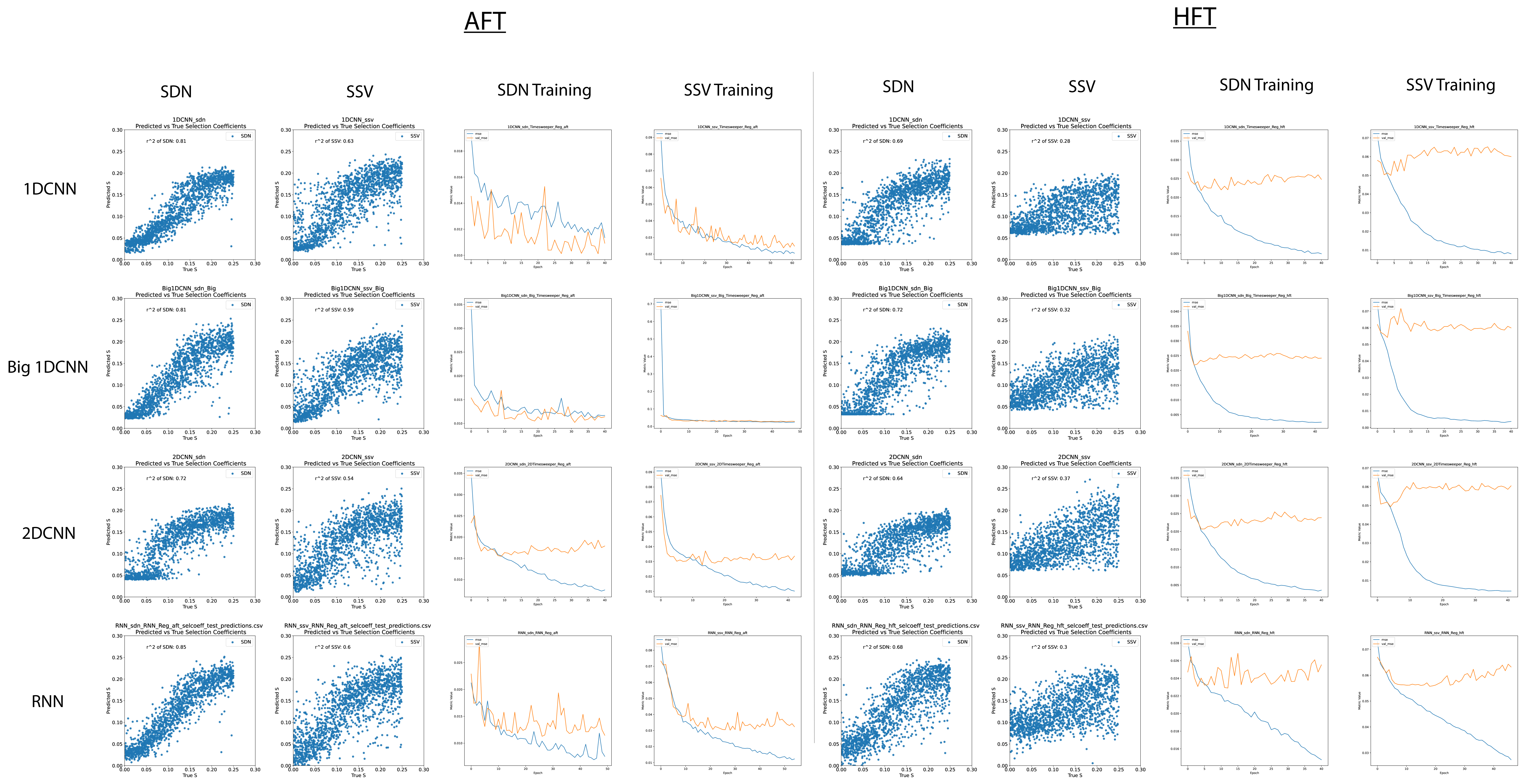

### S10_Saliency_maps.pdf

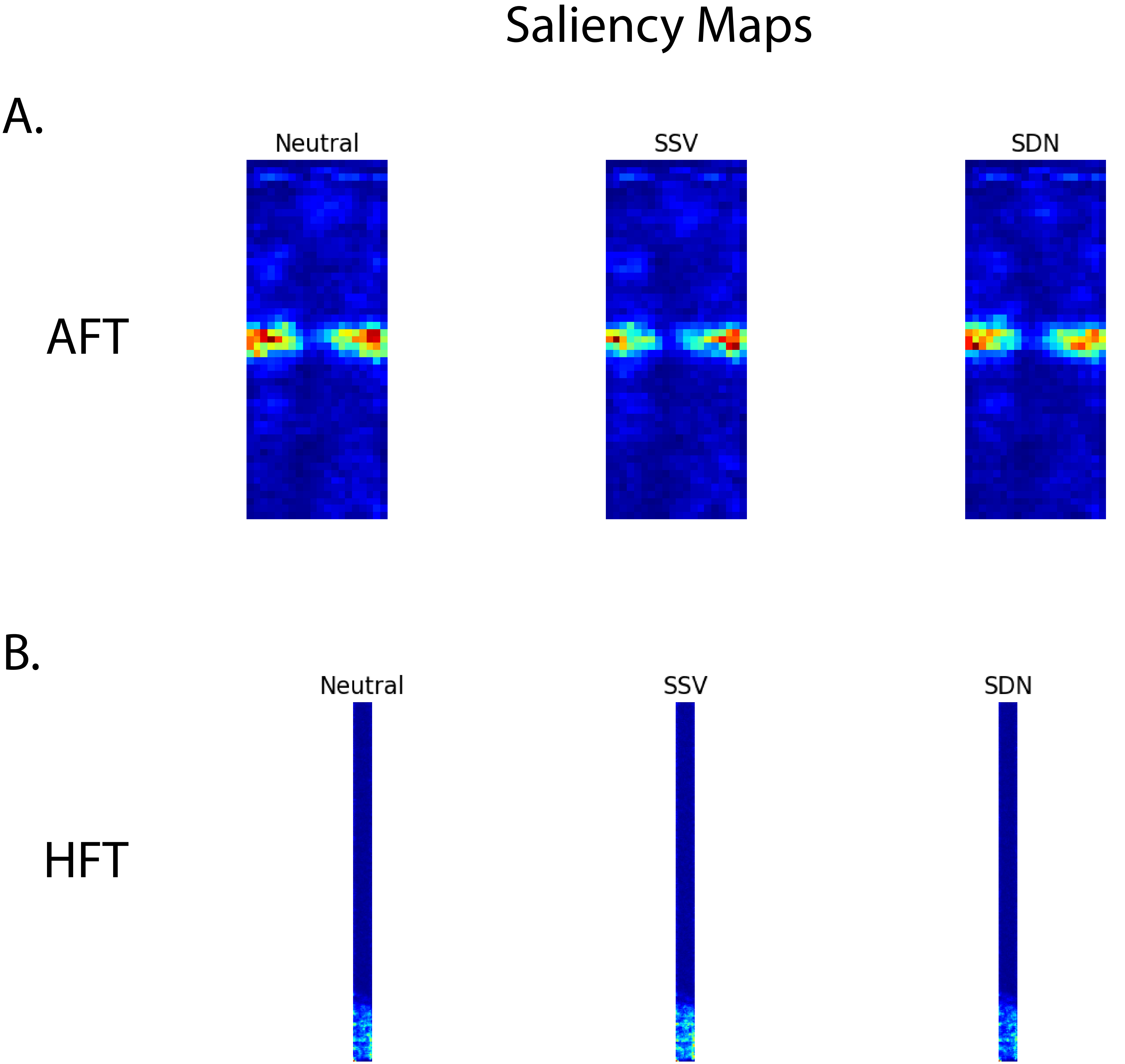

### S11_Win_Sizes_Class.pdf

#### Classification - Selection Timing

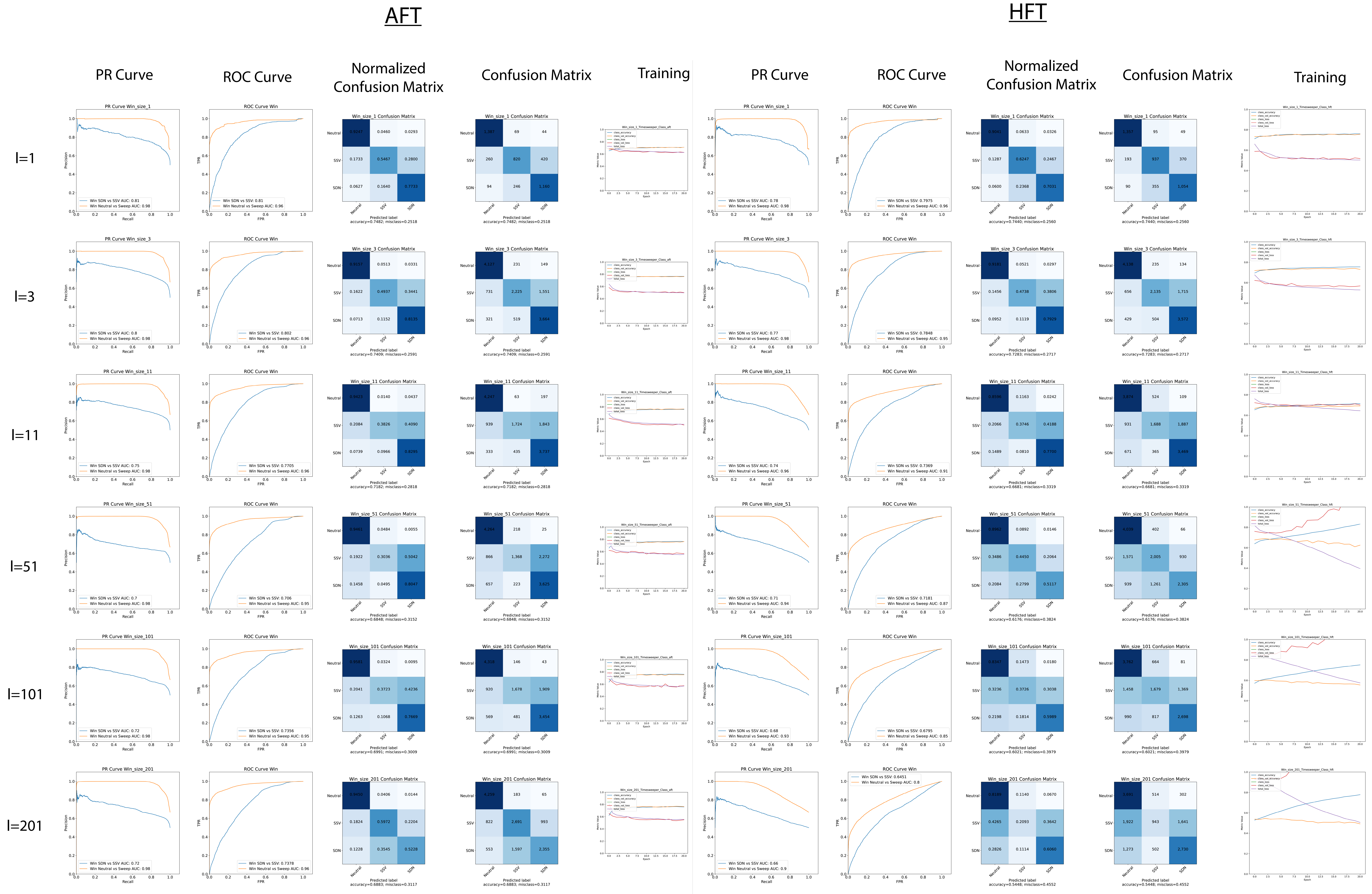

### S12_Win_Sizes_ROC.pdf

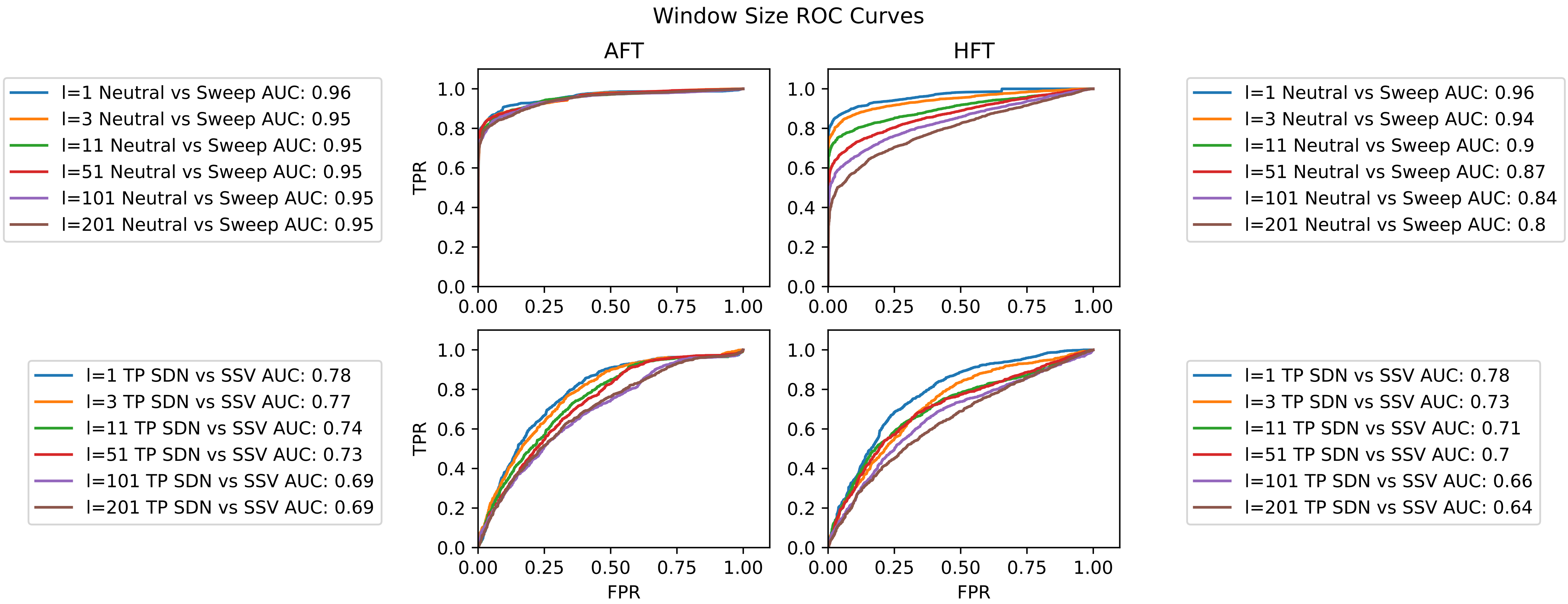

### S13_Win_Sizes_PR.pdf

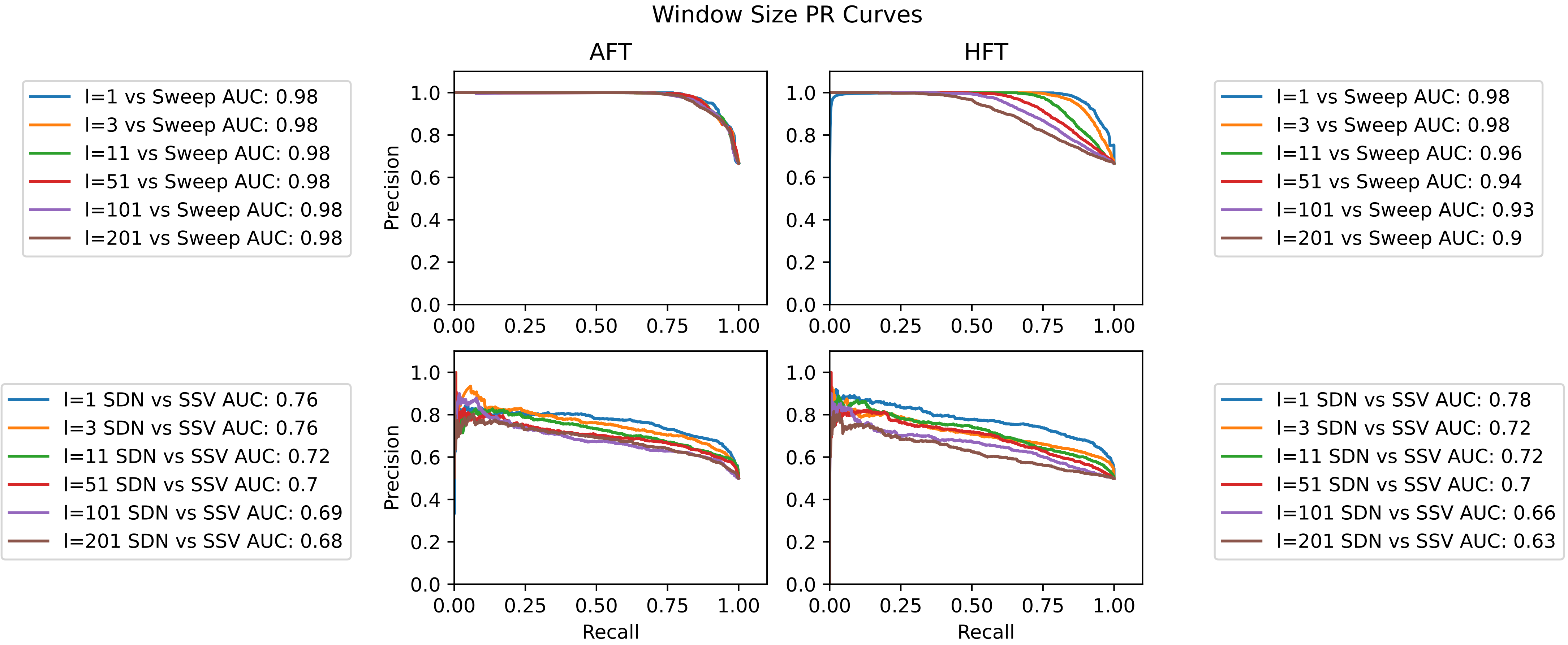

### S14_Win_Sizes_Reg.pdf

#### Regression - Window Sizes

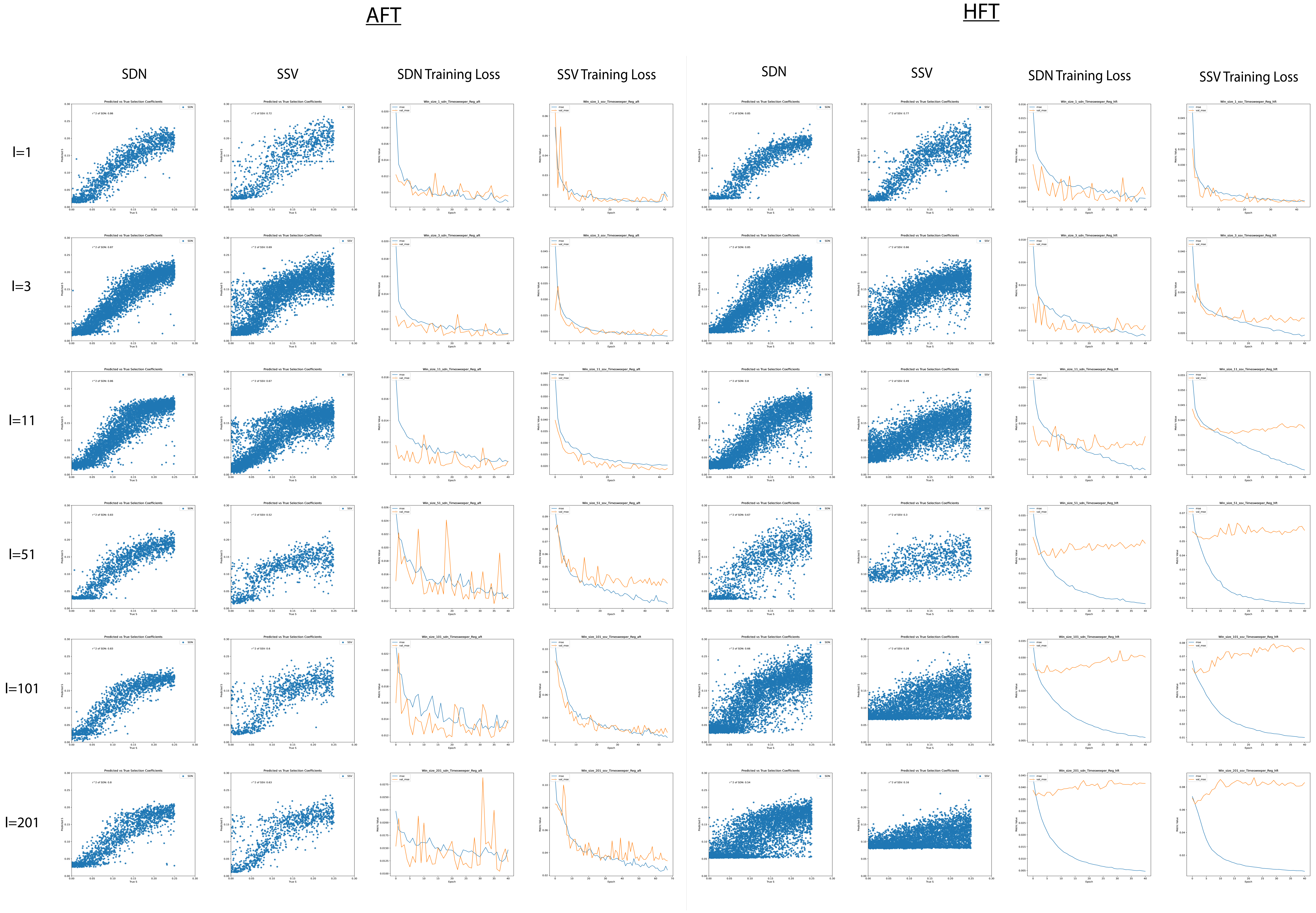

### S15_Samp_Size_Class.pdf

#### Classification - Sample Size

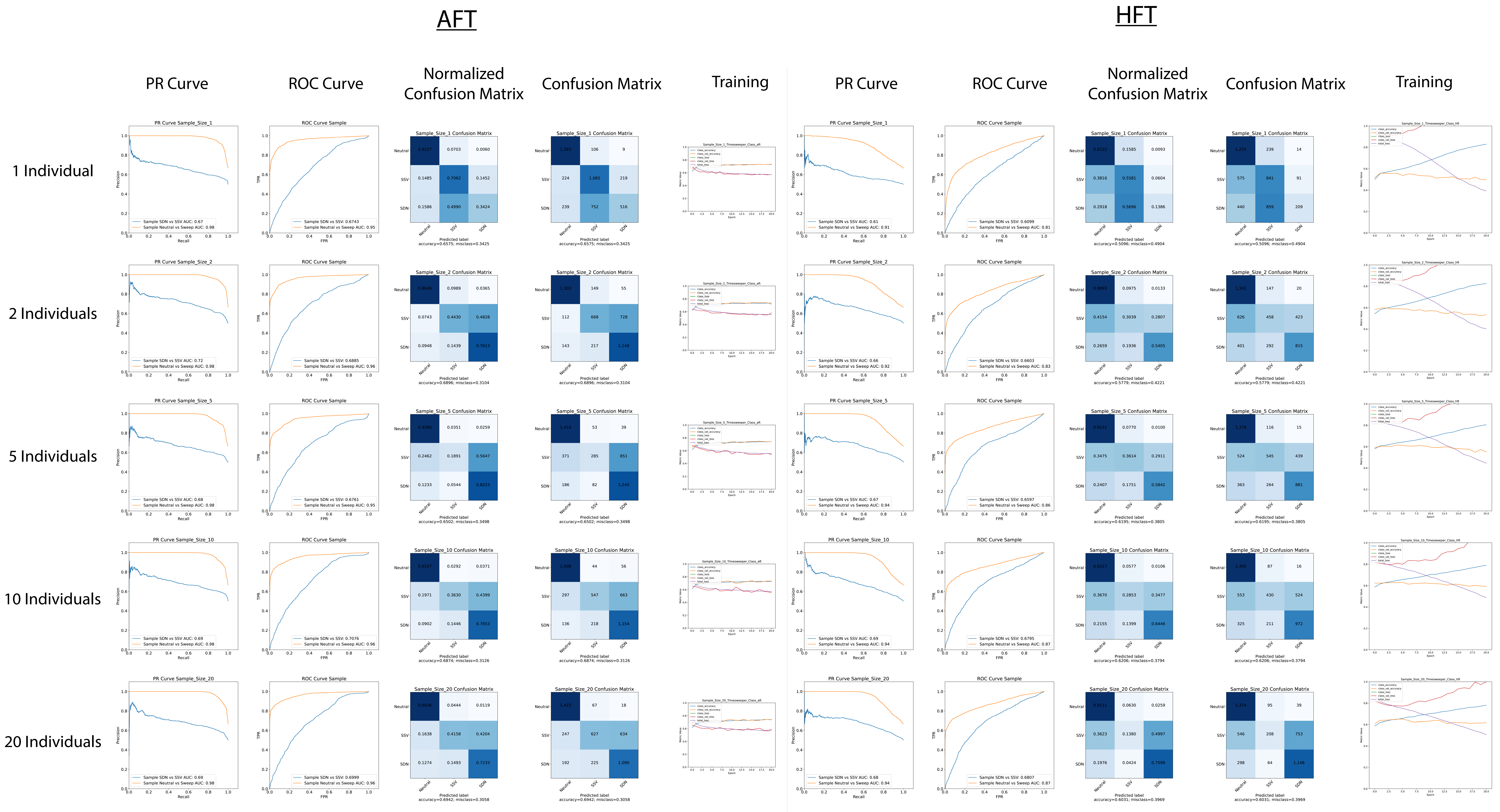

### S16_Samp_Size_ROC.pdf

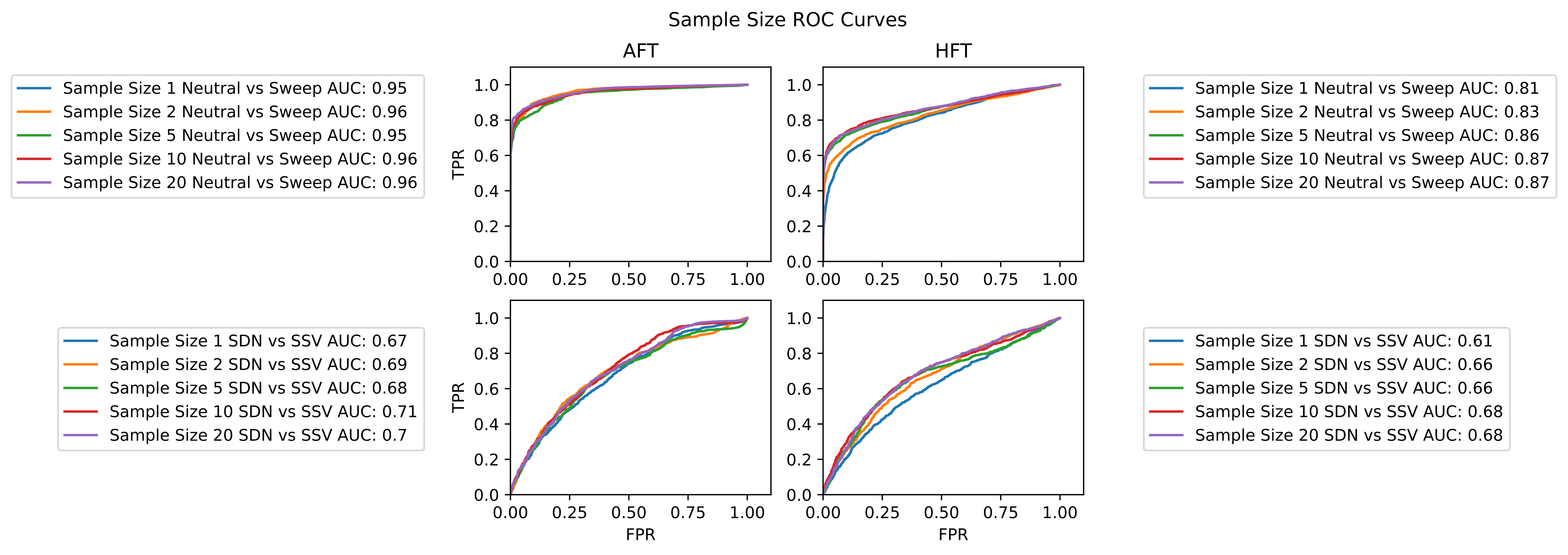

### S17_Samp_Size_PR.pdf

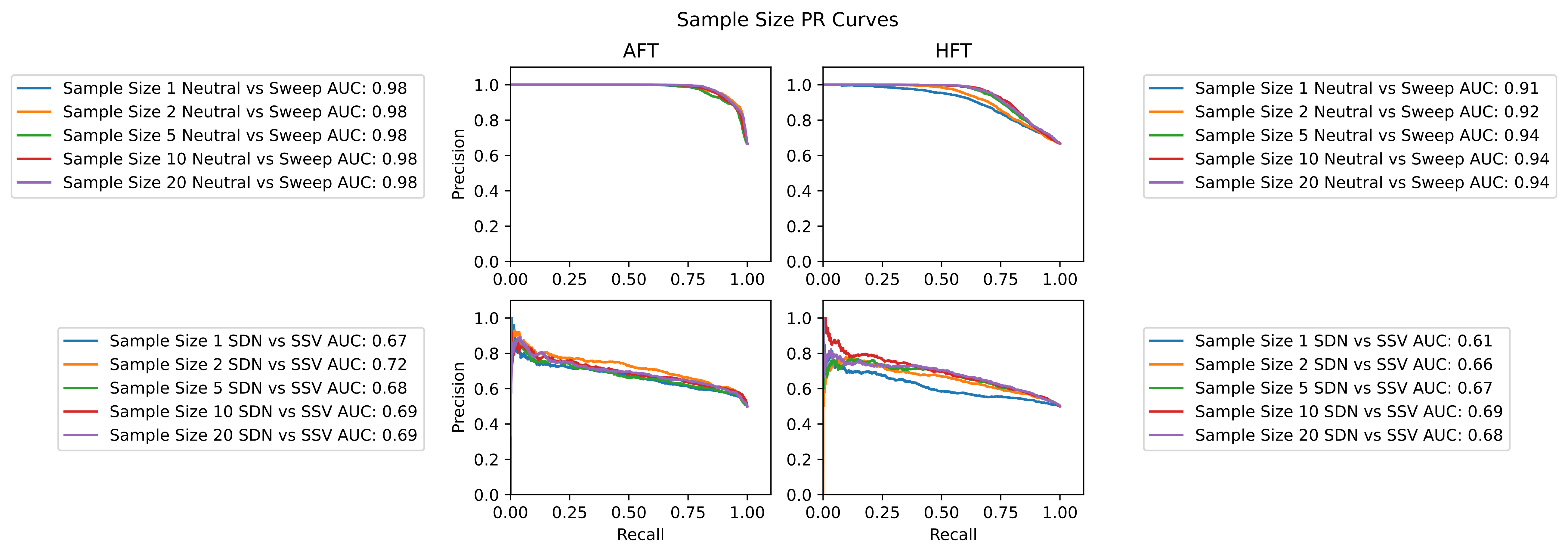

### S18_Samp_Size_Reg.pdf

#### Regression - Sample Size

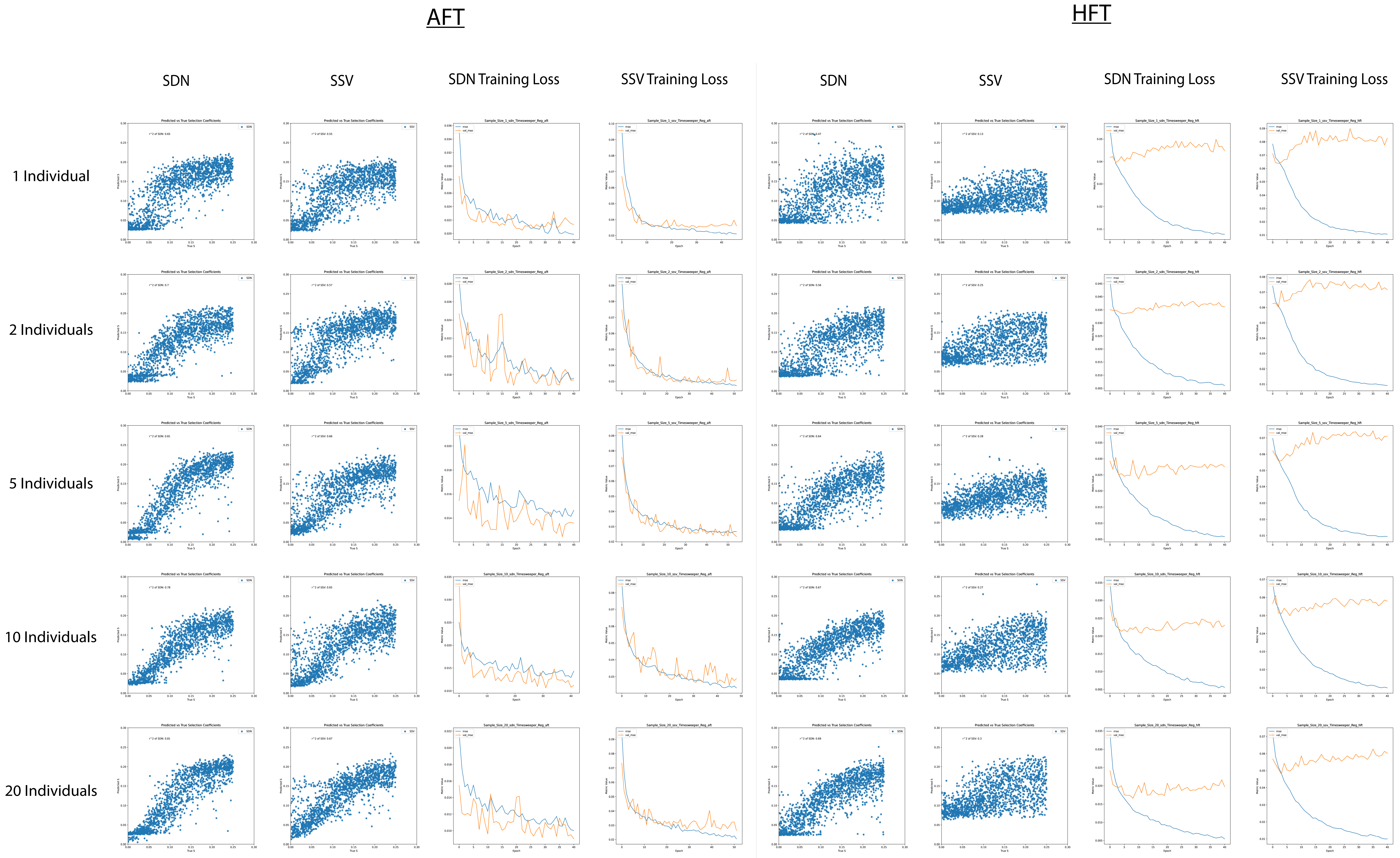

### S19_TPs_Vary_Class.pdf

#### Classification - Varying Timepoints

##### AFT

##### HFT

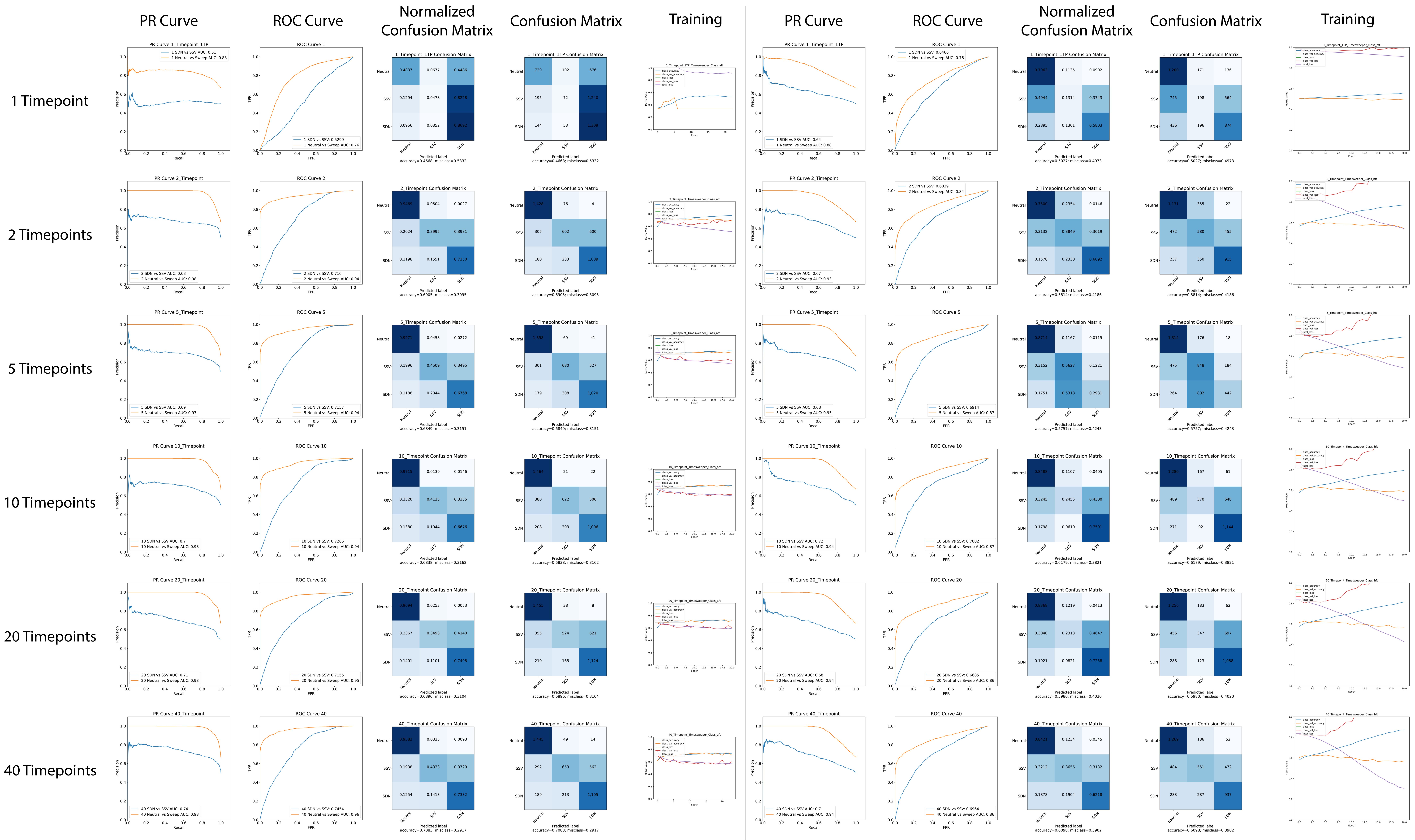

### S20_TPs_Vary_ROC.pdf

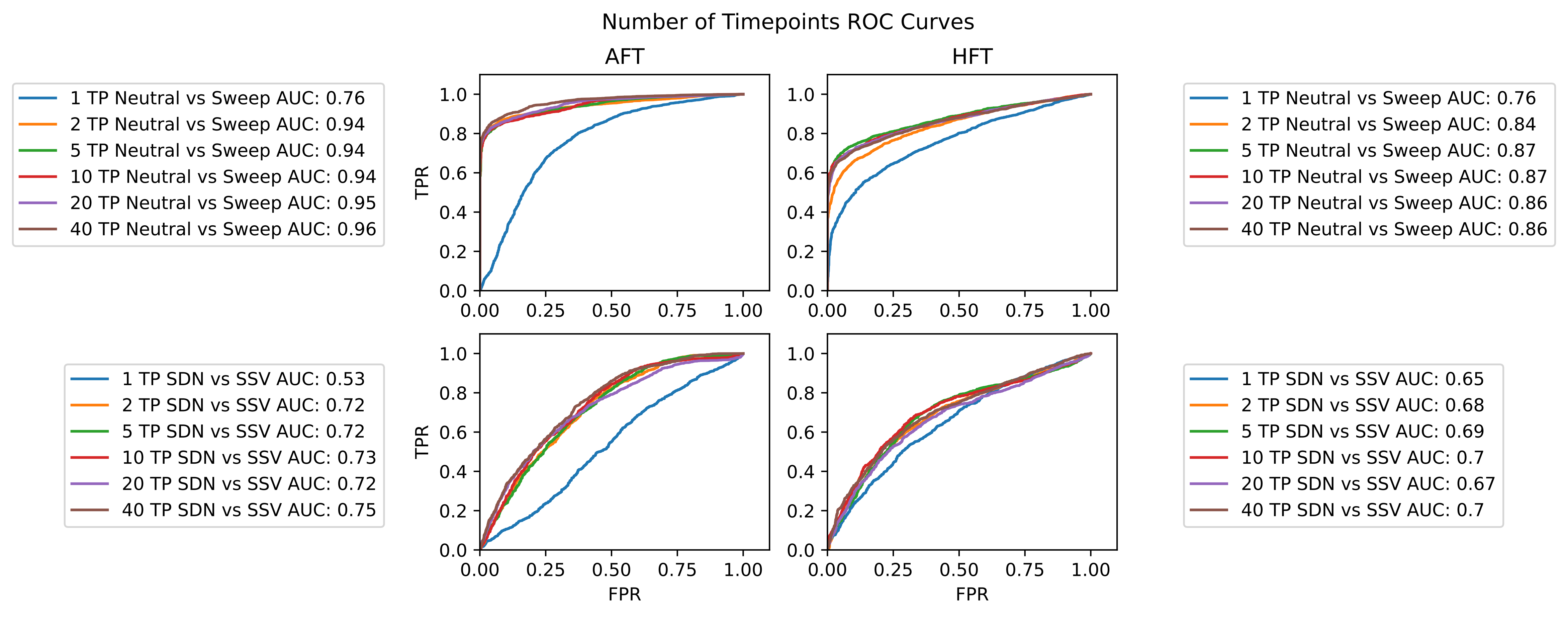

### S21_TPs_Vary_PR.pdf

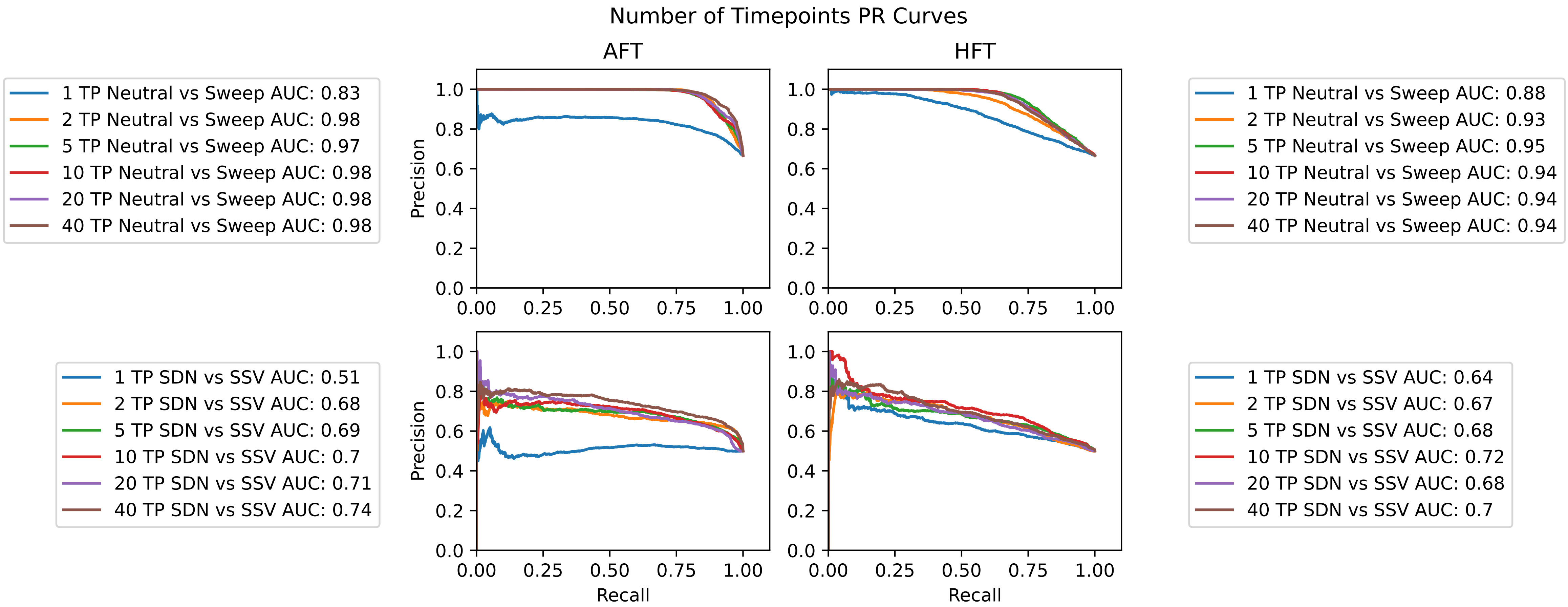

### S22_TPs_Vary_Reg.pdf

#### Regression - Varying Timepoints

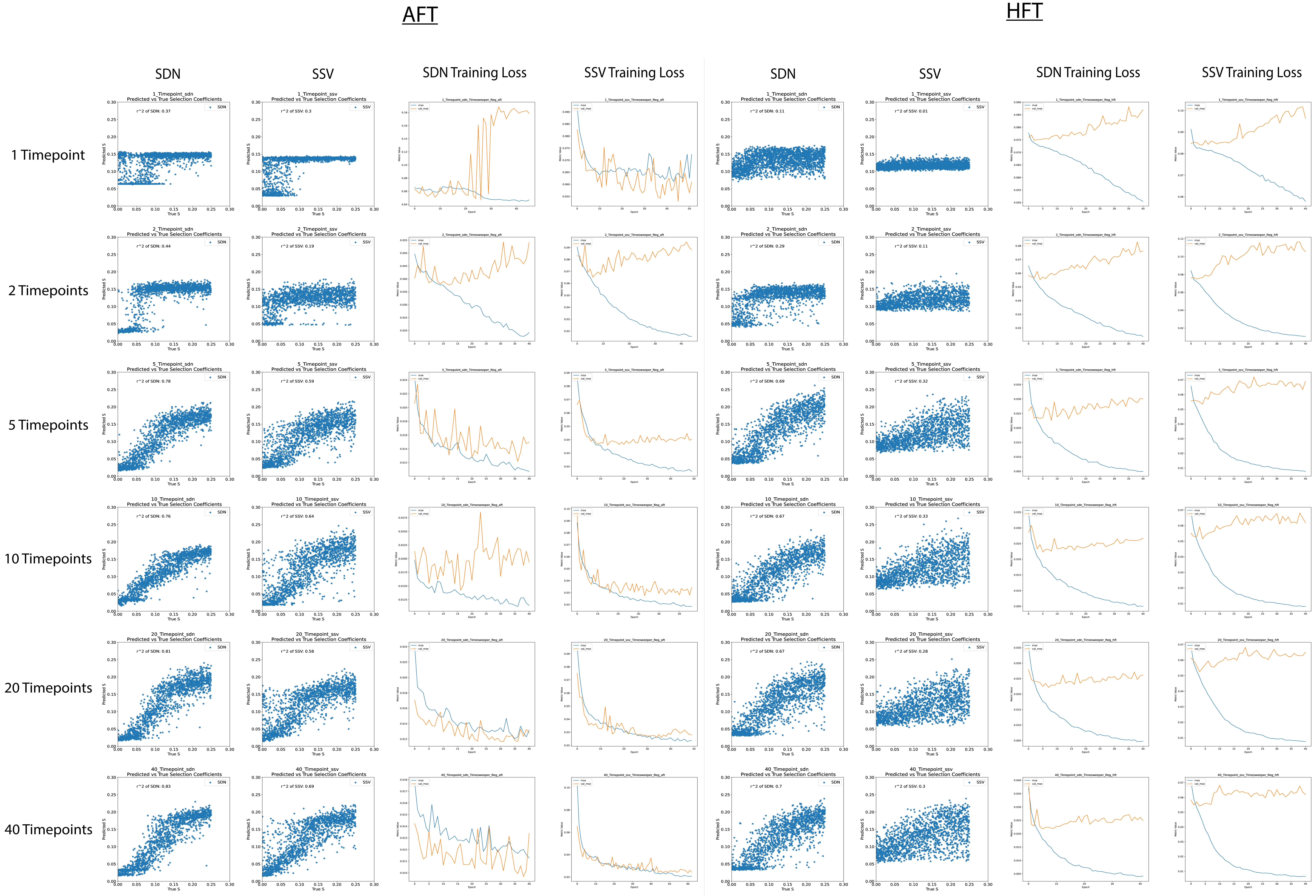

### S23_Sel_Timing_Class.pdf

#### Classification - Selection Timing

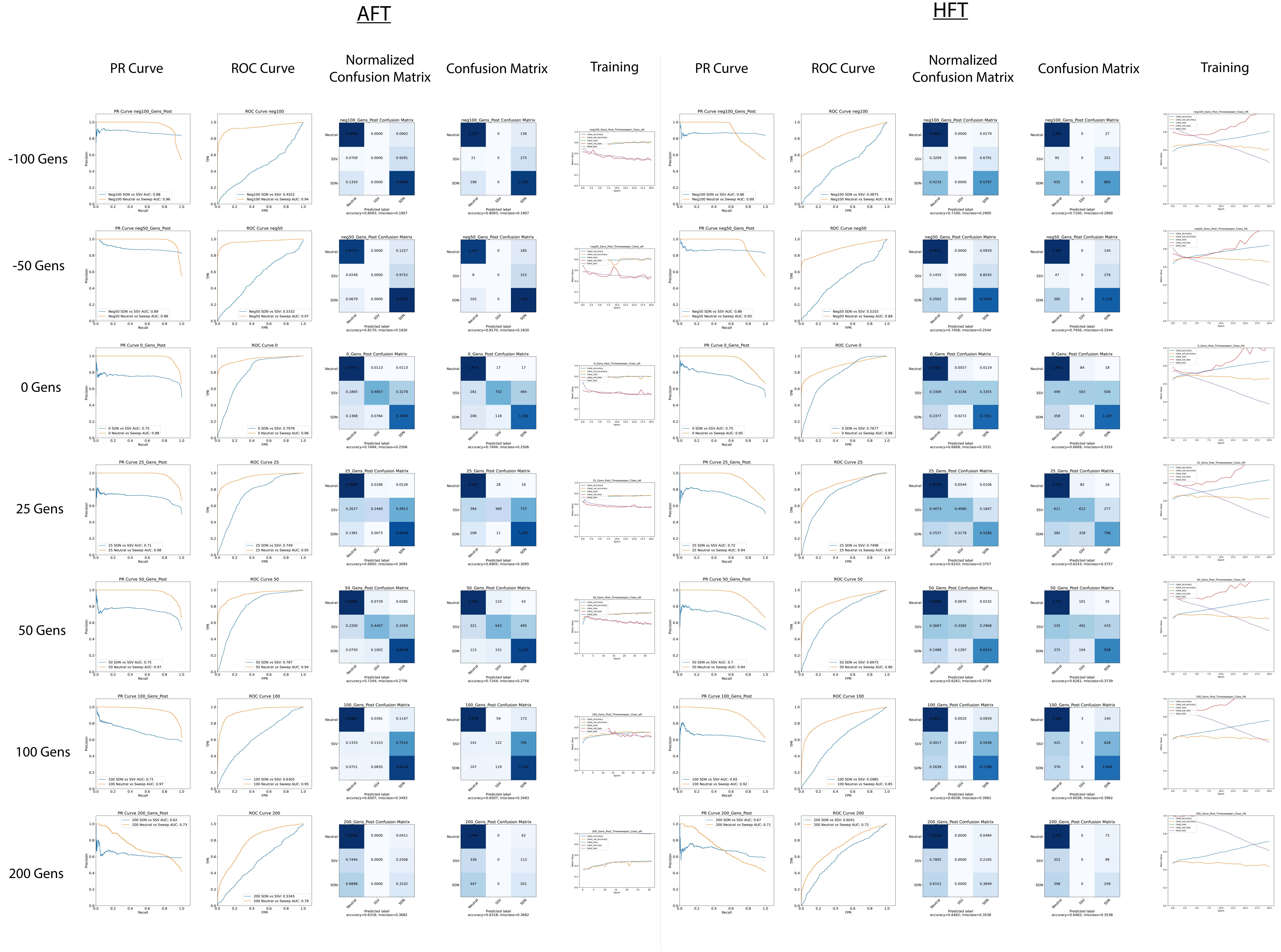

### S24_Sel_Timing_ROC.pdf

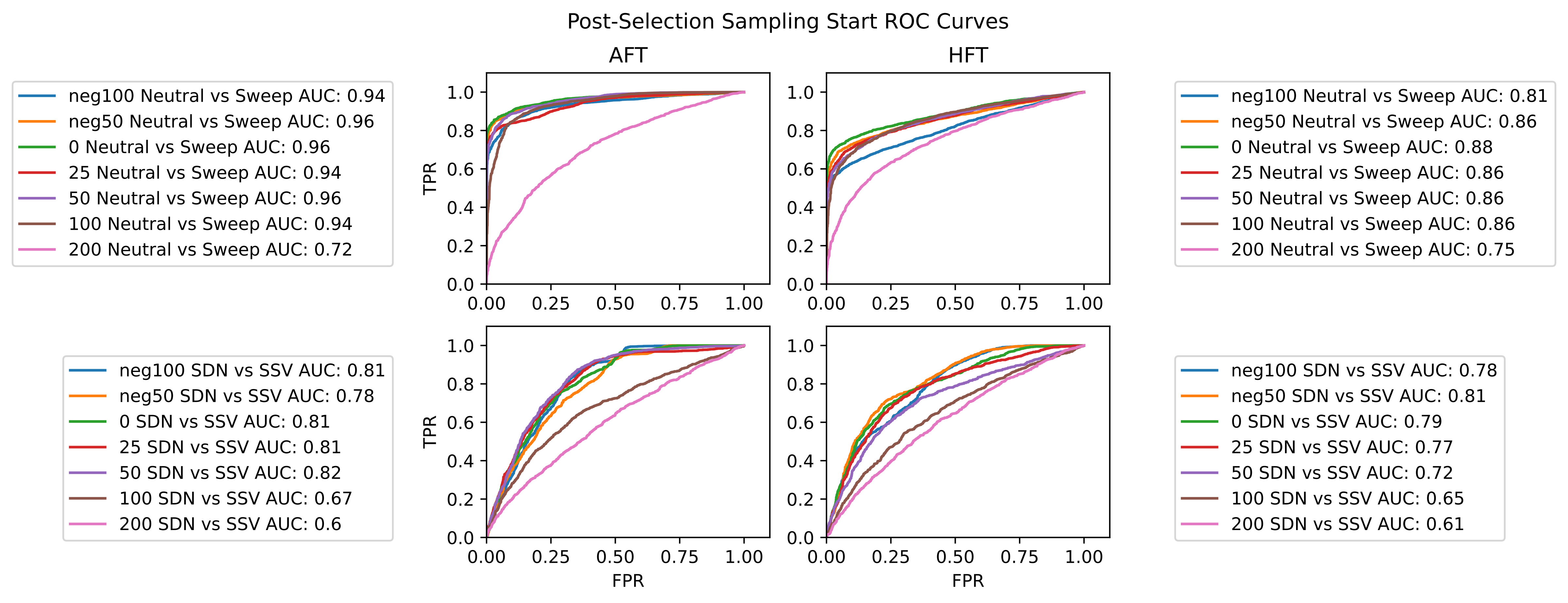

### S25_Sel_Timing_PR.pdf

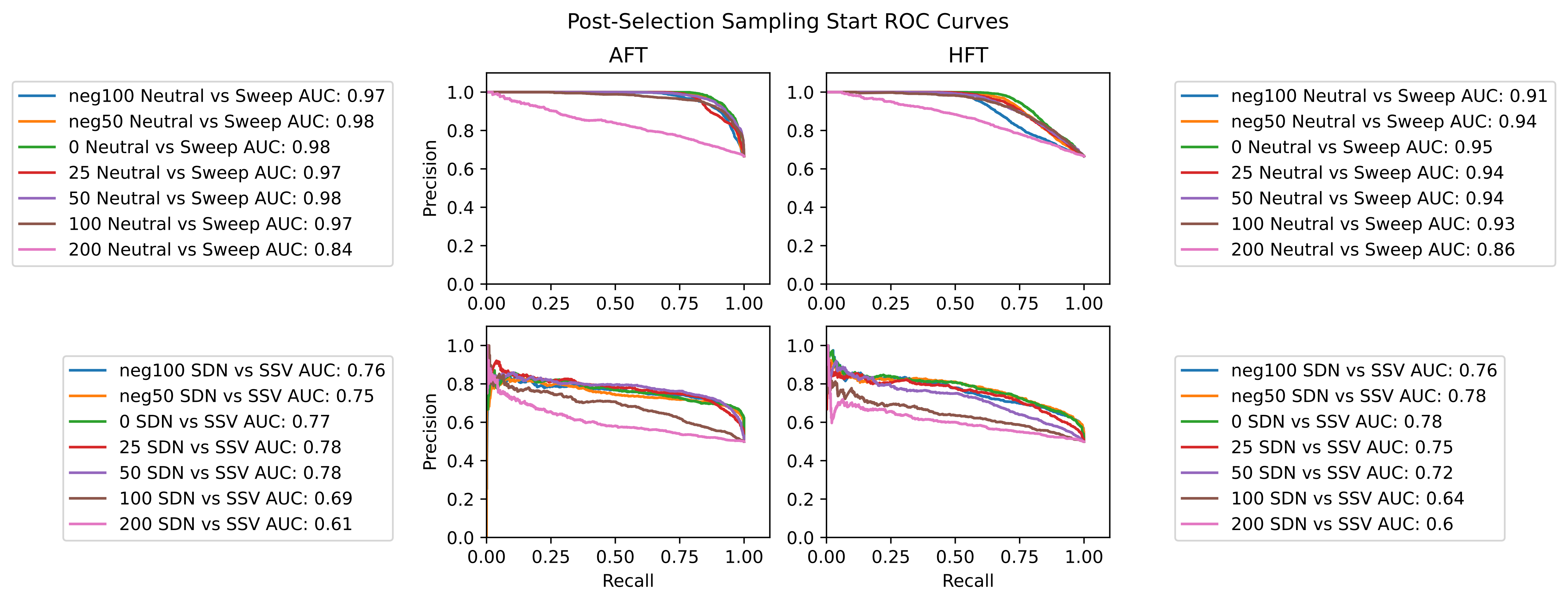

### S26_Sel_Timing_Reg.pdf

#### Regression - Selection Timing

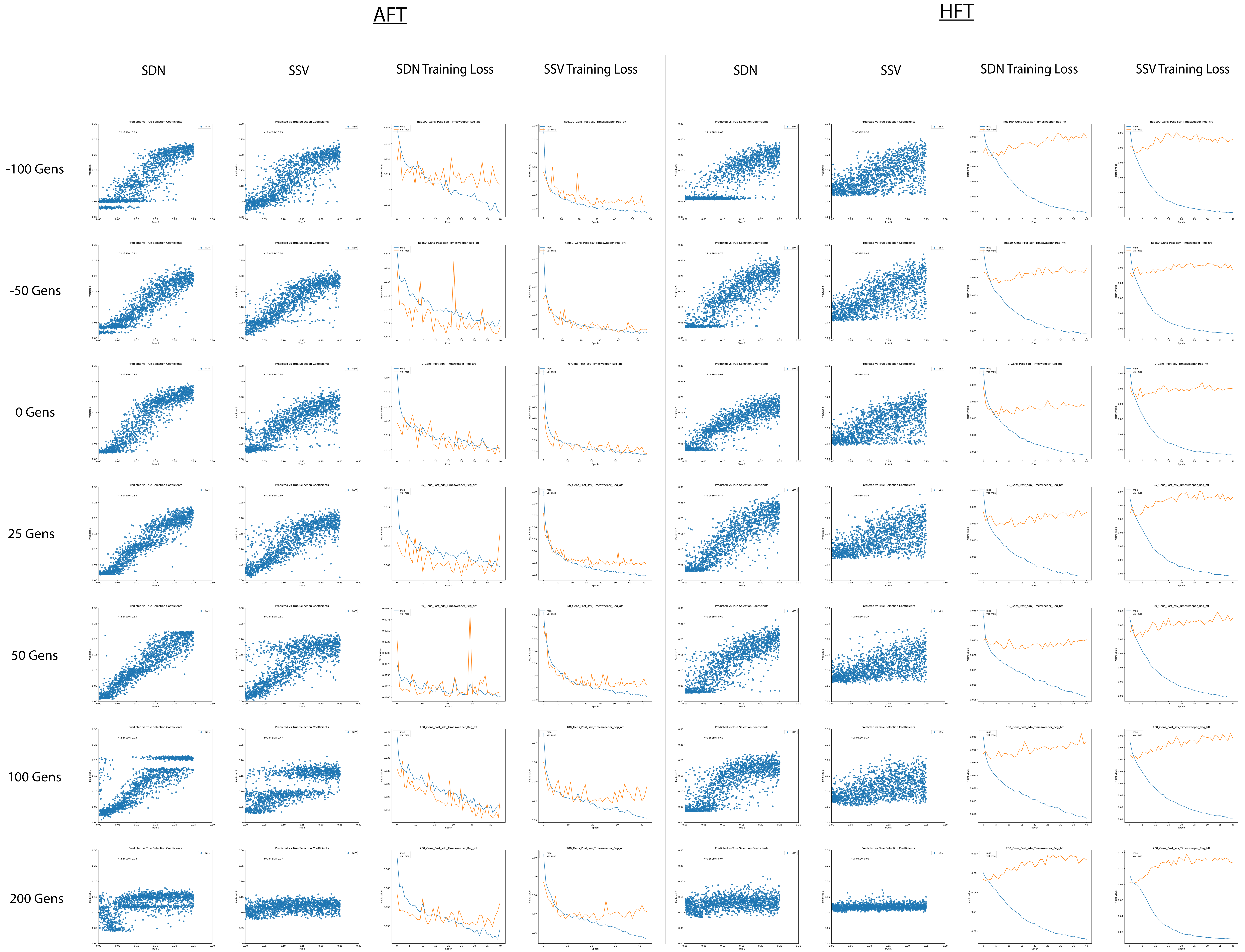

### S27_Num_Reps_Class.pdf

#### Classification - Number of Replicates

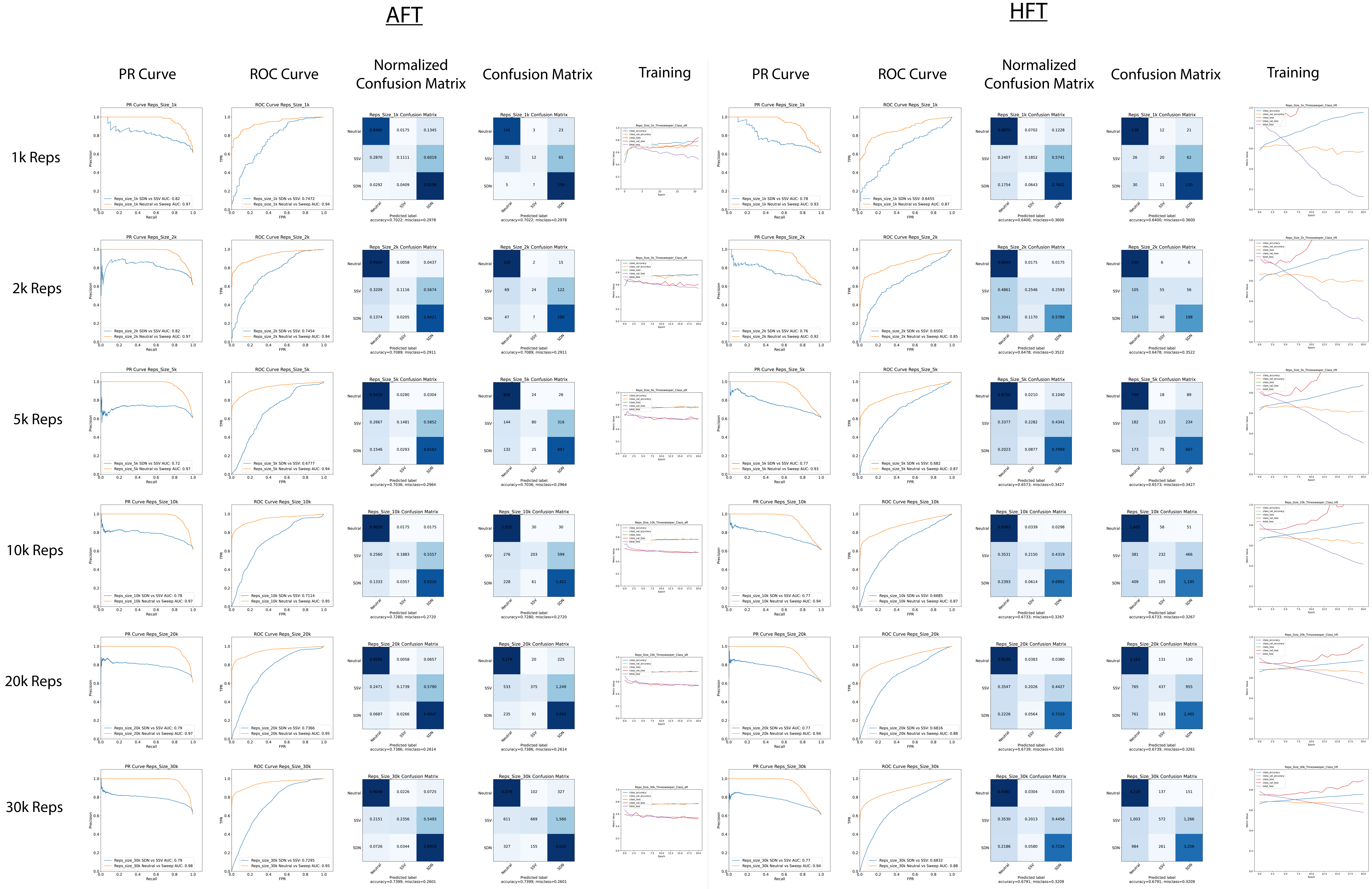

### S28_Num_Reps_ROC.pdf

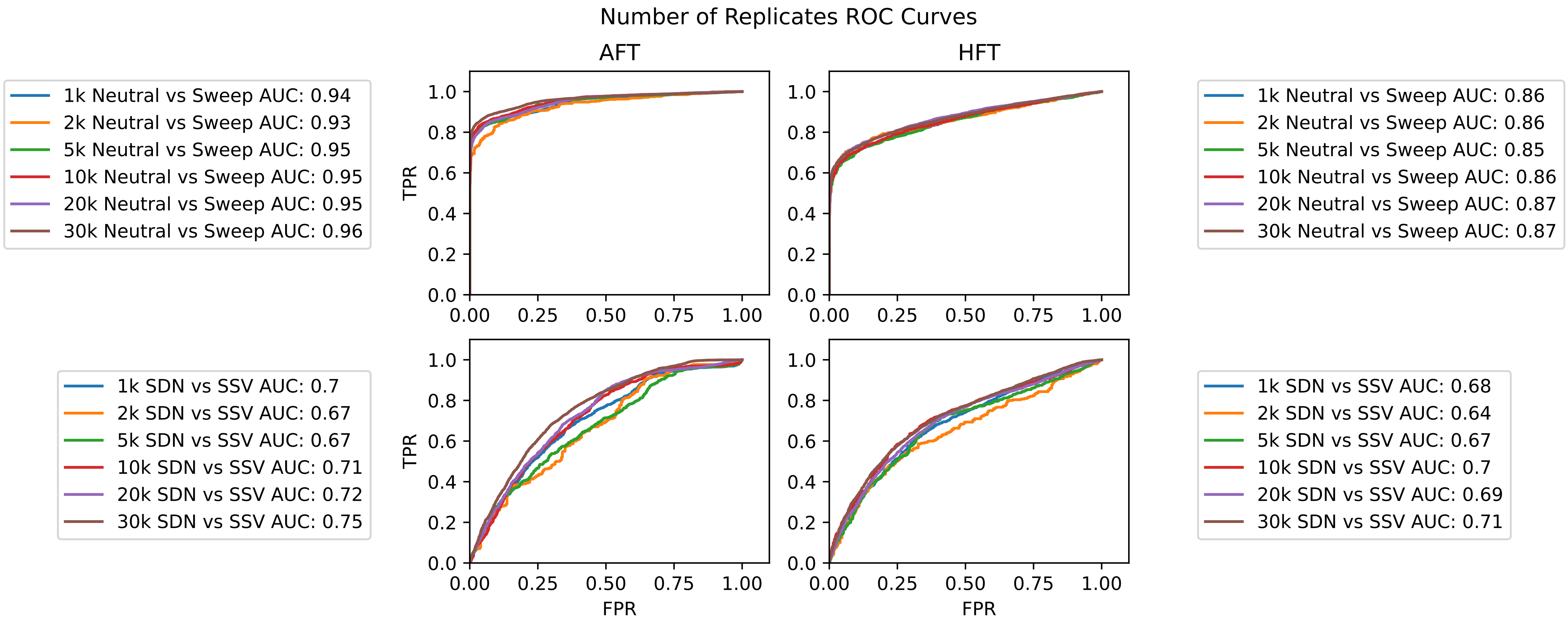

### S29_Num_Reps_PR.pdf

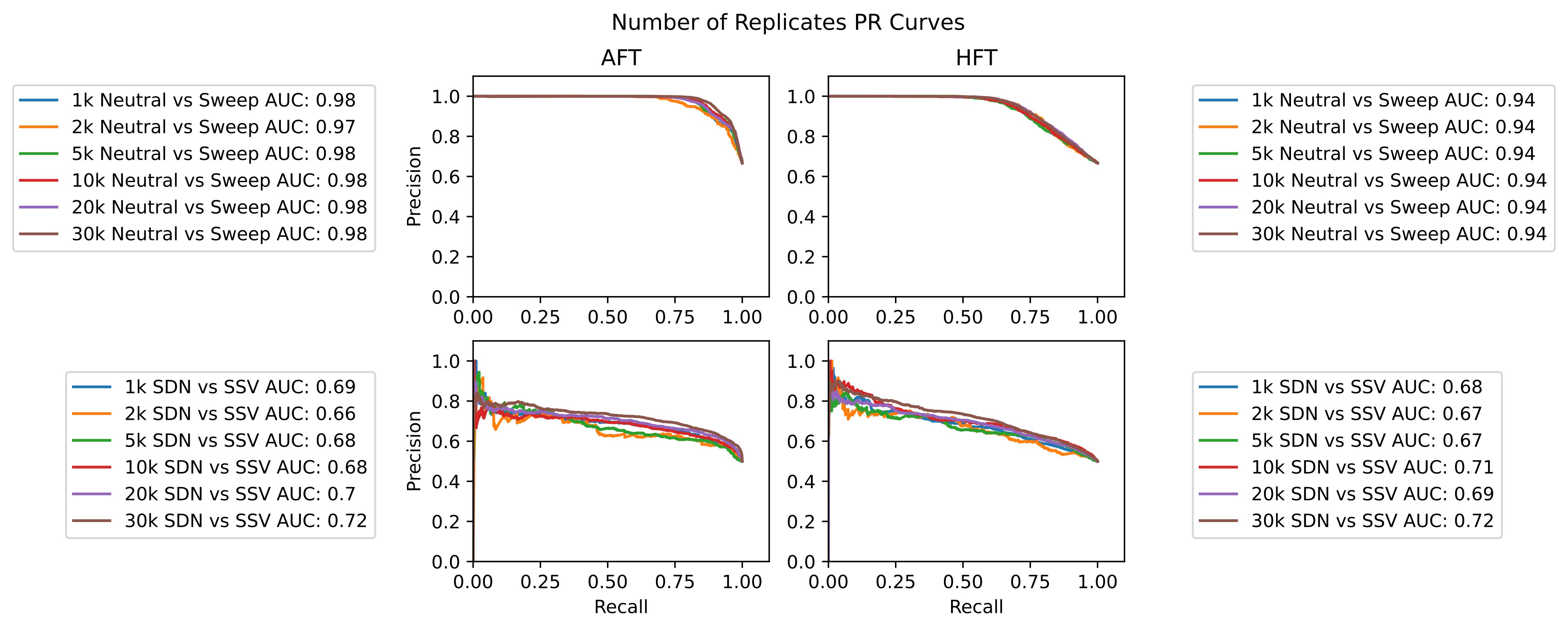

### S30_Num_Reps_Reg.pdf

#### Regression - Number Replicates

##### AFT

##### HFT

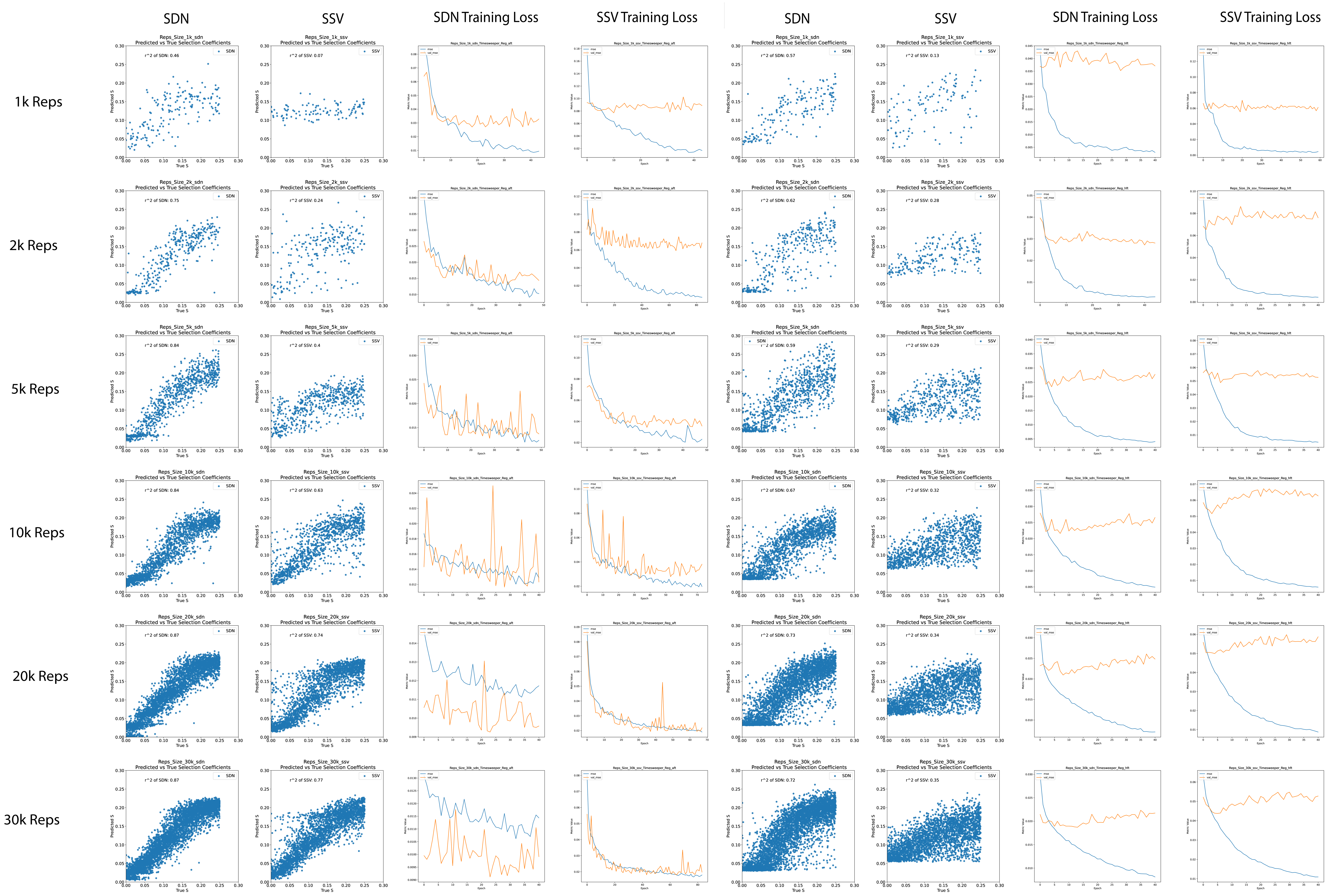

### S31_Zoomed_Spikes.pdf

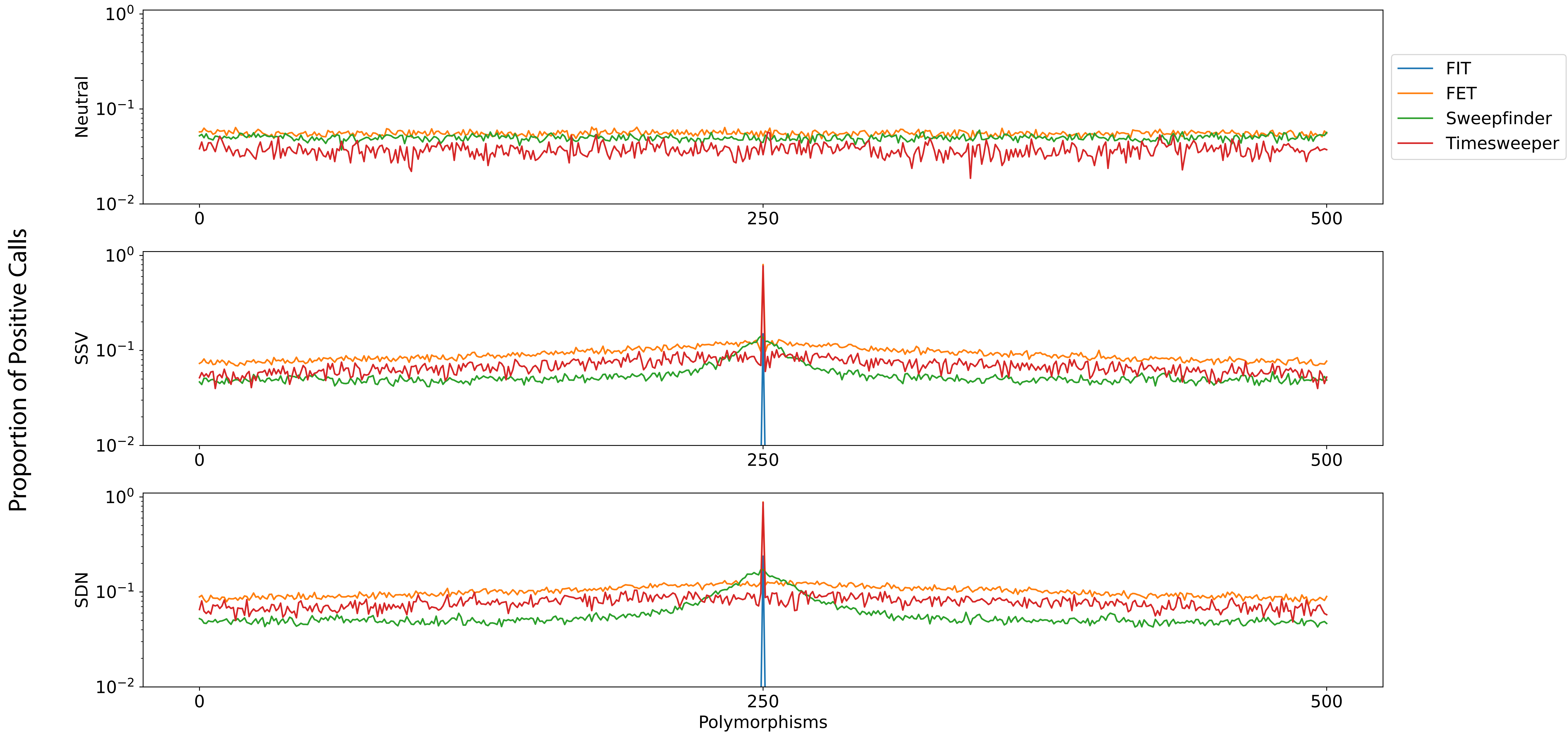

### S32_Misspec_Classification.pdf

#### Misspecification Classification ROC and PR Curves

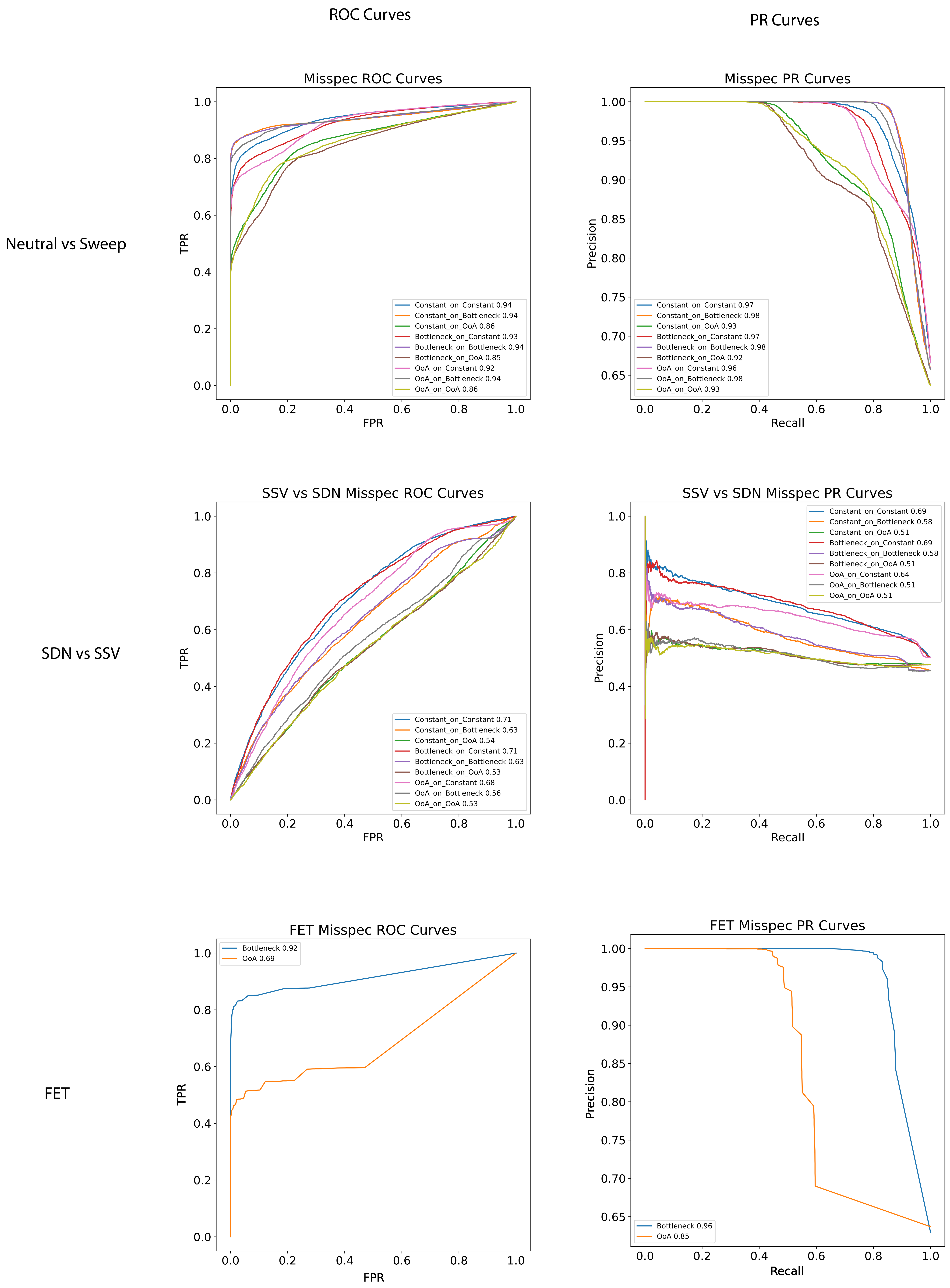

### S33_Misspec_Regression.pdf

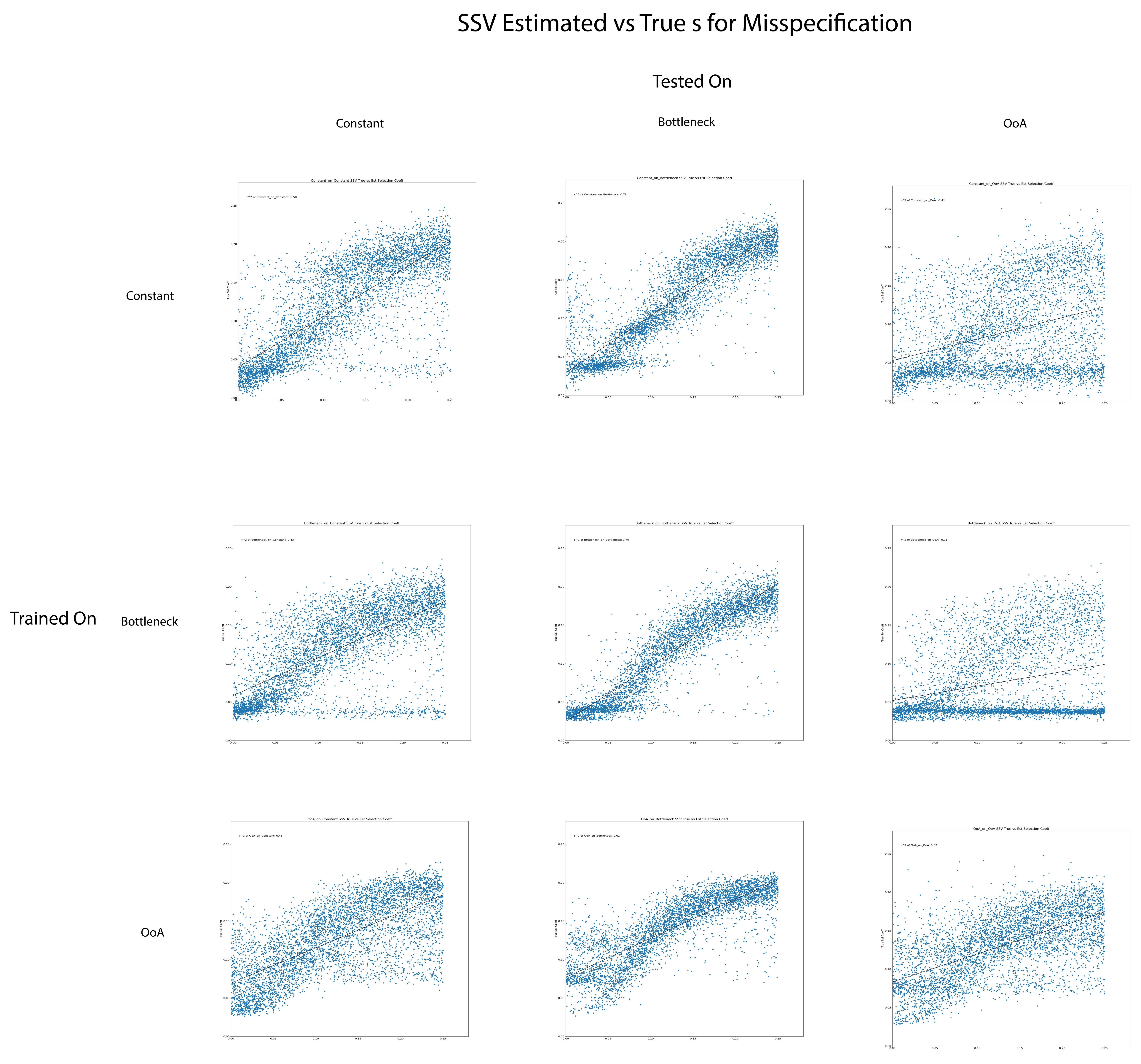

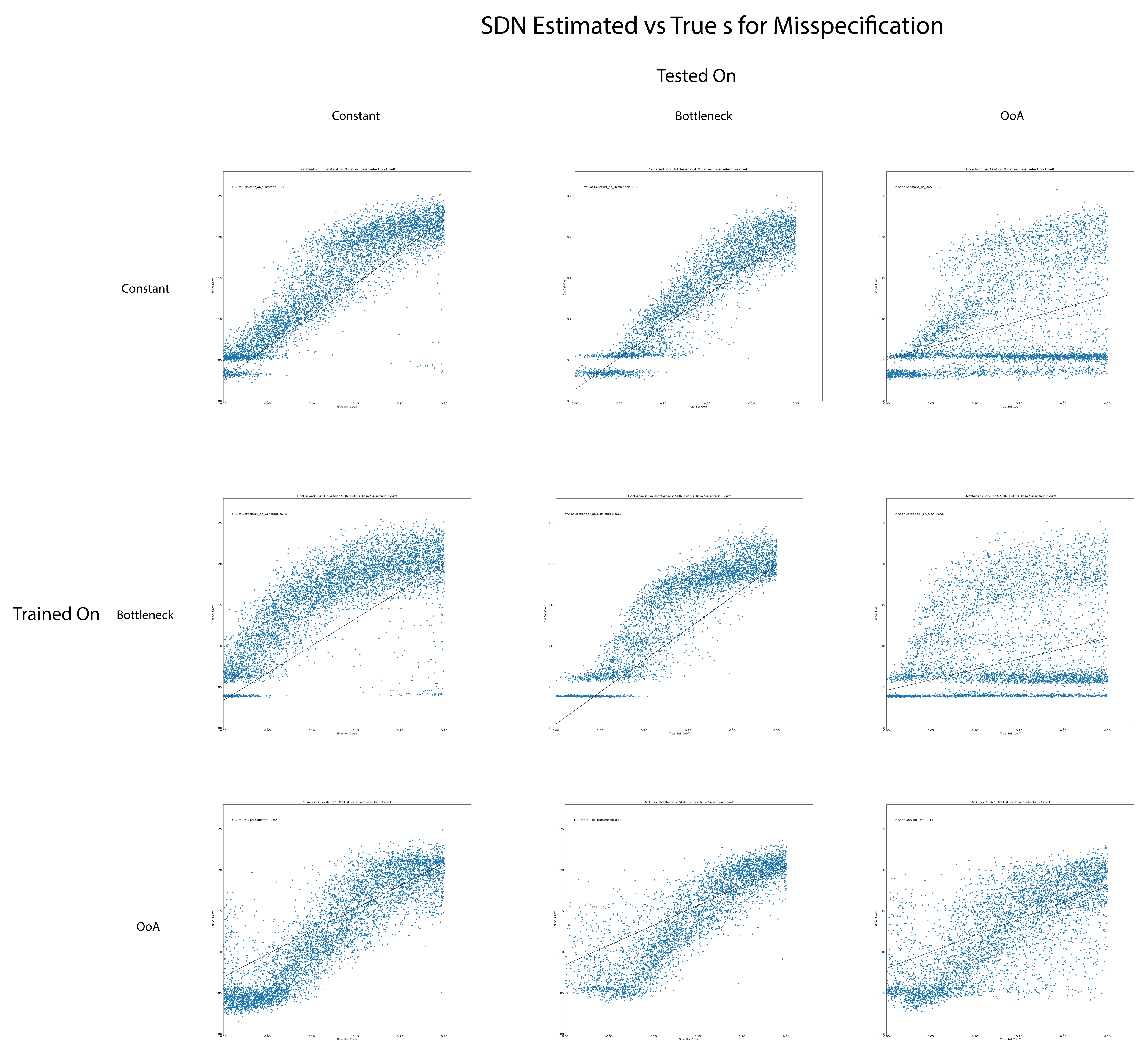

### S34-D_sim_Inputs.pdf

#### D. simulans AFT Input Data

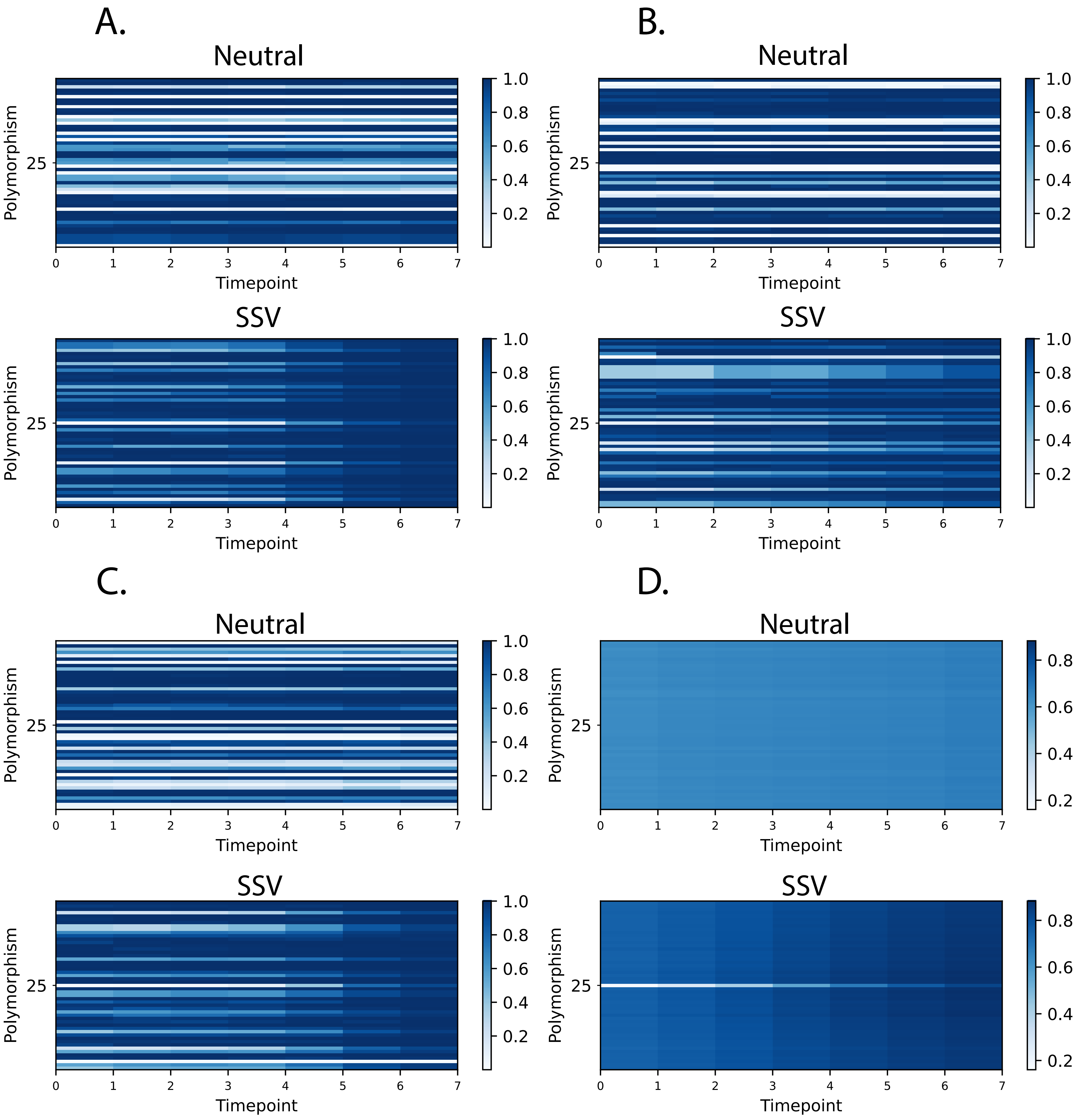

### S35_D_sim_Training_Metrics.pdf

#### D. simulans AFT Model Metrics

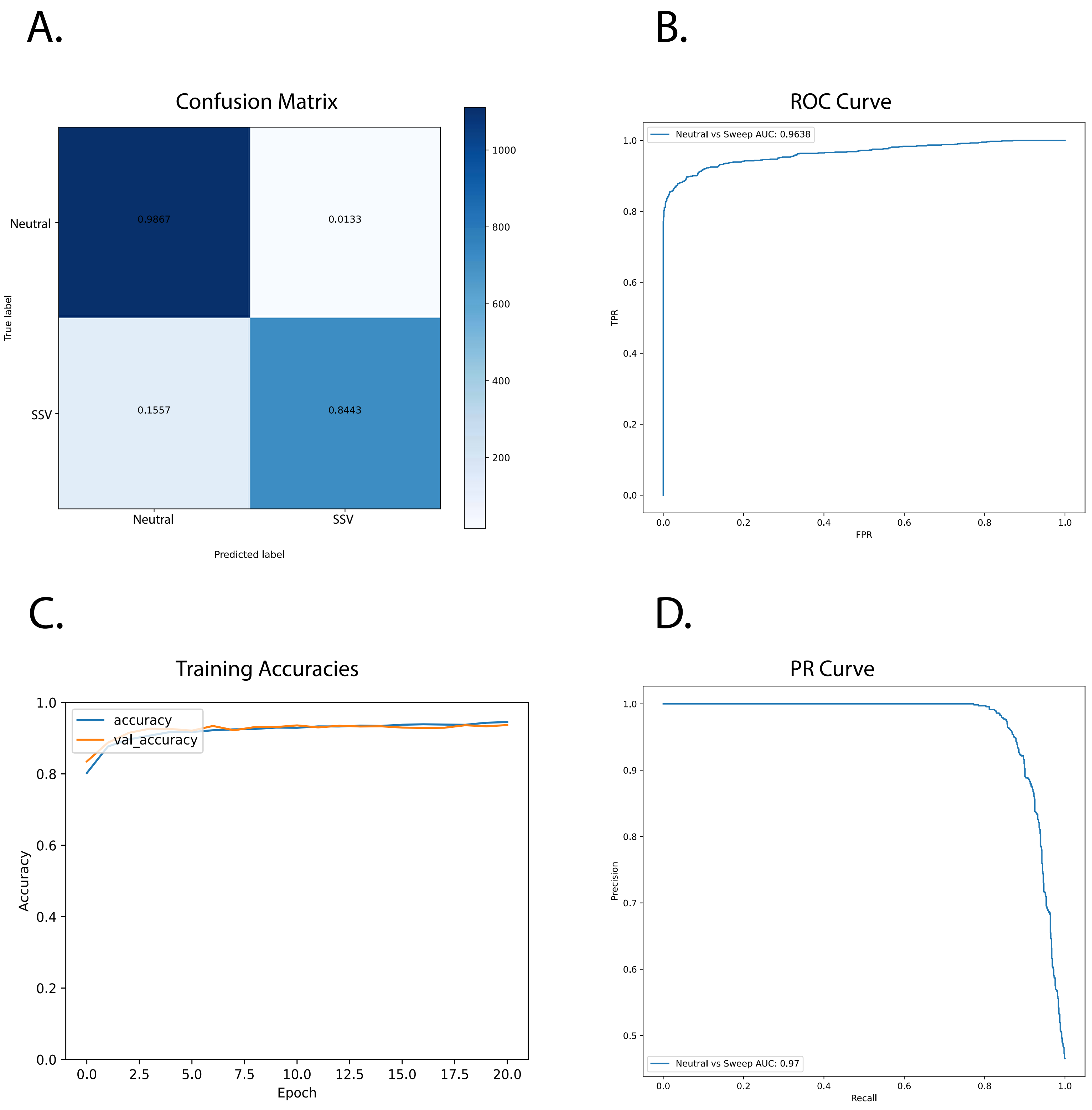

### S36_Rep_Hist.pdf

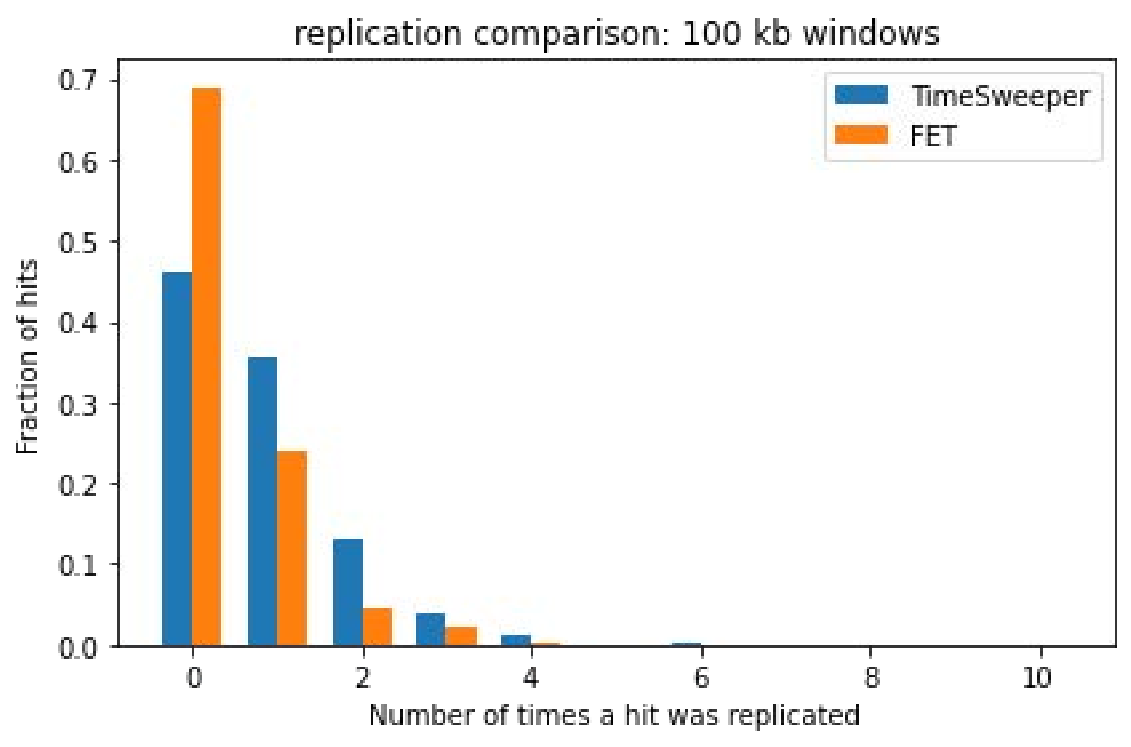
